# Physical interaction with Ephrin B1 promotes CXCR4 intracellular localization and oncogenic potential

**DOI:** 10.1101/2024.07.10.602459

**Authors:** Alessandro Rabbito, Omolade Otun, Amos Fumagalli, Martial Séveno, Sonya Galant, Manuel Counson, Thierry Durroux, Cherine Bechara, Martine J Smit, Sébastien Granier, Martyna Szpakowska, Andy Chevigné, Séverine Chaumont-Dubel, Philippe Marin

## Abstract

Chemokine receptor 4 (CXCR4) is a member of the chemokine receptor exclusively activated by the chemokine CXCL12. While CXCR4 regulates numerous physiological processes associated with cell migration and embryogenesis, its overexpression has been involved in various cancer types. Studies suggest that intracellular CXCR4 expression rather than CXCR4-operated signaling underlies its pro-tumorigenic functions. Given the role of GPCR interacting proteins in their trafficking and subcellular localization, we characterized the CXCR4 interactome using an affinity purification coupled to mass spectrometry (AP-MS) strategy. The most abundant protein identified in the CXCR4 interactome is Ephrin B1, a member of the Ephrin protein family that shares several functions with CXCR4, such as the regulation of cell migration and proliferation. Further studies showed that interaction between CXCR4 and Ephrin B1 is direct and enhanced by treating cell with CXCL12. They also indicated that Ephrin B1 prevents CXCR4 N-glycosylation, decreases CXCR4 cell surface expression and consistently inhibits CXCL12-induced CXCR4 coupling to G_αi1-3_ and the recruitment of β-arrestins 1 and 2. Conversely, Ephrin B1 signals to Erk1,2 through CXCR4 activation and mediates the decrease in Death Receptor 5 expression elicited by intracellular CXCR4. Collectively, these findings identify Ephrin B1 as a key mediator of CXCR4 tumorigenic signaling.

## Introduction

The C-X-C chemokine receptor type 4 (CXCR4) is a conventional chemokine receptor that unlike many chemokine receptors binds to a single chemokine, CXCL12, also referred to as SDF-1. CXCR4 is expressed in numerous cell types, including endothelial cells, lymphocytes, fibroblasts hematopoietic stem cells, neurons and glial cells where it plays a pivotal role in migration, proliferation and differentiation (Teixidó et al., 2018). CXCR4 is also involved in several pathological processes, including the Warts, Hypogammaglobulinemia, Immunodeficiency, and Myelokathexis (WHIM) syndrome and Human Immunodeficiency Virus-1 infection (Liu et al., 2016; Moore et al., 2004). It is overexpressed in numerous cancer types and has been involved cancer progression and metastasis (Müller et al., 2001).

CXCR4 is canonically coupled to Gα_i_ proteins and subsequently inhibits adenylyl cyclase. CXCR4 also activates the Erk1,2 pathway through several mechanisms involving either its phosphorylation by GRK3 and GRK6, and the recruitment of β-arrestin 1 or the activation of PI3Kγ heterodimers p110γ-p101 through Gβγ proteins (Busillo et al., 2010; Khater et al., 2021). Despite the unequivocal role of CXCR4-operated signaling, such as PI3K-Erk signaling, in cancer growth and metastasis, the limited therapeutic efficacy of CXCR4 antagonists, including the FDA-approved compound AMD3100, in cancer treatment, suggests that some tumorigenic effects under the control of the receptor are independent of the CXCL12-CXCR4 signaling axis (De Clercq, 2019). This conundrum has started to be solved with the recent demonstration that intracellular CXCR4 rather than CXCL12-CXCR4-operated signaling promotes cancer cell survival through the downregulation of the Death Receptor 5 (DR5), thereby rendering cancer cells resistant to chemotherapeutic drugs like paclitaxel (Nengroo et al., 2021). These findings suggest that future therapies targeting CXCR4 should not only consider CXCR4-associated signaling but also mechanisms underlying the targeting of CXCR4 to intracellular compartments.

There is accumulating evidence indicating that the trafficking and targeting of G protein-coupled receptors (GPCRs) to specific cellular compartments depends on their association with protein partners. For instance, while the association of CXCR4 with Filamin A stabilizes the receptor at the plasma membrane by blocking its endocytosis (Gómez-Moutón et al., 2015), its interaction with Reticulon 3 (RTN3) promotes its translocation to the cytoplasm. Even though RTN3 is known to enhance rather than inhibit TRAIL-induced apoptosis through the induction of DR5 expression in cancer cells (Lee et al., 2009), these findings highlight the potential of characterizing the CXCR4 interactome to identify novel mechanisms underlying its tumorigenic effects, an issue we addressed to thanks to a proteomic strategy combining affinity purification of receptor-interacting proteins and their identification by high-resolution mass spectrometry.

This interactomic screen identified Ephrin B1, a member of the single transmembrane domain Ephrin-B family overexpressed in numerous cancer types and involved in tumor progression and metastasis as well as resistance to chemotherapies (Pasquale, 2023). A previous study showed that Ephrin B1 modulate G protein signaling induced by CXCR4 activation via the recruitment of the regulator of G protein signaling 3 (RGS3) protein through its C-terminal PDZ domain, thus establishing functional interactions between CXCR4 and Ephrin B1 (Lu et al., 2001). In light of these findings, we further explored the impact of Ephrin B1 on CXCR4 post-translational modifications, cellular localization and signal transduction. We show that Ephrin B1 targets CXCR4 to intracellular compartments and mediates CXCR4-dependent decrease in Death Receptor 5 expression, suggesting that it might be a key player contributing to CXCR4 oncogenic potential.

## Materials and Methods

### Materials

#### Plasmids

The HA-CXCR4-pCDNA3.1 construct was obtained from cDNA.org (Bloomsburg University), and the FLAG-Ephrin B1-pcDNA3.1 plasmid was provided from the Montpellier Genomic Collections platform. The pcDNA3.1 plasmids encoding Venus-Tagged G protein subunit gamma 2 (Venus-γ2), FLAG-tagged G-protein subunit beta 2 (FLAG-β2), β-arrestin2-RLuc and Gαi3-RLuc were provided by Dr. D. Maurel (IGF, Montpellier, France). ACKR3-NeonGreen, CXCR4-NeonGreen, were described previously (Meyrath et al., 2021). HiBIT-CXCR4 plasmid was described in a previous study (Kumar et al., 2023). Ephrin B1-NeonGreen, Ephrin B1-Nluc, were cloned as previously described (Meyrath et al., 2021). Lyn-NeonGreen, ACKR3-Nluc, CXCR4-Nluc, LgBIT-β-arrestin1/2, CXCR4-smBIT in pcDNA3.1 were previously described (Luís et al., 2022; Palmer et al., 2022).

#### Reagents

CXCL12, purchased from R&D System (Ref 350-NS-CF) was dissolved in sterile PBS to a concentration of 10^-5^ M, aliquoted and stored at −80°C. AMD3100 purchased from Tocris (Ref 3299) was dissolved in DMSO to a final concentration of 20 mM, aliquoted and stored at −20°C. Gallein purchased from Tocris (Ref 3090) was dissolved in DMSO to a final concentration of 75 mM, aliquoted and stored at −20°C.

#### Antibodies

|  | Protein | Host<br>Species | Dilution &<br>Application | Supplier | REF |
| --- | --- | --- | --- | --- | --- |
| Primary<br>antibodies | CXCR4 | Rabbit | 1/1,000 WB | Abcam | AB124824 |
|  | Ephrin b1 | Rabbit | 1/2,000 WB | Biorbyt | orb395013 |
|  | HA | Rat | 1/1,000 WB | Roche | 11867423001 |
|  | CXCR4 pS<sup>324/325</sup> | Rabbit | 1/1,000 WB | 7-TM | 7TM0071A |
|  | CXCR4 pS<sup>330</sup> | Rabbit | 1/1,000 WB | 7-TM | 7TM0071C |
|  | CXCR4 pS<sup>346/347</sup> | Rabbit | 1/1,000 WB | 7-TM | 7TM0071B |
|  | DR5 | Rabbit | 1/1,000 WB | Cell signaling | 8074 |
|  | GAPDH | Rabbit | 1/5,000 WB | Cell signaling | 51332 |
|  | p44/42 MAPK | Rabbit | 1/1,000 WB | Cell signaling | 9102 |
|  | p-p44/42 MAPK | Rabbit | 1/1,000 WB | Cell signaling | 9101 |
|  | LAMP1 | Rabbit | 1/250 ICC | Cell signaling | 9091 |
|  | Calnexin | Rabbit | 1/250 ICC | Transduction laboratories | C45520 |
|  | GM130 | Mouse | 1/250 ICC | BD Transduction | 610823 |
|  | Label | Target<br>Species | Dilution &<br>Application | Supplier | REF |
| Secondary<br>antibodies | Alexa Fluor® 633 | Rabbit | 1/500 ICC | Invitrogen | 210071 |
|  | Alexa Fluor® 633 | Mouse | 1/500 ICC | Invitrogen | 210052 |
|  | HRP | Rat | 1/5,000 WB | Jackson ImmunoResearch | 112-035-003 |
|  | HRP | Rabbit | 1/5,000 WB | Cell signaling | 7074 |

### Cell cultures and transfection

HEK293T cells, purchased from ATCC (Anassas, VI, ATCC, CRL-1573), were grown in Dulbecco’s Modified Eagle’s Medium (DMEM, Thermo Fisher Scientific, Ref 419960) supplemented with 10% heat-inactivated fetal bovine serum (Thermo Fisher Scientific, Ref 10099-133), 1% penicillin/streptomycin, and maintained in humidified atmosphere containing 5% CO_2_ at 37°C. Cells were passed twice a week and used between passages 10 and 25. They were transfected using polyethylenimine (jetPEI^®^, Polyplus, Ref 101000053) and used 24/48 h after transfection.

MCF7 cells, provided by Dr. Stéphan Jalaguier (Institut de Recherche en Cancérologie de Montpellier), were cultured in /Dulbecco’s modified Eagle’s medium/F12 supplemented with 10% heat-inactivated bovine serum (Thermo Fisher Scientific, Ref 10099-133), 1% penicillin/streptomycin, and maintained in humidified atmosphere containing 5% CO_2_ at 37°C. Cells were passed twice a week and used between passages 10 and 25. They were transfected using Lipofectamine^TM^ 2000 (Thermo Fisher Scientific, Ref 11668019) and used 48 h after transfection.

### Co-immunoprecipitation

HEK293T cells transfected with empty vector or vectors expressing HA- or Flag-tagged proteins were lysed in ice-cold lysis buffer containing 1% n-Dodecyl-β-D-Maltopyranoside (DDM, Antrace, Ref D310), 20 mM Tris-HCl (pH 7.5), 100 mM NaCl, 2.5 mM CaCl_2_, phosphatase inhibitors (NaF, 10 mM; Na^+^-vanadate, 2 mM; Na+-pyrophosphate, 1 mM; and β-glycerophosphate, 50 mM) and the cOmplete Protease Inhibitor Cocktail (Merck, Ref 11836145001). Samples were maintained under gentle agitation for 1 h at 4 °C and centrifuged for 15 min at 15,000 × g to eliminate insoluble material. Soluble proteins were quantified by bicinchoninic acid assay (BCA, Merck, Refs B29643 and C2284) and equal protein amounts (5 or 0.5 mg for IP followed by mass spectrometry or Western blotting, respectively) were incubated with agarose-conjugated anti-HA antibody (Merck, Ref A2095) or with anti-FLAG M2 affinity gel (Sigma Aldrich, Ref A2220) overnight at 4 °C. Samples were then washed twice with an ice-cold solution of 0.5 M of NaCl and phosphatase inhibitors and three times with 0.25 M NaCl and phosphatase inhibitors. Immunoprecipitated proteins were then eluted in Laemmli sample buffer.

### Protein identification by mass spectrometry

Immunoprecipitated proteins from cells transfected with an empty vector (Mock) or a vector HA-tagged CXCR4 were separated by SDS-PAGE and stained with Protein Staining Solution (Euromedex, Ref 10-0911). Gel lanes were cut into seven gel pieces and destained with 50 mM TriEthylAmmonium BiCarbonate (TEABC, Merck, Ref T7408) followed by three washes in 100% acetonitrile. After reduction (with 10 mM dithiothreitol in 50 mM TEABC at 60 °C for 30 min) and alkylation (with 55 mM iodoacetamide TEABC at room temperature for 60 min), proteins were digested in-gel using trypsin (500 ng/band, Gold, Promega, Ref V5280). Digest products were dehydrated in a vacuum centrifuge and reduced to 3 μL. The generated peptides were analyzed online by nano-flow liquid chromatography coupled to tandem-mass spectrometry (nanoLCMS/MS) using an Orbitrap Elite mass spectrometer (Thermo Fisher Scientific, Waltham USA) coupled to an Ultimate 3000 HPLC (Thermo Fisher Scientific). Desalting and pre-concentration of samples were performed on-line on a Pepmap® pre-column (0.3 mm × 10 mm, Dionex). A gradient consisting of 0–40% B for 60 min and 80% B for 15 min (A = 0.1% formic acid, 2% acetonitrile in water; B = 0.1% formic acid in acetonitrile) at 300 nL/min was used to elute peptides from the capillary reverse-phase column (0.075 mm × 150 mm, Acclaim Pepmap 100® C18, Thermo Fisher Scientific). Eluted peptides were electrosprayed online at a voltage of 1.8 kV into the Orbitrap Elite mass spectrometer. A cycle of one full-scan mass spectrum (MS1, 400–2,000 m/z) at a resolution of 120,000 (at 400 m/z), followed by 20 data-dependent tandem-mass (MS2) spectra was repeated continuously throughout the nanoLC separation. All MS2 spectra were recorded using normalized collision energy (33%, activation Q 0.25 and activation time 10 ms) with an isolation window of 2 m/z. Data were acquired using the Xcalibur software (v 2.2). For all full scan measurements with the Orbitrap detector a lock-mass ion from ambient air (m/z 445.120024) was used as an internal calibrant63. Mass spectra were processed using the MaxQuant software package (v 1.5.5.1) and MS2 using the Andromeda search engine “[http://coxdocs.org/doku.php?id=maxquant:andromeda:start]” against the UniProtKB Reference proteome UP000005640 database for Homo sapiens (release 2017_10) and the contaminant database in MaxQuant. The following parameters were used: enzyme specificity set as Trypsin/P with a maximum of two missed cleavages, Oxidation (M) and Phosphorylation (STY) set as variable modifications and carbamidomethyl (C) as fixed modification, and a mass tolerance of 0.5 Da for-fragment ions. The maximum false peptide and protein discovery rate was specified as 0.01. Seven amino acids were required as minimum peptide length. When precursor peptides were present in MS1 spectra but not selected for fragmentation and identification by MS2 in given runs, peptide identification were based on accurate mass and retention times across LC-MS runs using the matching between runs tool in MaxQuant. Only proteins identified in all three biological replicates in at least one group were considered for further analysis. Relative protein quantification in IP from CXCR4-expressing cells and Mock cells was performed using the label-free quantification (LFQ) algorithm “[https://maxquant.net/maxquant/]”. For statistical analysis, missing values were defined using the imputation tool of the Perseus software (v. 1.5.6.072) “[https://maxquant.net/perseus/]”. The mass spectrometry data have been deposited to the ProteomeXchange Consortium via the PRIDE partner repository with the dataset identifier PXDXXX, [https://www.ebi.ac.uk/pride/archive/projects/ <u>PXDXXX</u>].

### Western blotting

Proteins were separated by SDS-PAGE using 10% polyacrylamide gels and transferred electrophoretically to nitrocellulose membranes (Bio-Rad, Ref 1704271). Membranes were incubated in blocking buffer (Tris-HCl, 50 mM, pH 7.5; NaCl, 200 mM; Tween-20, 0.1% and skimmed dried milk, 5%) for 1 h at room temperature and overnight with primary antibodies in incubating buffer (Tris-HCl, 50 mM, pH 7.5; NaCl, 200 mM; Tween-20, 0.1% and Bovine Serum Albumin (Merck, Ref A2153), 3%) at 4 °C. Then membranes were immunoblotted with either or anti-rat (Jackson ImmunoResearch, Ref 112-035-003) or anti-rabbit (Cell Signaling, Ref 7074) horseradish peroxidase (HRP)-conjugated secondary antibodies (1/5,000) in blocking buffer for 1 h at room temperature. Immunoreactivity was detected with an enhanced chemiluminescence method (Western lightning® Plus-ECL, Perkin Elmer, Ref NEL103E001EA) on a ChemiDoc™ Touch Imaging System (Bio-Rad). Quantification was performed using the Image Lab software (Bio-Rad).

### Production and purification of CXCR4 and Ephrin B1

For production in insect cells, the full-length genes encoding human CXCR4 or Ephrin B1 was subcloned into pFastBac1 to enable infection of sf9 insect cells. The CXCR4 construct bore a hemagglutinin signal peptide followed by a Flag-tag preceding the receptor sequence. Mutations N11Q, S18A and N33Q residues were introduced to avoid N- or O-glycosylation for improved sample heterogeneity. The Ephrin B1 sequence was followed by a C-terminal 3C human rhinovirus 3C (HRV3C) protease cleavage site and a twin-Strep-tag (Strep-Ephrin B1).

Flag-CXCR4 and Strep-Ephrin B1 were expressed in Sf9 cells using the pFastBac baculovirus system (Thermo Fisher Scientific). Cells were grown in suspension in EX-CELL 420 medium (Sigma-Aldrich) and infected at a density of 4 × 10^6^ cells/ml with the recombinant baculovirus. Flasks were shaken for 48 h at 28 °C, subsequently harvested by centrifugation (3,000 × *g*, 20 min) and stored at −80°C until usage. Cell pellets were first thawed and lysed by osmotic shock in a lysis buffer containing 10 mM Tris-HCl (pH 7.4), 1 mM EDTA, 2 mg/ml iodoacetamide, and protease inhibitors: 50 μg/ml Leupeptin (Euromedex), 0.1 mg/ml Bensamidine (Sigma-Aldrich) and 0.1 mg/ml Phenylmethylsulfonyl fluoride (PMSF; Euromedex). Lysates were centrifuged (38,400 × *g*, 10 min) and the resulting pellet was solubilized in buffer containing 50 mM Tris (pH 7.5), 150 mM NaCl, 2 mg/ml iodoacetamide, 0.5% (w/v) dodecyl maltoside (DDM, Anatrace), Cholesteryl hemisuccinate (CHS, 0.1% w/v) and protease inhibitors (50 μg/ml Leupeptin, 0.1 mg/ml Bensamidine and 0.1 mg/ml PMSF) using a Dounce homogenizer. Resulting homogenates were subsequently stirred for 1 h at 4 °C and centrifuged (38,400 × *g*, 30 min). CXCR4 supernatant was loaded onto M2 anti-Flag affinity resin (Sigma-Aldrich Ref A2220) using gravity flow. Ephrin B1 supernatant was supplemented with BioLock (0.75 ml/L) and incubated with Streptactin resin (IBA-Lifesciences, Ref 2-1206-025) for 1 h. Resins were each washed with 10 column volumes (CV) of wash buffer containing 50 mM Tris-HCl pH 7.4, 150 mM NaCl, 0.1% (w/v) DDM and 0.02% (w/v) CHS. The resin was finally washed with a last wash buffer containing 50 mM Tris-HCl (pH 7.4), 150 mM NaCl, DDM 0.02% (w/v), CHS 0.002% (w/v) and eluted in last wash buffer supplemented with 0.4 mg/ml Flag peptide for elution of CXCR4 or 2.5mM desthiobiotin for elution of Ephrin B1. Each eluate was concentrated using a 50 kDa molecular weight cutoff (MWCO) concentrator (Millipore). CXCR4 was then purified by size exclusion chromatography (SEC) using a Superdex 200 Increase (10/300 GL column) connected to an ÄKTA purifier system (GE Healthcare) and eluted in SEC buffer (50 mM Tris-HCl (pH 7.4), 150 mM NaCl, 0.02% (w/v) DDM, 0.002% (w/v) CHS). Buffers used for CXCR4 purification were supplemented with the CXCR4 antagonist IT1t (1 µM in the lysis and solubilization buffers, 0.1 µM in the wash and elution buffers, Bio-Techne) to stabilize the receptor.

### Co-immunoprecipitation of purified CXCR4 and Ephrin B1

Complexes of the two purified proteins were formed by mixing them at 2:1 ratio and incubating samples at 4°C overnight. The samples were then loaded onto M2 anti-Flag affinity resin, washed with 6CV of SEC buffer and eluted with SEC buffer supplemented with 0.4 mg/ml Flag peptide. The resulting eluate was loaded onto a column containing Streptactin resin, washed with 6CV of SEC buffer and eluted with SEC buffer supplemented with 2.5 mM Desthiobiotin.

### BRET

HEK293T cells were co-transfected with the indicated constructs and seeded (25,000 cells/well) in white polyornithine-coated-96-well plates (SPL Life Sciences). Twenty-four hours after transfection cells were washed with PBS. Coelenterazine (Nanolight Technology, Ref 50909-86-9) was added at a final concentration of 5 μM for 10 min at 37°C. Cells were then exposed to vehicle or CXCL12 and luminescence was measured using a Mithras LB 940 plate reader (Berthold Biotechnologies) that allows the sequential integration of light signals detected with two filter settings (Rluc/NLuc filter, 485 ± 20 nm; and YFP filter, 530 ± 25 nm). Data were collected using the MicroWin2000 software (Berthold Biotechnologies).

### Surface Nanoluciferase Complementation (HiBiT)

Receptor cellular distribution in basal conditions and upon ligand stimulation was monitored by using a nanoluciferase complementation assay based on the NanoGlo HiBiT extracellular detection system (Promega). HEK293T cells were cotransfected with pHiBiT vector encoding CXCR4 N-terminally fused to HiBiT and a vector encoding Ephrin B1. Forty-eight hours after transfection, cells were distributed in white 96-well plates (50,000 cells per well) and stimulated with CXCL12 (10 or 50 nM) for 30 min at 37°C. After addition of Nano-Glo® HiBiT reagent, containing soluble LgBiT protein, in ratio of 1:100 of the final volume, luminescence was recorded for 30 min with a GloMax plate reader (Promega). In unstimulated conditions, surface and total receptor expression were determined using the Nano-Glo HiBiT extracellular detection system (Promega) and Nano-Glo HiBiT lytic detection system (Promega), respectively.

### Nanoluciferase Complementation (nanoBiT)

β-arrestin-1 and β-arrestin-2 recruitment to CXCR4, was monitored using a nanoluciferase complementation-based assay (NanoBiT, Promega). HEK293T cells were co-transfected with vectors encoding human β-arrestins N-terminally fused with LgBiT CXCR4 C-terminally fused with SmBiT in the absence or presence of a vector encoding Ephrin B1. Twenty-four hours after transfection, cells were harvested, incubated for 20 min at 37°C with coelenterazine H in Opti-MEM, and distributed into white 96-well plates (150,000 cells/well). CXCL12 was then added at the indicated concentration and the luminescence generated upon nanoluciferase complementation was measured with a GloMax plate reader (Promega).

### Immunocytochemistry and confocal microscopy

HEK293T cells grown on glass coverslips were fixed with a 4% solution of PFA in PBS for 10 min. Excess of PFA was quenched by washing cells in a 0.1 M solution of glycine in PBS for 10 min. Cells were permeabilized with a PBS solution containing 0.5% heat-inactivated bovine serum and 0.1% Triton X-100 for 15 min. Cells were washed three times and incubated with primary antibodies (anti-LAMP1 (rabbit or mouse) or anti-GM130 (mouse) or anti-Calnexin (rabbit)) in PBS containing 0.5% heat-inactivated bovine serum and 0.1% Triton X-100. After three washes in PBS, cells were incubated for 1 h at room temperature in PBS containing 0.5% bovine serum, 0.1% Triton X-100, the appropriate secondary antibody (Alexa Fluor® 633-conjugated anti-mouse antibody (1/250, Invitrogen, Ref 210052), Alexa Fluor® 633-conjugated anti-rabbit antibody (1/250, Invitrogen Ref 210071)) and DAPI (1µg/ml). Cells were then washed three times with PBS and coverslips were mounted on Superfrost ultra plus glass slides using fluorescent mounting medium. Pictures were acquired with a LSM980 confocal microscope (Zeiss) equipped with a 40X oil-immersed lens, with a 2791 × 2791 resolution, and 1 µm between focus points.

### Statistics

Statistical analysis of the CXCR4 interactome was performed using the Perseus software (v 1.5.6.072). Proteins were considered statistically significant using a t-test by setting the randomization number at 250, the False Discovery Rate at 0.01 and the S0 at 0.1. All other statistical analyses were performed using Prism (v.8.0, GraphPad Software Inc) and the statistical tests used are indicated in each legend. Dose-response curves (**Figures 3C** and **4D**) were fitted by the log (agonist) vs. response (four parameters) non-linear regression using Prism. Saturation BRET experiments were analyzed by using Prism, and to obtain the curve fit, the one-site specific binding model was used. All data are presented as means ± SEM. Significance levels were defined as p < 0.05 (*), p < 0.01 (**), p < 0.001 (***), and p < 0.0001 (****).

## Results

### Characterization of the CXCR4 interactome in HEK-293T cells reveals a physical interaction between CXCR4 and Ephrin B1

We analyzed the CXCR4 interactome in human embryonic kidney (HEK-293T) cells that endogenously express CXCR4 and where CXCR4-dependent signaling has been extensively investigated (Busillo et al., 2010; Khater et al., 2021; Mueller et al., 2013). Due to the lack of a CXCR4 antibody providing receptor immunoprecipitation yields compatible with mass spectrometry analysis, we expressed hemagglutinin (HA)-tagged CXCR4 in the cells and CXCR4-interacting proteins were immunoprecipitated using an anti-HA monoclonal antibody immobilized onto agarose beads. Control immunoprecipitations were performed using cells transfected with an empty plasmid (Mock condition). Systematic analysis by LC-MS/MS of proteins in immunoprecipitates from both conditions in three biological replicates identified 1,203 proteins. Label-free quantification of their relative abundance in both conditions showed that 19 of them exhibited significant enrichment in immunoprecipitates from CXCR4-expressing cells, compared with immunoprecipitates from control cells (**Figure 1A and Supplementary Table 1**). As expected, CXCR4 (bait protein) was the most enriched one (log_2_LFQ_CXCR4_/LFQ_Mock_ = 8.13), while the second ranked protein was Ephrin B1 (log_2_LFQ_CXCR4_/LFQ_Mock_ = 7.29, -log_10_ p-value = 4.96, **Supplementary Table 1 and Figure 1A**). Ephrin B1 is a member of the Ephrin-B family comprising three proteins (Ephrin B1-3) that act as ligands of the tyrosine kinase Eph receptors involved in numerous biological processes, including axon guidance and cell migration during neurodevelopment, and the proliferation of various cancer cell types. Notably, Ephrin B1 was the only member of the Ephrin ligand families identified in the CXCR4 interactome.

**Figure 1:**
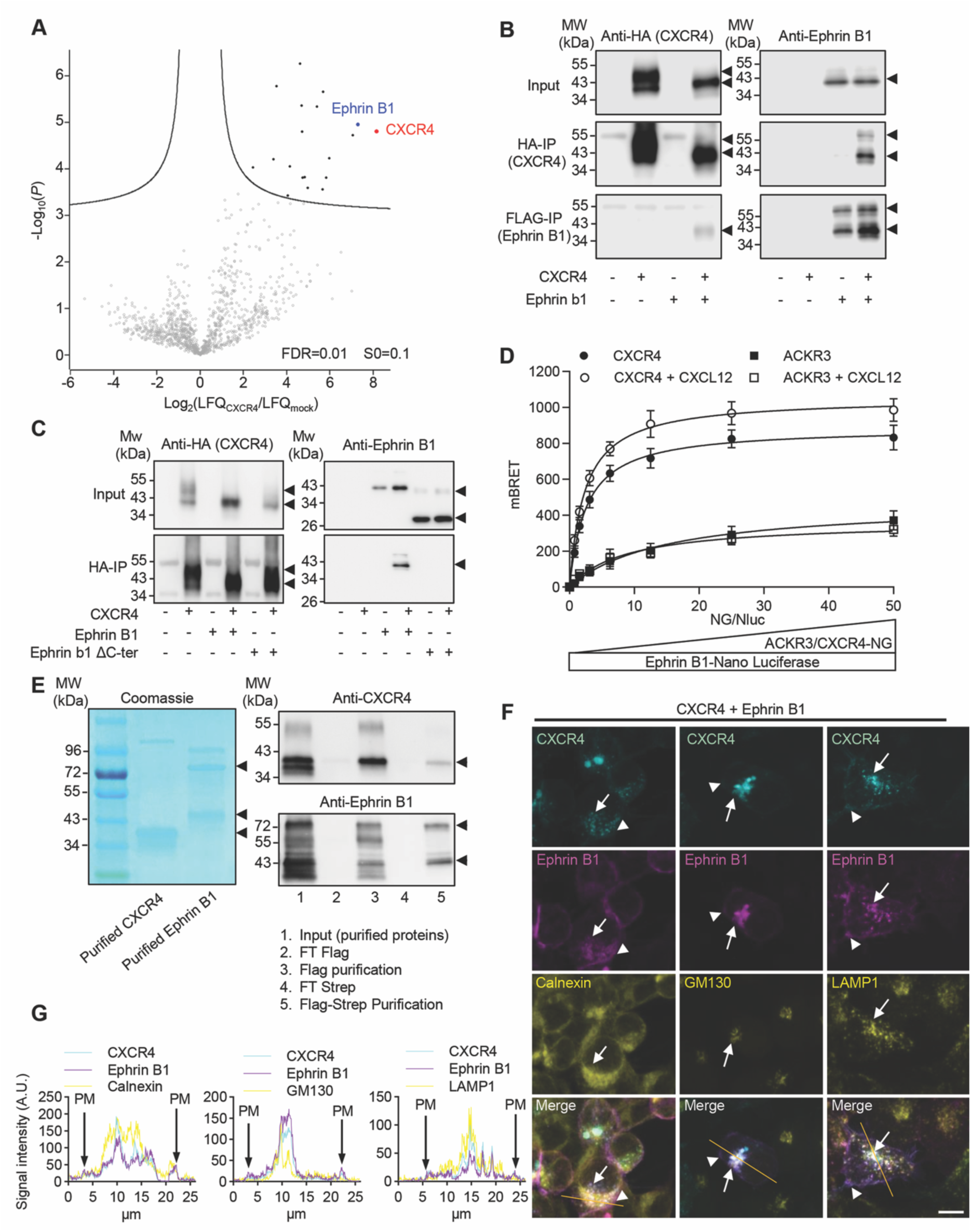
Physical interaction of Ephrin B1 with CXCR4. **A** Volcano plot representing proteins identified by nanoLC-MS/MS in immunoprecipitations from HEK293T cells transfected with HA-CXCR4 or empty plasmid. **B** Representative Western blot of HA and Flag immunoprecipitations performed from HEK293T cells expressing or not HA-CXCR4 or FLAG-Ephrin B1 or both proteins. **C** Representative Western blot of HA-immunoprecipitations performed from HEK293T cells expressing or not HA-CXCR4, or FLAG-Ephrin B1 or Ephrin B1 deleted of its C-terminal domain (Ephrin B1 ΔC-ter) alone or in combination. In **B** and **C**, the illustrated blots are representative of three independent experiments performed from different sets of cultured cells. **D** Normalized BRET values measured in HEK293T cells co-expressing the indicated proteins, BRET signal variations were fitted with the one-site total binding equation and background constraint to constant value of 0 using Prism. **E** Left panel: Coomassie blue staining of purified FLAG-CXCR4 and STREP-Ephrin B1 produced in the SF9 cell line. Right panels: Western blots of sequential Flag/Strep chromatography purification. The illustrated data are representative of three independent experiments. FT: flow-through. **F** Stacked images of CXCR4-YFP (cyan) and Ephrin B1-RFP (magenta) fluorescent signals in different cellular compartments in HEK293T cells co-expressing CXCR4-YFP and Ephrin B1-RFP. The anti-calnexin immunostaining was used to label the endoplasmic reticulum, the anti-GM130 immunostaining the cis-Golgi, and anti-LAMP1 immunostaining lysosomes (yellow). The illustrated fields are representative of three independent experiments performed from different sets of cultured cells. Scale bar: 10 μm. **G** Line graphs generated in Image J using the lines represented on the merge images highlighting the colocalization of CXCR4 and Ephrin B1 in the three compartments. PM, plasma membrane signal.

Considering the strong enrichment of Ephrin B1 in the CXCR4 interactome, their role in common cellular and pathological processes and the existence of functional interactions between both proteins (Holm et al., 2007; Konoplev et al., 2007; McKinney et al., 2015), we then focused on CXCR4-Ephrin B1 interaction. Co-immunoprecipitation followed by Western blotting from cells co-expressing HA-CXCR4 and Flag-Ephrin B1 confirmed that Ephrin B1 co-immunoprecipitated with CXCR4 and vice versa (**Figure 1B**). The C-terminal domain of GPCRs has been identified as a major site involved in their interaction with their protein partners (Bockaert et al., 2003). Nevertheless, the co-immunoprecipitation of Ephrin B1 with CXCR4 was not affected by the deletion of its 15 C-terminal residues (major mutation responsible for the WHIM syndrome, CXCR4-WHIM, (Lagane et al., 2008)) nor of its entire C-terminal domain (CXCR4ΔCter, **Supplementary Figure 1**), indicating that the CXCR4 C-terminal domain is not involved in the recruitment of Ephrin B1. In contrast, the C-terminal domain of Ephrin B1 seems to be essential to the formation of the CXCR4-Ephrin B1 complex, as its deletion abolished the co-immunoprecipitation of Ephrin B1 with CXCR4 (**Figure 1C**).

We next analyzed CXCR4-Ephrin B1 interaction in living HEK-293T cells using bioluminescence resonance energy transfer (BRET). Under conditions of constant Ephrin B1-NLuc expression, the BRET signal increased hyperbolically as a function of the CXCR4-Neon Green expression level (**Figure 1D**). Further, the BRET signal was significantly increased upon CXCR4 activation by CXCL12 (50 nM, 5 min), indicating that the recruitment of Ephrin B1 is promoted by agonist stimulation of CXCR4. Corroborating previous interactomics studies that did not identify Ephrin B1 as a protein partner of ACKR3 (Fumagalli et al., 2020), an atypical chemokine receptor known to form heteromers and to be functionally linked with CXCR4 (Levoye et al., 2009), the BRET signal was much lower in cells co-expressing Ephrin B1-NLuc and increasing amounts of ACKR3-Neon Green than that measured in cells co-expressing Ephrin B1 and CXCR4 (**Figure 1D**). Collectively, these observations indicate a close and specific interaction between CXCR4 and Ephrin B1. To establish whether the interaction between both proteins is direct, purified recombinant Flag-CXCR4 and Strep-Ephrin B1 were obtained from Sf9 cells, mixed at 2:1 protein ratio and subjected to sequential affinity purification on M2 anti-Flag affinity resin and, after elution of the retained material, on Streptactin resin. We found that Strep-Ephrin B1 was co-purified with Flag-CXCR4 on Flag resin while Flag-CXCR4 was co-copurified with Strep-Ephrin B1 on Streptactin resin, indicative of a direct physical interaction between both proteins.

### Impact of Ephrin B1 on CXCR4 glycosylation of phosphorylation

Co-expression of Ephrin B1 with CXCR4 modified the pattern of CXCR4 migration in SDS-PAGE, leading to a single immunoreactive band at around 40 kDa, instead of two bands (the 40 kDa band and another band of higher apparent molecular weight) when the receptor was expressed alone, suggesting that Ephrin association with CXCR4 might affect CXCR4 post-translational modifications. Treatment of protein extracts from cells expressing the receptor alone with N-glycosidase F, but not O-glycosidase, led to the disappearance of the higher molecular weight band and to a receptor migration pattern identical to that observed in presence of Ephrin B1 (single band of apparent molecular weight of 40 kDa, **Figure 2A**). Moreover, N-glycosidase F treatment did not affect the receptor migration pattern in cells co-expressing Ephrin B1. Collectively, these results suggest that the co-expression of Ephrin B1 prevents CXCR4 N-glycosylation. Further supporting this hypothesis, Ephrin B1 expression did not affect the migration of a mutant CXCR4 where residues known to be N-glycosylated (positions 11 and 176) or O-glycosylated (position 18) were mutated into alanine (CXCR4ΔGlyc, **Figure 2A**).

**Figure 2:**
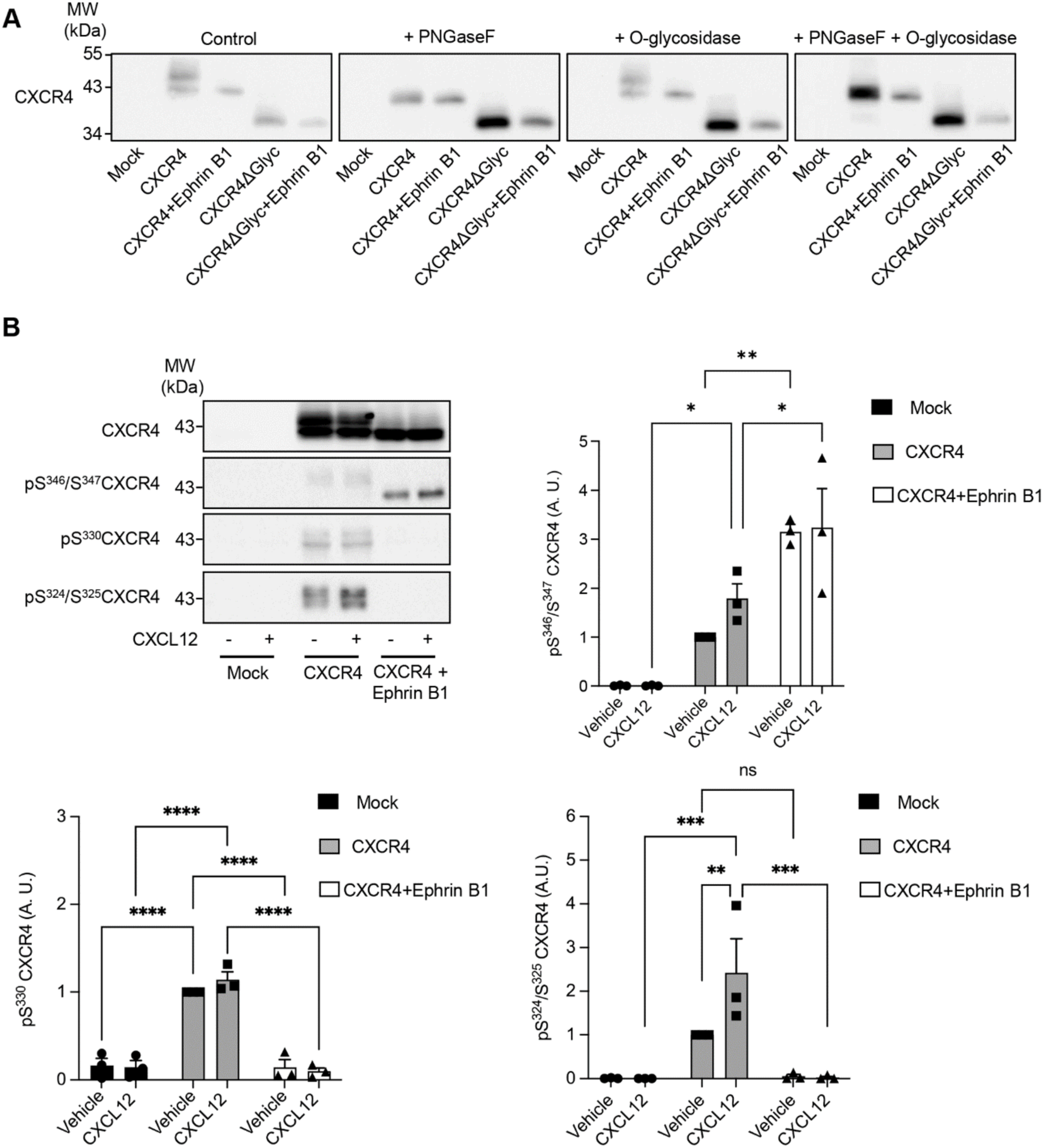
Impact of Ephrin B1 on CXCR4 glycosylation and phosphorylation A. Representative Western blots obtained from HEK293T cells expressing either wild-type HA-CXCR4 or CXCR4 mutated into alanine at positions 11, 18, and 176 (glycosylated residues, CXCR41′Glyc) in the presence or absence Flag-Ephrin B1. Cell lysates were treated or not with PNGaseF (10 IU/100 μg) and/or O-glycosidase (20 IU/100 μg) for 30 min at 37°C to remove N-glycosylation and/or O-glycosylation, respectively. The illustrated blots are representative of 3 independent experiments performed from different sets of cultured cells. **B** Representative WB performed from HEK293T cells, expressing or not HA-CXCR4 in the absence or presence of FLAG-Ephrin B1. Data are means ± SEM from 3 independent experiments performed from different sets of cultured cells. They were analyzed with the 2-way ANOVA with Bonferroni’s multiple comparison test. ns: non-significant.

We also examined the impact of Ephrin B1 expression on CXCR4 phosphorylation. CXCR4 is known to be sequentially phosphorylated on multiple residues located in its C-terminal domain, first on Ser^346/347^ by GRK2/3 and then on Ser^330^, Ser^324/325^ and Ser^338/339^ by GRK6 (Busillo et al., 2010; Mueller et al., 2013). Studies also suggested that PKC can phosphorylate Ser^346/347^ and Ser^324/325^ (Busillo et al., 2010; Mueller et al., 2013). Using phosphosite specific antibodies, we found that Ephrin B1 expression differentially affects the phosphorylation of Ser^346/347^, Ser^330^ and Ser^324/325^: while it enhanced Ser^346/347^ phosphorylation in cells exposed or not to CXCL12 for 5 min, it abolished the phosphorylation of both Ser^330^ and Ser^324/325^ in cells treated or not with CXCL12 (**Figure 2B**).

### Impact of Ephrin B1 on CXCR4 subcellular localization

Given the role of CXCR4 post-translational modifications in its trafficking, we next examined the effect of Ephrin B1 expression on plasma membrane localization of CXCR4-NLuc receptor using a nano-BRET assay and Neon-Green-LYN as an acceptor plasma membrane protein (Luís et al., 2022). The BRET signal decreased as a function of the amount of Ephrin B1 co-expressed with the receptor (**Figure 3A**). Note that the co-expression of Ephrin B1 did not modify the total expression of CXCR4, as assessed by the CXCR4-NLuc signal (**Supplementary Figure 2**). The decreased plasma membrane localization of CXCR4 elicited by Ephrin B1 co-expression was confirmed by confocal fluorescent microscopy in cells expressing CXCR4-YFP. While an important fraction of CXCR4-YFP was detected at the plasma membrane when it was expressed alone, it showed in cells co-expressing Ephrin B1-RFP a more pronounced distribution in intracellular compartments, where both proteins were colocalized (**Figures 1F** and **G** and **Supplementary Figure 3**). Co-immunostaining of cells co-expressing CXCR4-YFP and Ephrin B1-RFP (**Figures 1F** and **G**) with a calnexin (endoplasmic reticulum) or GM130 (Golgi apparatus) antibody showed an important fraction of the receptor in the endoplasmic reticulum and Golgi apparatus specifically in cells co-expressing CXCR4-YFP or Ephrin B1-RFP. Likewise, a fraction of the receptor was detected in the lysosomal compartment of cells co-expressing both proteins, as shown by co-immunostaining with a LAMP1 antibody, (**Figures 1F** and **G**).

**Figure 3:**
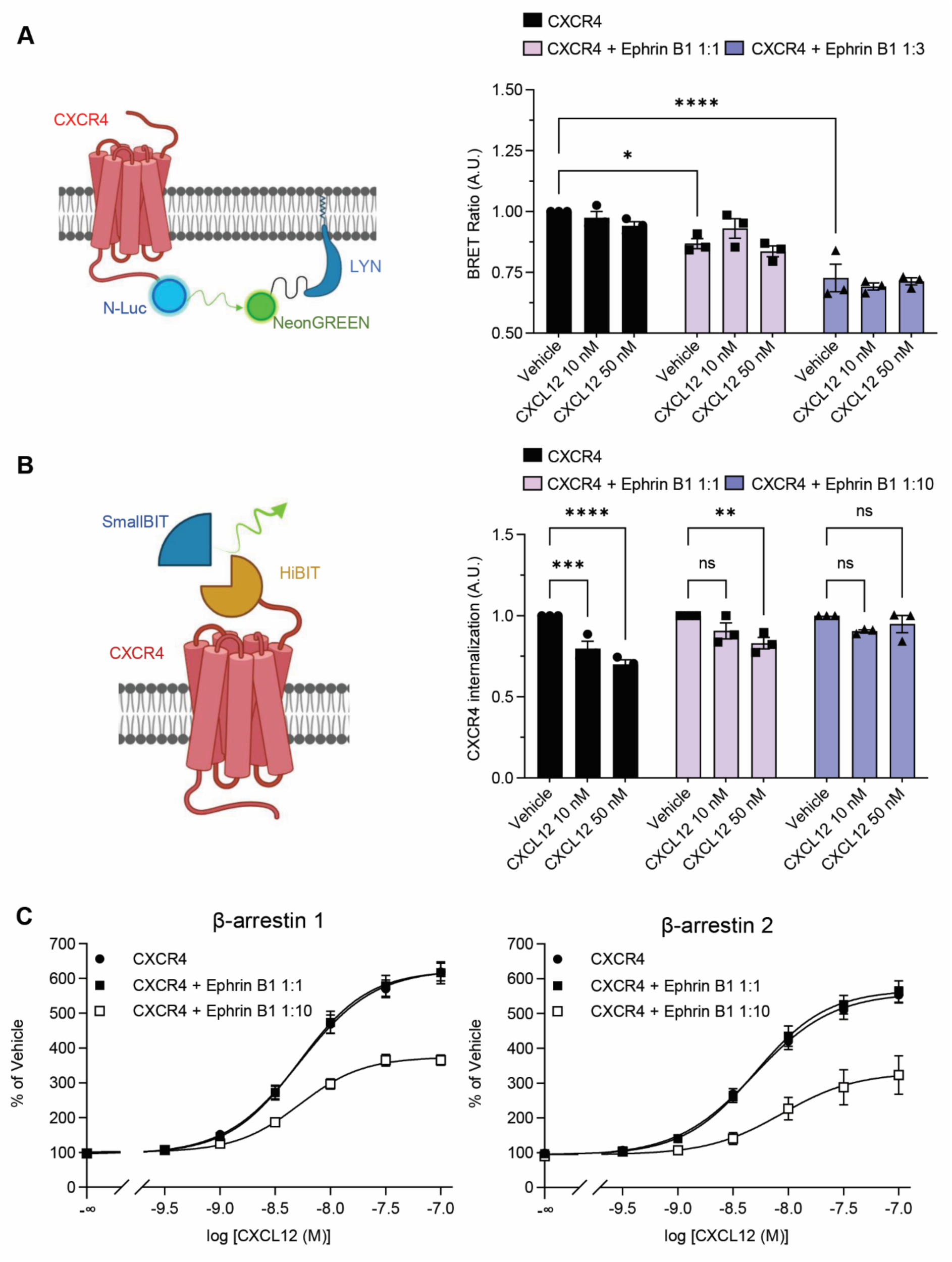
Influence of Ephrin B1 on CXCR4 cell surface expression and internalization. **A** *Left:* schematic representation of the nano-BRET assay assessing the interaction between CXCR4-NLuc and Lyn-mNeonGreen used to evaluate the plasma membrane localization of CXCR4 in cells expressing CXCR4 or coexpressing CXCR4 and Ephrin B1 (transfected plasmid ratios, 1:1 and 1:3). Cells were exposed to vehicle or CXCL12 (10 or 50 nM) for 60 min. *Right:* the data represent the BRET ratio normalized to the vehicle condition in cells expressing CXCR4-NLuc alone. They are the means ± SEM of values obtained in duplicate in three independent experiments performed on different sets of cultured cells. **B** *Left:* schematic representation of the nano-luciferase complementation assay used to evaluate the CXCL12-induced internalization of CXCR4. *Right:* the data represent CXCL12-induced internalization of CXCR4 normalized to vehicle in the corresponding condition, in cells expressing HiBIT-CXCR4 alone or coexpressing HiBIT-CXCR4 and FLAG-Ephrin B1 (plasmid ratios, 1:1 and 1:10). They are means ± SEM of duplicate determinations in three independent experiments performed on different sets of cultured cells. In **A** and **B**, the statistical analysis was the 2-way ANOVA with Bonferroni’s multiple comparison test. ns: non-significant. **C** β-arrestin 1 and β-arrestin 2 recruitment to CXCR4 in response to exposure to the indicated concentrations of CXCL12 in HEK293T cells co-expressing CXCR4-LgBIT and SmallBIT-β-arrestin1 or 2 in the absence or presence of FLAG-Ephrin B1 (CXCR4-LgBIT/FLAG-Ephrin B1 plasmid ratios, 1:1 and 1:10). The data are the means ± SEM of duplicate determinations performed in three independent experiments performed on different sets of cultured cells. Curves were fitted by the log[CXCL12] *vs.* luminescence non-linear regression using Prism.

Given the limited level of CXCR4 internalization measured upon stimulation by CXCL12 for 60 min in our nano-BRET assay using Neon-Green-LYN (**Figure 3A**), we assessed the impact of Ephrin B1 on CXCL12-induced CXCR4 internalization using a highly sensitive cell surface detection approach based on the HiBiT nanoluciferase complementation technology. Cells expressing N-terminally HiBiT-tagged CXCR4 were stimulated with CXCL12 and the remaining receptors at the plasma membrane were quantified by adding soluble LgBiT protein. As expected, treatment of cells expressing CXCR4 alone with CXCL12 (10 nM, 30 min) already induced a significant internalization of the receptor that was further enhanced in cells exposed to 50 nM CXCL12, while co-expression of Ephrin B1 prevented CXCL12-induced receptor internalization (**Figure 3B**). Consistent with these observations and with the β-arrestin-dependent CXCR4 internalization elicited by its activation by CXCL12 (Cheng et al., 2000), co-expression of Ephrin B1 with CXCR4 inhibited the recruitment of both β-arrestin 1 and β-arrestin 2 by the receptor, as assessed using a nanoluciferase complementation-based assay (**Figure 3C**).

### Impact of Ephrin B1 on CXCR4 signaling

Consistent with previous findings demonstrating that CXCR4 is canonically coupled with Gi proteins (Busillo and Benovic, 2007), saturation BRET analysis under conditions of constant RLuc-Gα_i1_ or RLuc-Gα_i3_ expression and increasing CXCR4-YFP expression level showed that CXCR4 recruits both Gα_i1 and_ Gα_i3_ proteins in HEK-293T cells (**Figure 4A**). Treatment of cells of CXCL12 did not modify the recruitment of both G proteins by the receptor, whereas it was strongly reduced by the co-expression of Ephrin B1 with the receptor. Corroborating these observations, Ephrin B1 expression also decreased the ability of CXCL12 stimulation of CXCR4 to promote dissociation Gα_i1_ and Gα_i3_ from Gβγ, as assessed by the decrease in the BRET signal between venus-Gψ_2_ and Gα_i1_-RLuc or Gα_i3_-RLuc (**Figure 4B**).

**Figure 4:**
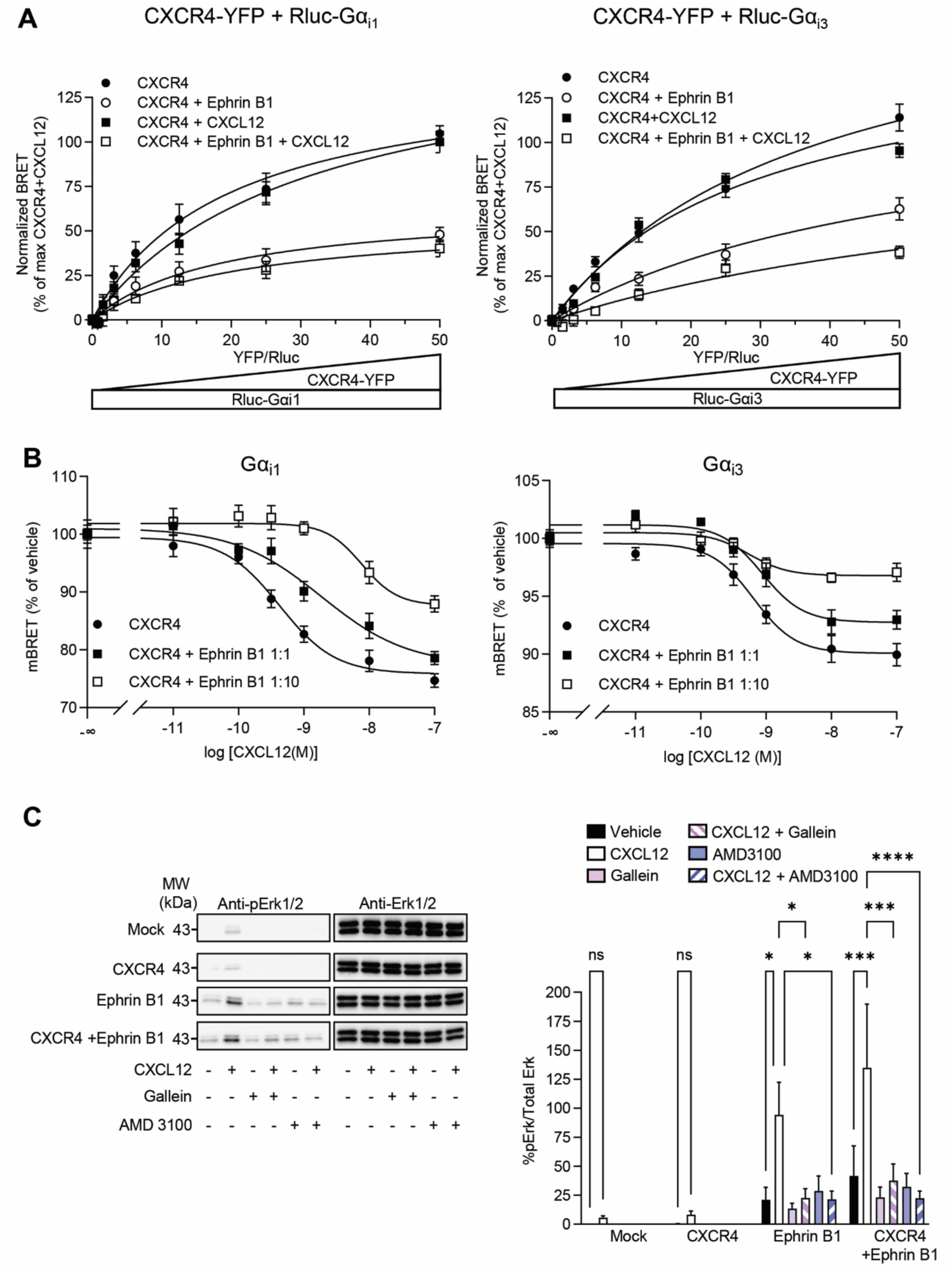
Influence of Ephrin B1 on CXCR4 signaling. **A** Gα_i1/3_ recruitment to CXCR4 in HEK293T cells expressing CXCR4-YFP alone or co-expressing CXCR4-YFP and FLAG-Ephrin B1, in presence of either Rluc-Gα_i1_ or Rluc-Gα_i3_ and exposed or not to CXCL12 (10 nM). BRET values were normalized to the maximum BRET measured in cells expressing CXCR4 and challenged with CXCL12. The data are means ± SEM of triplicate determinations performed in three independent experiments performed in different sets of cultured cells. Normalized BRET variations were fitted with the one-site total binding equation and background constraint to constant value of 0 using Prism. **B** Gα_i1/3_-Gβγ dissociation elicited by the indicated concentrations of CXCL12 exposure of HEK-293T cells expressing HACXCR4, alone or in combination with FLAG-Ephrin B1 (plasmid ratios 1:1 and 1:10), Rluc-Gαi1/3, Venus-γ2, and Flag-β2. The data were normalized to the values measured in vehicle-treated cells for in each condition. They are means ± SEM of triplicate determinations performed in three independent experiments performed in different sets of cultured cells. Curves were fitted by the log[CXCL12] *vs.* mBRET non-linear regression using Prism. **C** Representative Western blots performed from HEK293T cells expressing HA-CXCR4 or FLAG-Ephrin B1 or co-expressing HA-CXCR4 and FLAG-Ephrin B1, and treated with vehicle or CXCL12 (10 nM) for 5 min. When indicated, Gallein (25 µM) or AMD 3100 (1 µM) were added to cell 30 min before the vehicle/CXCL12 treatment. The data represent the pErk1,2/Total Erk1,2 ratio (in %). They are means ± SEM of values obtained in three independent experiments performed in different sets of cultured cells. They were analyzed with the 2-way ANOVA with Bonferroni’s multiple comparison test. ns: non-significant.

We next investigated the effect of Ephrin B1 expression on the ability of CXCR4 to activate the Erk1,2 pathway, a key signaling cascade underlying tumor progression elicited by the CXCL2-CXCR4 axis. As previously shown (Cassier et al., 2017), stimulation of CXCR4 endogenously expressed in HEK-293T cells with CXCL12 (10 nM, 5 min) induced an increase in Erk1,2 phosphorylation that was slightly but not significantly enhanced in cells transfected with a CXCR4 construct (**Figure 4C**). In contrast, the CXCL12 response was strongly enhanced in cells transfected with an Ephrin B1 construct alone or co-transfected with the CXCR4 and Ephrin B1 constructs. The strong Erk1,2 activation induced by CXCL12 in cells overexpressing Ephrin B1 did involve CXCR4 activation, as it was abolished by pretreating cells with the CXCR4 antagonist AMD3100 (1 μM, 30 min, **Figure 4C**). The activation of the Erk1,2 signaling cascade by GPCR ligands is often a complex process that involves G protein-dependent and β-arrestin-dependent mechanisms (Fumagalli et al., 2019). As shown on **Figure 3C**, expression of Ephrin B1 inhibitss β-arrestin 1 and β-arrestin 2 recruitment by CXCR4, making unlikely a role of β-arrestins in the enhanced CXCR4-operated Erk1,2 signaling observed in the presence of Ephrin B1. A previous study has demonstrated a role Gβγ and its translocation to the Golgi apparatus in Erk1,2 activation induced by CXCR4 activation in several cancer cell lines and HEK-293 cells (Khater et al., 2021). Consistent with these findings and the presence of CXCR4 and Ephrin B1 in Golgi apparatus in cells expressing both protein partners (**Figure 1F**), the activation of Erk1,2 induced by CXCL12 in cells expressing both protein partners (or overexpressing Ephrin B1 alone) was abolished by pretreating cells with the Gβγ pharmacological inhibitor Gallein (25 μM, 30 min, **Figure 4C**).

### Role of Ephrin B1 in the regulation of Death Receptor 5 expression by CXCR4 in breast cancer cells

Previous studies have shown that intracellular CXCR4 constitutively promotes the downregulation of Death Receptor 5 (DR5) independently of agonist stimulation of receptor signaling (Nengroo et al., 2021). In light of these findings and the ability of Ephrin B1 to promote intracellular localization of the receptor, we next examined the effect of CXCR4 and/or Ephrin B1 expression on DR5 level in MCF7 cells. Expression of CXCR4 alone significantly reduced DR5 levels (**Figure 5A**, left panel), consistent with the intracellular localization of the receptor (**Figure 5B**). Treatment of the cells with the CXCR4 antagonist AMD3100 (1 μM), did not affect the reduction of DR5 level elicited by CXCR4 expression (**Figure 5A**, left panel), indicating that it is independent of receptor stimulation. Expression of Ephrin B1 alone or in combination with CXCR4 reduced DR5 level to a similar extent as that measured in cells overexpressing CXCR4 (**Figure 5A**). To explore the role of Ephrin B1 in CXCR4-regulated DR5 expression, we silenced its expression using a shRNA. As expected, co-transfection of the Ephrin B1 shRNA with the Ephrin B1 plasmid, which induced plasma membrane localization of a fraction of CXCR4 (**Figure 5B**), restored a normal level of DR5 in cells (**Figure 5A**). Silencing Ephrin B1 expression also prevented the decrease in DR5 level elicited by CXCR4 overexpression, indicating a key role of Ephrin B1 in the regulation of Death Receptor 5 expression by CXCR4 in breast cancer cells.

**Figure 5:**
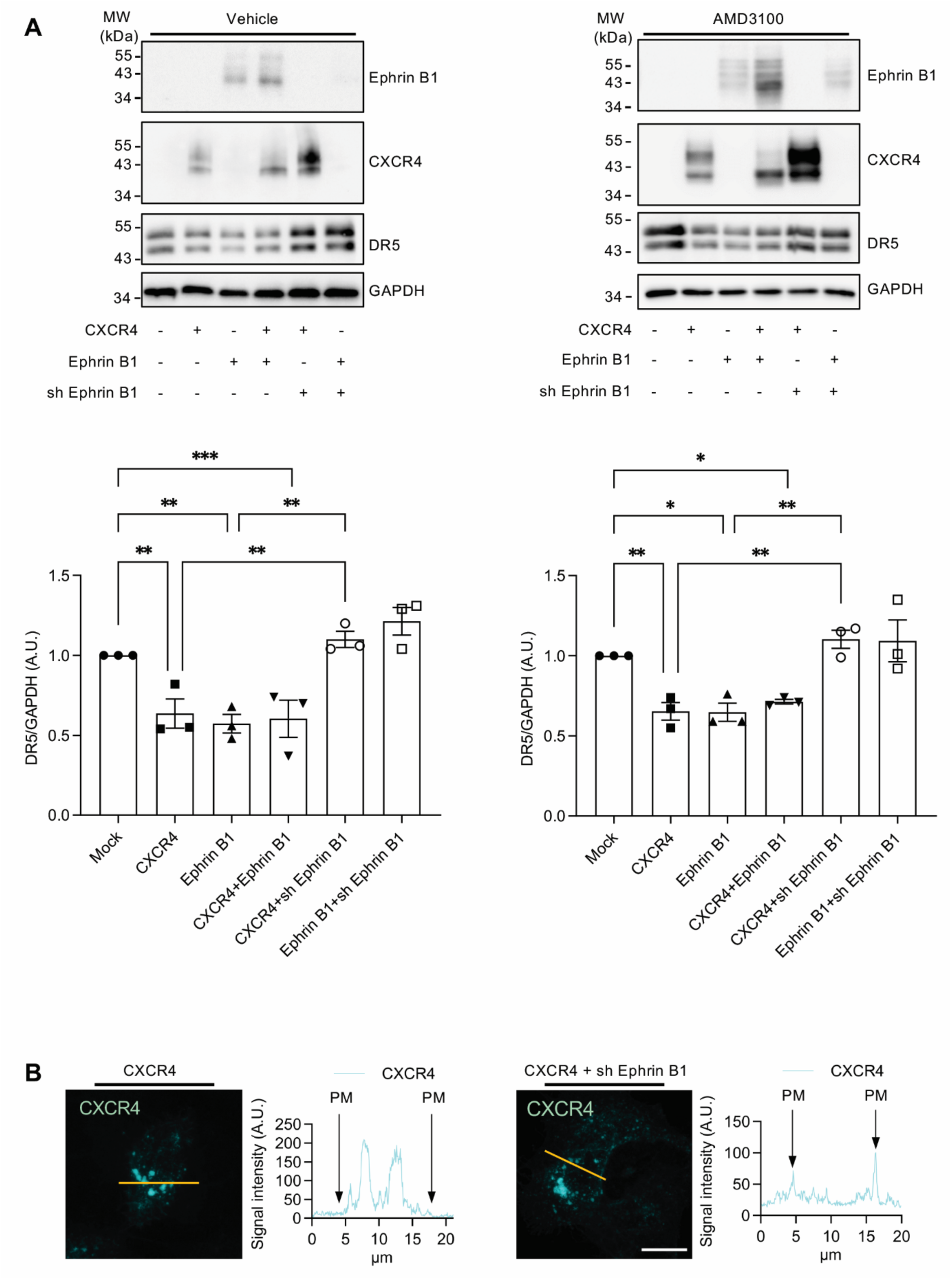
Role of Ephrin B1 on CXCR4 regulation of Death Receptor 5 expression. **A** Representative Western blots assessing Ephrin B1, CXCR4, DR5 and GAPDH expression in MCF7 cells transfected with constructs encoding HA-CXCR4 or FLAG-Ephrin B1 (alone or in combination) or constructs encoding HA-CXCR4 or FLAG-Ephrin B1 and a shRNA targeting Ephrin B1 and exposed to either vehicle (left panel) of AMD3100 (x 1μM, added to cells after the transfection, right panel). The histogram shows immunoreactive signals of DR5 normalized to GAPDH signal. The data represent the means ± SEM of values obtained in three independent experiments performed in different sets of cultured cells. They were analyzed with the 2-way ANOVA with Neuman Student Keul’s test. **B** Stacked images of CXCR4-YFP fluorescent signal in MCF7 cells expressing CXCR4-YFP alone or in combination with the Ephrin B1 shRNA. The illustrated fields are representative of three independent experiments performed from different sets of cultured cells. Scale bar: 10 μm. Line graphs generated in Image J using the lines represented on the images highlight the subcellular localization of CXCR4 in cells co-expressing or not the Ephrin B1 shRNA. PM, plasma membrane signal.

## Discussion

To identify mechanisms underlying tumorigenic effects of intracellular CXCR4 that are independent of receptor-operated signaling, we characterized the CXCR4 interactome and identified Ephrin B1 as a novel receptor-interacting protein. To our knowledge, this is the first demonstration of a physical interaction between a GPCR and a member of the Ephrin ligand family that comprises 5 GPI-anchored Ephrin-A ligands (Ephrins A1-5) and 3 transmembrane Ephrin-B ligands (Ephrins B1-3). Strikingly, Ephrin B1 was the most highly enriched protein identified in the CXCR4 interactome, suggesting a robust interaction between both partners. Corroborating this observation, we found that purified CXCR4 and Ephrin B1 can form a complex *in vitro*, indicative of a direct interaction between both proteins. The ability of Ephrin B1 to associate with CXCR4 in living cells was validated by BRET experiments, which also demonstrated that agonist stimulation of CXCR4 promotes the recruitment of Ephrin B1 by the receptor, indicating that CXCR4-Ephrin B1 interaction is a dynamic process regulated by the receptor’s conformational state. Furthermore, BRET experiments showed a low ability of Ephrin B1 to interact with ACKR3, an atypical chemokine receptor sharing with CXCR4 the chemokine CXCL12 as ligand, reminiscent of a previous interactomic screen performed in the same cellular background (HEK-293T cells) that did not identify Ephrin B1 as an ACKR3 partner (Fumagalli et al., 2020). As ACKR3 and CXCR4 are known to form heteromers in HEK-293T cells (Levoye et al., 2009), the low BRET signal measured in cells co-expressing ACKR3 and Ephrin B1 might reflect an indirect recruitment of Ephrin B1 through the heteromerization of ACKR3 with endogenously expressed CXCR4. Further supporting the specificity of CXCR4-Ephrin B1 interaction, Ephrin B1 was the only member of the Ephrin ligand family identified in our interactomic screen.

The common role of CXCR4 and Ephrin ligands and their receptors in cancer progression and resistance to chemotherapies prompted us to further investigate the reciprocal impact of Ephrin B1 and CXCR4 on their cellular localization and signaling. Using both a nanoBRET assay assessing plasma membrane localization of CXCR4 and confocal fluorescence microscopy, we provide converging evidence that Ephrin B1 expression promotes intracellular localization of the receptor in HEK-293T cells challenged or not with CXCL12. Likewise, while Ephrin B1 was mainly detected at the plasma membrane in cells expressing RFP-Ephrin B1 alone, it was predominantly found in intracellular compartments in cells co-expressing CXCR4-YFP. These include the endoplasmic reticulum and the Golgi apparatus as well as lysosomes, where both proteins were colocalized. Furthermore, using HiBIT surface Nano-luciferase complementation, we found that Ephrin B1 prevents rather than enhances CXCL12-induced CXCR4 internalization, an effect that correlates with the ability of Ephrin B1 to inhibit β-arrestin 1 and β-arrestin 2 recruitment by the receptor.

β-arrestin recruitment to GPCRs, which functions to promote their internalization, is known to depend on their phosphorylation on residues located in their C-terminal domain by GRKs and other protein kinases. Our results show that Ephrin B1 expression enhances CXCRX4 phosphorylation on Ser^346-347^, two residues phosphorylated by GRK2/3 located in the cytoplasm (Gardner et al., 2024; Touhara et al., 1994). In contrast, Ephrin B1 expression abolishes the phosphorylation of residues (Ser^324/325^ and Ser^330^) phosphorylated by GRK6, which is constitutively localized at the plasma membrane (Gardner et al., 2024; Jiang et al., 2007). Collectively, these findings are consistent with the predominant intracellular localization of the receptor in cells expressing Ephrin B1. CXCR4 phosphorylation at these sites also induce contrasting effects on β-arrestin recruitment and receptor internalization (Busillo et al., 2010; Mueller et al., 2013). Whereas the phosphorylation of Ser^346-347^ seems to be prerequisite to β-arrestin recruitment, inhibition of Ser^324/325^, Ser^330^ and Ser^339^ phosphorylation by their mutation into alanine favors β-arrestin recruitment (Busillo et al., 2010). Our results demonstrating that Ephrin B1 expression increases Ser^346-347^ phosphorylation while it abolishes Ser^324/325^ and Ser^330^ phosphorylation suggest that its effect on β-arrestin recruitment to CXCR4 (and receptor internalization) are independent of its modulation of the receptor phosphorylation state. Furthermore, the fact that Ephrin B1 inhibits CXCR4 internalization suggests that it promotes CXCR4 intracellular localization through the inhibition of its targeting to the plasma membrane. Consistent with this hypothesis, the ability of Ephrin B1 to associate with non-glycosylated CXCR4 with *in vitro* and to abolish its glycosylation suggests that CXCR4 and Ephrin B1 can associate early in the biosynthetic pathway, an effect leading to the inhibition of the forward trafficking and the intracellular sequestration of an important fraction of both proteins.

The decrease in CXCR4 plasma membrane localization in cells co-expressing Ephrin B1 might be one of the mechanisms underlying the inhibitory effect of Ephrin-B1 on CXCL12-induced Gα_i_ protein activation. A previous study has demonstrated that Ephrin B1 interacts with Regulator of G protein signaling 3 (RGS3), a member of the RGS protein family that negatively regulate G protein activation by GPCRs *via* the GTPase activating protein activity of their RGS domain. This results in the inhibition of the migration of cerebellar granule neurons induced by CXCL12 activation CXCR4 (Lu et al., 2001). It is likely that RGS3 linked to Ephrin B1 also contributes to the inhibition of Gα_i_ activation observed under expression of Ephrin B1, a process which could be favored by the physical interaction of Ephrin B1 with CXCR4.

Ephrin forward and reverse signaling involve extensive crosstalk with major cytosolic signaling pathways known to be engaged by tyrosine kinase receptors that control cell survival, migration and differentiation, such as Erk1,2 signaling (Boyd et al., 2014). Here, we show that Ephrin B1 expression that already increases Erk1,2 activation in absence of a CXCR4 agonist, strongly potentiates the CXCL12 response, while treatment of cells with AMD3100 abolished both basal and CXCL12-induced Erk1,2 activation. These findings indicate that Ephrin B1 engages Erk1,2 signaling through active CXCR4 even in the absence of extrinsic CXCL12. Further supporting the implication of CXCR4 in Ephrin B1-operated Erk1,2 signalling, the inhibition of Gβγ proteins which are essential for CXCR4-dependent Erk1,2 activation, also abolished basal and CXCL12-stimulated Erk1,2 activity in cells expressing Ephrin B1. Collectively, these observations are reminiscent of recent findings indicating that the EGF receptor and the HER3/HER2 receptors signal to Erk1,2 through active CXCR4/ACKR3 even in the absence of CXCL12 (Neves et al., 2022) and suggest that components of the Ephrin/Eph systems can likewise highjack the CXCR4 signaling machinery to activate the Erk1,2 pathway.

The synergistic effects of CXCR4 and Ephrin B1 on Erk1,2 signaling might be one of the mechanisms underlying their common deleterious influence on cancer cell growth, invasion and metastasis. Besides CXCR4-operated Erk signaling, it has recently been proposed that CXCR4 located intracellularly in cancer cells plays a critical role in the receptor’s tumorigenic effects *in vitro* and *in vivo* through the downregulation of DR5 expression (Nengroo et al., 2021). This prompted us to explore the influence of Ephrin B1 on DR5 expression in breast cancer cells. While Ephrin B1 over-expression reduced DR5 level in MCF7 cells to a similar extent as that measured in cells over-expressing CXCR4, its invalidation abolished the effect of CXCR4 expression, suggesting that Ephrin B1 regulates DR5 expression by promoting intracellular localization of CXCR4. Collectively, these findings indicate that Ephrin B1 can promote the tumorigenic potential CXCR4 through the enhancement of its oncogenic signaling and its sequestration in intracellular compartments where it inhibits cell apoptosis and thus renders cancer cells resistant to chemotherapeutic drugs. Although the role of the physical CXCR4-Ephrin B1 interaction could not be fully established in the absence of the identification of binding motifs within the sequences of both partners, they suggest that targeting Ephrin B1 reverse signaling and/or its interaction with CXCR4 might be a relevant strategy in complement to CXCR4 antagonists to dampen the tumorigenic effects of this receptor.

## Acknowledgements

We thank all our colleagues from the ONCORNET 2.0 consortium for continuous scientific discussions and support. LC-MS/MS analyses were performed using the facilities of the Functional Proteomics Platform managed by the service unit Biocampus Montpellier, BRET experiments using the facilities of the Arpege pharmacological screening platform (Biocampus Montpellier) and confocal microscopy image acquisition using the facilities of the Montpellier Ressources Imagerie (MRI) platform (Biocampus Montpellier). This research was funded by a European Union’s Horizon2020 MSCA Program (H2020-MSCA Program, Grant agreement 860229-ONCORNET 2.0). P.M. and S.C.D. are also supported by fundings from CNRS, INSERM and the University of Montpellier.

## Author contributions

A.R. designed, performed, and analyze most of the experiments, and wrote the manuscript; O.T produced purified recombinant CXCR4 and Ephrin B1; A.F. performed the interactomic screen; M. Seveno performed LC-MS/MS experiments and analyzed MS/MS data; S.G. performed confocal microscopy image acquisition and supervised image analysis; M.C. performed β-arrestins recruitment experiments; T.D. contributed to the design of experiments and manuscript **r**evision; M.J.S. participated in the project design and funding and revised the manuscript, M. Szpakowska designed and supervised Nano-BRET and Nano-luciferase Complementation experiments, and revised the manuscript; A.C. designed nano-BRET and nanoluciferase complementation experiments, and revised the manuscript, S.C.D. supervised the project, designed experiments and revised the manuscript; P.M. conceived and supervised the study, and wrote the manuscript.

