## Supplemental Data for "Physical interaction with Ephrin B1 promotes CXCR4 intracellular localization and oncogenic potential"

Alessandro Rabbito[1], Omolade Otun[1], Amos Fumagalli[1], Martial Séveno[2], Sonya Galant[1], Manuel Counson[3], Thierry Durroux[1], Cherine Bechara[1], Martine J Smit[4], Sébastien Granier[1], Martyna Szpakowska[3], Andy Chevigné[3], Séverine Chaumont-Dubel[1,2]*, Philippe Marin[1]*.

[1] Institut de Génomique Fonctionnelle, Université de Montpellier, CNRS, INSERM, Montpellier, France.

[2] Biocampus Montpellier, Université de Montpellier, CNRS, INSERM, Montpellier, France.

[3] Immuno-Pharmacology and Interactomics, Department of Infection and Immunity, Luxembourg Institute of Health, Esch-sur-Alzette, Luxembourg.

[4] Amsterdam Institute for Molecules Medicines and Systems, Division of Medicinal Chemistry, Faculty of Sciences, VU University Amsterdam, 1081 HV Amsterdam, The Netherlands.

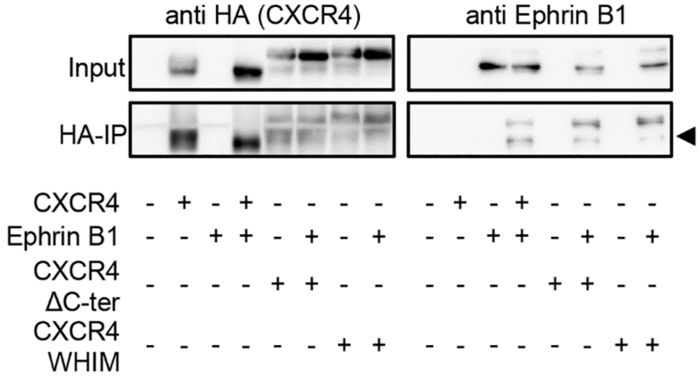


**Supplementary Figure 1: Co-immunoprecipitation of Ephrin B1 with CXCR4 with truncation of its C-terminal domain**. Western blots of HA immunoprecipitations performed from HEK293T cells expressing either wild type HA-CXCR4 or HA-CXCR41013 WHIM mutant (CXCR4-WHIM), and CXCR4 deleted of its entire C-terminal domain (HA-CXCR4ΔC-ter) alone or in combination with Flag-Ephrin B1. Note that none of the truncations affected co-immunoprecipitation of Ephrin B1 with CXCR4. The illustrated blots are representative of three independent experiments performed in different sets of cultured cells.

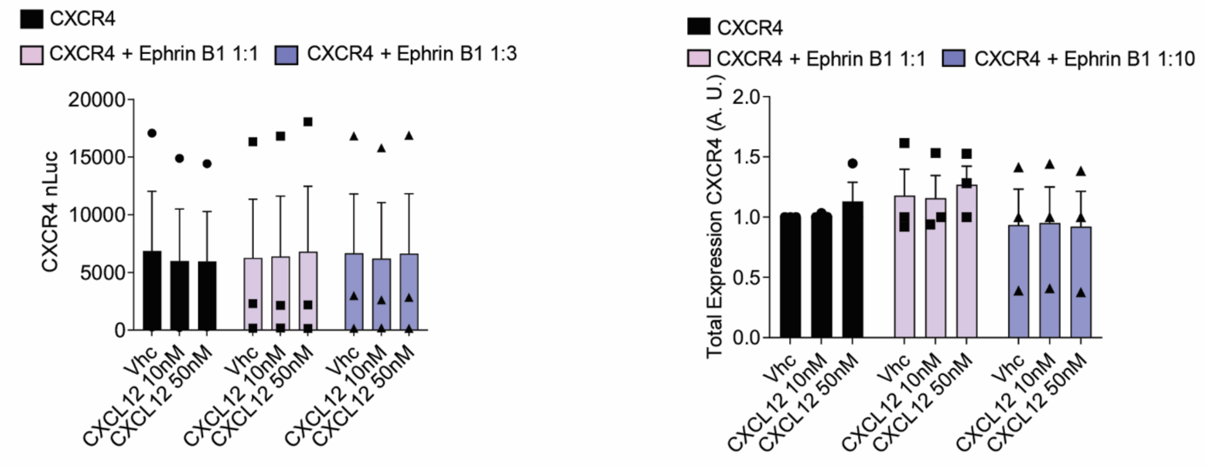


**Supplementary Figure 2: Total expression of CXCR4 in the NanoBRET and HiBIT experiments**

*Left*: the histograms show the CXCR4-NLuc signal measured in the NanoBRET-based assay, assessing CXCR4-NLuc-Lyn-mNeonGreen interaction in HEK293T cells (**Figure 3A**). The data are means ± SEM of duplicate determinations made in three independent experiments performed on different sets of cultured cells. *Right*: the histograms show the HiBIT-CXCR4 luminescence signal assessing the total expression of the receptor in the nano-luciferase complementation assay (**Figure 3B**). The data are means ± SEM of duplicate determinations made in three independent experiments performed on different sets of cultured cells.

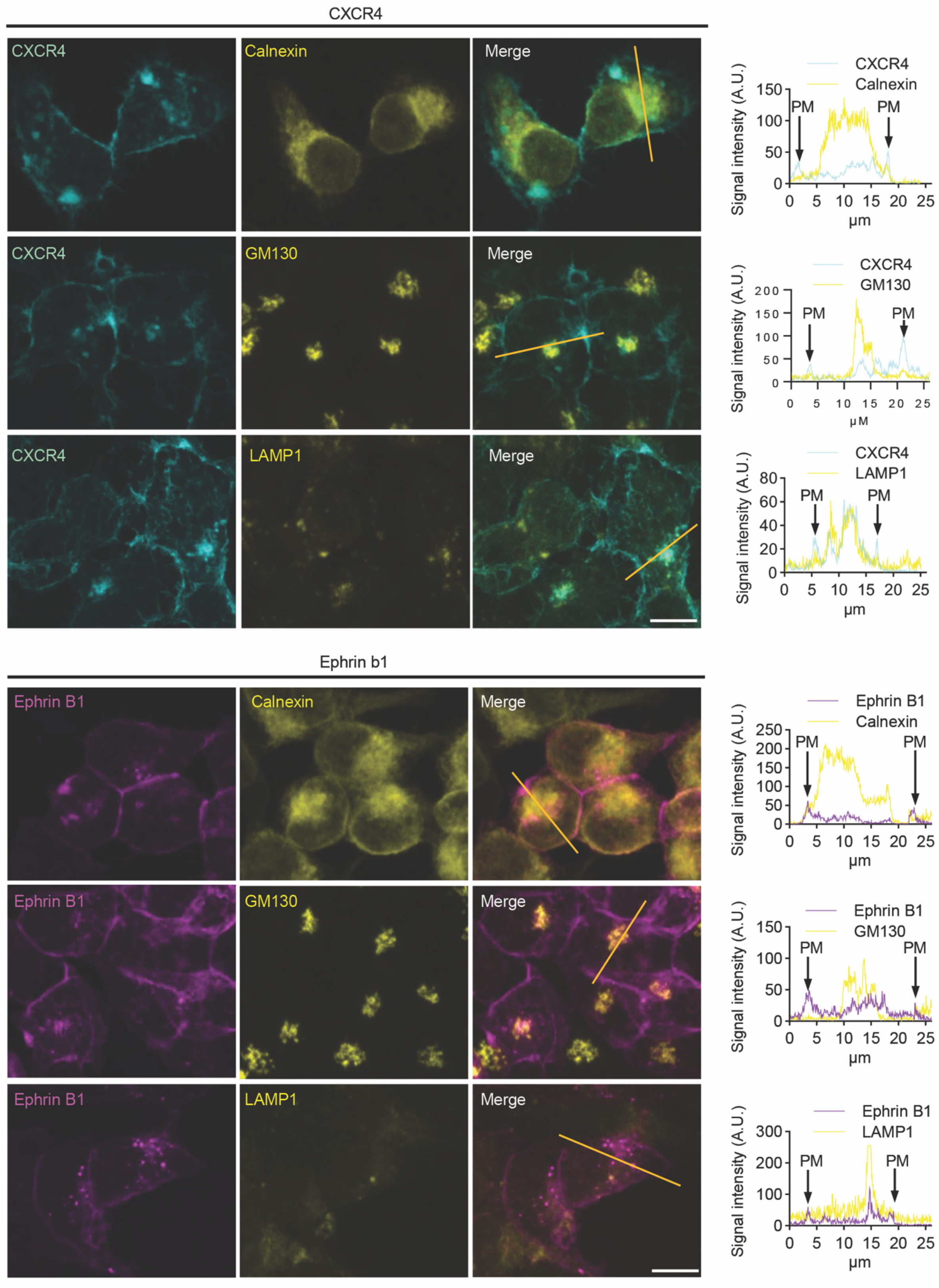


**Supplementary Figure 3:** Stacked images of CXCR4-YFP (cyan, top panels) and Ephrin B1-RFP (magenta, bottom panels) fluorescent signals in different cellular compartments in

HEK293T cells expressing either CXCR4-YFP or Ephrin B1-RFP alone. The anti-calnexin immunostaining was used to label the endoplasmic reticulum, the anti-GM130 immunostaining the cis-Golgi, and anti-LAMP1 immunostaining lysosomes (yellow). The illustrated fields are representative of three independent experiments performed from different sets of cultured cells. Scale bar: 10 μm. The right panels show line graphs generated in Image J using the lines represented on the merge images highlighting the subcellular localization of CXCR4 and Ephrin B1 in the three compartments. PM, plasma membrane signal.

**Supplementary Table 1**: List of proteins identified by LC-MS/MS analysis in immunoprecipitates from CXCR4-expressing HEK-293T cells and Mock cells. The protein names, gene names, – log P values (P value), LFQ difference between CXCR4-expressing cells and mock cells (Difference) and LFQ values measured in three biological replicates (CXCR4_1,CXCR4_2, CXCR4_3, *vs* MOCK_1, MOCK_2, MOCK_3) in both conditions are indicated. The statistical analysis was performed using the Perseus software, as detailed in the “Materials and Methods” section. Proteins were ranked according to their difference in abundance in immunoprecipitates from CXCR4-expressing cells *vs.* Mock cells. Proteins statistically enriched according to their LFQ level in the immunoprecipitates from CXCR4-expressing cells compared to Mock cells are indicated in left column.

| Significant | Protein names | UniProtID | Gene names | -LOG(P-value) | Difference | CXCR4_1 | CXCR4_2 | CXCR4_3 | MOCK_1 | MOCK_2 | MOCK_3 |
|---|---|---|---|---|---|---|---|---|---|---|---|
| + | C-X-C chemokine receptor type 4 | P61073 | CXCR4 | 4,81752 | 8,13412 | 23,07285 | 23,62912 | 24,20038 | 15,50000 | 15,50000 | 15,50000 |
| + | Ephrin-B1 | P98172 | EFNB1 | 4,95875 | 7,29111 | 23,29810 | 22,69285 | 22,38237 | 15,50000 | 15,50000 | 15,50000 |
| + | Speckle targeted PIP5K1A-regulated poly(A) polymerase | Q9H6E5 | TUT1 | 4,72740 | 7,05343 | 23,10594 | 22,08657 | 22,46777 | 15,50000 | 15,50000 | 15,50000 |
| + | Glutaredoxin-3 | O76003 | GLRX3 | 4,23064 | 5,96408 | 22,11975 | 21,25931 | 21,01318 | 15,50000 | 15,50000 | 15,50000 |
| + | Minor histocompatibility antigen H13 | Q8TCT9 | HM13 | 3,80840 | 5,81814 | 21,49897 | 21,93575 | 20,51970 | 15,50000 | 15,50000 | 15,50000 |
| + | 26S proteasome non-ATPase regulatory subunit 14 | O00487 | PSMD14 | 5,65340 | 5,69013 | 21,46129 | 21,11839 | 20,99072 | 15,50000 | 15,50000 | 15,50000 |
| + | Short/branched chain specific acyl-CoA dehydrogenase, mitochondrial | P45954 | ACADSB | 3,56233 | 5,65867 | 21,72995 | 21,52052 | 20,22553 | 15,50000 | 15,50000 | 15,50000 |
| + | Nuclear pore membrane glycoprotein 210 | Q8TEM1 | NUP210 | 5,33476 | 5,40417 | 21,14470 | 20,96769 | 20,60014 | 15,50000 | 15,50000 | 15,50000 |
| + | Electron transfer flavoprotein subunit beta | P38117 | ETFB | 3,59581 | 4,97812 | 20,68003 | 21,05795 | 19,69637 | 15,50000 | 15,50000 | 15,50000 |
| + | Beta-1,4-glucuronyltransferase 1 | O43505 | B4GAT1 | 3,82343 | 4,82992 | 20,79463 | 20,53811 | 19,65702 | 15,50000 | 15,50000 | 15,50000 |
| + | Baculoviral IAP repeat-containing protein 6 | Q9NR09 | BIRC6 | 3,81439 | 4,74226 | 20,88681 | 20,10789 | 19,73208 | 15,50000 | 15,50000 | 15,50000 |
| + | NEDD8-activating enzyme E1 regulatory subunit | Q13564 | NAE1 | 4,79763 | 4,72246 | 20,59933 | 20,08978 | 19,97827 | 15,50000 | 15,50000 | 15,50000 |
| + | Nucleoporin NDC1 | Q9BTX1 | NDC1 | 5,36129 | 4,69831 | 20,33156 | 20,33963 | 19,92373 | 15,50000 | 15,50000 | 15,50000 |
| + | FH1/FH2 domain-containing protein 1 | Q9Y613 | FHOD1 | 6,25869 | 4,61660 | 20,15561 | 19,96198 | 20,23222 | 15,50000 | 15,50000 | 15,50000 |
| + | Inositol-3-phosphate synthase 1 | Q9NPH2 | ISYNA1 | 4,05474 | 4,14048 | 19,83161 | 19,96000 | 19,12984 | 15,50000 | 15,50000 | 15,50000 |
| + | ER membrane protein complex subunit 4 | Q5J8M3 | EMC4 | 3,42766 | 4,04863 | 20,07091 | 19,72780 | 18,84717 | 15,50000 | 15,50000 | 15,50000 |
| + | COUP transcription factor 2;COUP transcription factor 1 | P24468 | NR2F2;NR2F1 | 5,77999 | 3,53283 | 18,87574 | 19,14629 | 19,07645 | 15,50000 | 15,50000 | 15,50000 |
| + | Origin recognition complex subunit 4 | O43929 | ORC4 | 4,20258 | 3,38933 | 18,51688 | 19,16828 | 18,98284 | 15,50000 | 15,50000 | 15,50000 |
| + | 26S protease regulatory subunit 6B | P43686 | PSMC4 | 4,02678 | 2,43658 | 22,64621 | 22,30955 | 22,79782 | 20,03979 | 20,21679 | 20,18725 |
| | 26S protease regulatory subunit 4 | P40939 | PSMC1 | 3,45567 | 1,74321 | 23,28644 | 23,07424 | 23,02672 | 21,52631 | 21,50954 | 21,12193 |
| | T-complex protein 1 subunit delta | Q7Z6Z7 | CCT4 | 3,32811 | 1,14567 | 24,61616 | 24,76921 | 24,78801 | 23,43122 | 23,75593 | 23,54922 |
| | Serine/threonine-protein phosphatase 6 regulatory subunit 1 | P55084 | PPP6R1 | 3,27061 | 3,60319 | 19,54278 | 19,36841 | 18,39837 | 15,50000 | 15,50000 | 15,50000 |
| | Evolutionarily conserved signaling intermediate in Toll pathway, mitochondrial | Q5T4S7 | ECSIT | 3,25481 | 4,15783 | 19,79478 | 20,29771 | 18,88101 | 15,50000 | 15,50000 | 15,50000 |
| | Ancient ubiquitous protein 1 | P78347 | AUP1 | 3,25013 | 4,78599 | 20,77820 | 20,75091 | 19,32885 | 15,50000 | 15,50000 | 15,50000 |
| | Clathrin heavy chain 1 | Q13501 | CLTC | 3,23548 | 2,70033 | 26,25288 | 25,62561 | 25,77603 | 23,50009 | 22,82499 | 23,22845 |
| | 60 kDa heat shock protein, mitochondrial | P01891 | HSPD1 | 3,13900 | 2,44292 | 27,09939 | 27,47251 | 27,24036 | 25,15128 | 24,36533 | 24,96688 |
| | T-complex protein 1 subunit alpha | Q99460 | TCP1 | 3,12317 | 1,16198 | 24,83012 | 25,06706 | 24,94060 | 23,87164 | 23,90536 | 23,57485 |
| | 26S protease regulatory subunit 8 | P49588 | PSMC5 | 3,12227 | 1,88044 | 23,33743 | 23,01708 | 22,79687 | 20,91284 | 21,30780 | 21,28940 |
| | D-3-phosphoglycerate dehydrogenase | O95071 | PHGDH | 3,01046 | 1,45074 | 24,82571 | 24,74904 | 24,73268 | 23,63179 | 23,25101 | 23,07240 |
| | Insulin receptor substrate 4 | Q9Y679 | IRS4 | 2,94165 | 1,22022 | 24,23710 | 24,10542 | 24,18033 | 23,23741 | 22,80453 | 22,82024 |
| | T-complex protein 1 subunit zeta | Q92621 | CCT6A | 2,94087 | 1,35975 | 24,11704 | 24,56046 | 24,27618 | 22,97164 | 23,12386 | 22,77892 |
| | Rap1 GTPase-GDP dissociation stimulator 1 | P04899 | RAP1GDS1 | 2,90383 | 2,96770 | 19,18171 | 17,97707 | 18,24433 | 15,50000 | 15,50000 | 15,50000 |
| | Serine--tRNA ligase, cytoplasmic | P54105 | SARS | 2,88861 | 0,83347 | 21,99638 | 22,09745 | 22,05841 | 21,26499 | 21,36030 | 21,02655 |
| | ATP-dependent Clp protease ATP-binding subunit clpX-like, mitochondrial | Q96P70 | CLPX | 2,85003 | 2,35380 | 21,64830 | 22,58868 | 22,16484 | 19,54219 | 19,96578 | 19,83245 |
| | T-complex protein 1 subunit epsilon | Q93008 | CCT5 | 2,81242 | 2,17370 | 24,08456 | 24,42490 | 24,59833 | 22,45566 | 21,71791 | 22,41310 |
| | C-1-tetrahydrofolate synthase, cytoplasmic;Methylenetetrahydrofolate dehydrogenase;Methenyltetrahydrofolate cyclohydrolase;Formyltetrahydrofolate synthetase;C-1-tetrahydrofolate synthase, cytoplasmic, N-terminally processed | Q15008 | MTHFD1 | 2,79294 | 2,19246 | 25,27391 | 24,79288 | 25,22041 | 22,53124 | 22,81259 | 23,36600 |
| | Serine/threonine-protein phosphatase 2A 65 kDa regulatory subunit A alpha isoform | Q13618 | PPP2R1A | 2,76017 | 3,19789 | 23,27355 | 24,14554 | 23,98720 | 20,98822 | 19,93589 | 20,88852 |
| | Probable ubiquitin carboxyl-terminal hydrolase FAF-X | Q99720 | USP9X | 2,66261 | 4,34572 | 24,65622 | 24,01333 | 23,25283 | 19,87030 | 20,29098 | 18,72394 |
| | Importin-5 | P61289 | IPO5 | 2,64482 | 2,11153 | 22,90857 | 22,57434 | 22,91612 | 21,24685 | 20,33385 | 20,48375 |
| | Cullin-associated NEDD8-dissociated protein 1 | Q14C86 | CAND1 | 2,61003 | 1,74660 | 24,10254 | 23,70504 | 23,73159 | 22,27736 | 22,36509 | 21,65692 |
| | Importin-4 | P22102 | IPO4 | 2,58782 | 3,78381 | 22,22609 | 21,71377 | 22,44397 | 17,30117 | 18,89288 | 18,83834 |
| | Trifunctional purine biosynthetic protein adenosine-3;Phosphoribosylamine--glycine ligase;Phosphoribosylformylglycinamidine cyclo-ligase;Phosphoribosylglycinamide formyltransferase | Q9BQ95 | GART | 2,57891 | 4,17181 | 23,56256 | 22,10130 | 22,69806 | 18,99153 | 19,15581 | 17,69914 |
| | Protein disulfide-isomerase TMX3 | O75694 | TMX3 | 2,51829 | 3,56909 | 19,68271 | 19,56588 | 17,95867 | 15,50000 | 15,50000 | 15,50000 |
| | Protein SCO2 homolog, mitochondrial | O00429 | SCO2 | 2,46759 | 3,16143 | 18,66543 | 19,54021 | 17,77867 | 15,50000 | 15,50000 | 15,50000 |
| | Cytochrome b-c1 complex subunit 2, mitochondrial | Q9HDC9 | UQCRC2 | 2,38290 | 1,60280 | 23,20247 | 22,76749 | 22,45671 | 21,53431 | 21,01686 | 21,06710 |
| | Ran GTPase-activating protein 1 | P16615 | RANGAP1 | 2,37116 | 1,65137 | 22,74255 | 22,71207 | 22,89345 | 21,54962 | 21,23561 | 20,60874 |
| | Leucine-rich PPR motif-containing protein, mitochondrial | Q9BSJ8 | LRPPRC | 2,36334 | 3,79437 | 24,99352 | 24,69853 | 24,97058 | 20,67951 | 22,35762 | 20,24237 |
| | Proteasome subunit beta type-5 | Q8IZP2 | PSMB5 | 2,32726 | 2,58087 | 20,83823 | 21,00690 | 19,91382 | 18,59550 | 17,80751 | 17,61332 |
| | 26S protease regulatory subunit 7 | Q31612 | PSMC2 | 2,32209 | 2,26809 | 24,09200 | 23,25997 | 23,47680 | 20,73281 | 21,76979 | 21,52191 |
| | Importin-9 | P35613 | IPO9 | 2,30595 | 4,36191 | 20,89734 | 22,44488 | 22,93720 | 17,24841 | 18,68147 | 17,26380 |
| | Cancer-related nucleoside-triphosphatase | P13073 | NTPCR | 2,29397 | 1,10627 | 22,18047 | 22,50359 | 22,47783 | 21,61840 | 21,13715 | 21,08751 |
| | Polypeptide N-acetylgalactosaminyltransferase 2;Polypeptide N-acetylgalactosaminyltransferase 2 soluble form | P17980 | GALNT2 | 2,29034 | 2,74906 | 20,60503 | 21,78141 | 21,25891 | 18,75551 | 17,75277 | 18,88988 |
| | T-complex protein 1 subunit beta | Q9UNM6 | CCT2 | 2,28965 | 1,28297 | 24,74950 | 24,11315 | 24,48640 | 23,43746 | 23,08305 | 22,97963 |

| Significant | Protein names | UniProtID | Gene names | -LOG(P-value) | Difference | CXCR4_1 | CXCR4_2 | CXCR4_3 | MOCK_1 | MOCK_2 | MOCK_3 |
|---|---|---|---|---|---|---|---|---|---|---|---|
| | Pachytene checkpoint protein 2 homolog | P42704 | TRIP13 | 2,25751 | 1,16107 | 21,51307 | 21,34147 | 21,35143 | 19,90654 | 20,61621 | 20,20000 |
| | Aldehyde dehydrogenase X, mitochondrial | Q7Z434 | ALDH1B1 | 2,25716 | 2,74168 | 21,73020 | 22,66540 | 21,96010 | 19,14954 | 20,18689 | 18,79424 |
| | Lysine-specific demethylase PHF2 | Q8TEX9 | PHF2 | 2,23501 | -0,76396 | 20,29053 | 19,93200 | 20,28568 | 21,08971 | 20,84153 | 20,86884 |
| | Dolichyl-diphosphooligosaccharide--protein glycosyltransferase subunit 2 | P49736 | RPN2 | 2,21383 | 1,61082 | 23,03080 | 23,15558 | 23,25551 | 21,41080 | 21,09610 | 22,10252 |
| | E3 ubiquitin-protein ligase HUWE1 | P0DN79 | HUWE1 | 2,20827 | 6,52108 | 24,02775 | 22,42116 | 23,04747 | 15,50000 | 18,93313 | 15,50000 |
| | HLA class I histocompatibility antigen, B-73 alpha chain | P31948 | HLA-B | 2,10536 | 3,88698 | 21,30691 | 20,95163 | 20,19785 | 15,50000 | 17,70192 | 17,59354 |
| | WD repeat-containing protein 6 | P27824 | WDR6 | 2,06896 | 1,19597 | 23,67842 | 23,06847 | 23,34881 | 21,87211 | 22,15857 | 22,47712 |
| | Aladin | Q13200 | AAAS | 2,02814 | 1,01424 | 21,07385 | 21,35159 | 20,79321 | 20,29457 | 20,08408 | 19,79730 |
| | T-complex protein 1 subunit theta | Q9UPN7 | CCT8 | 2,01760 | 1,87513 | 24,96080 | 24,47797 | 24,32540 | 23,35891 | 22,64123 | 22,13865 |
| | DNA damage-binding protein 1 | P00403 | DDB1 | 2,01639 | 1,58930 | 23,20599 | 22,10815 | 22,48581 | 21,09706 | 20,78556 | 21,14942 |
| | Stomatin-like protein 2, mitochondrial | Q96JJ7 | STOML2 | 2,00974 | 1,59112 | 23,58635 | 23,23078 | 22,62415 | 21,72255 | 21,78281 | 21,16257 |
| | Ornithine aminotransferase, mitochondrial;Ornithine aminotransferase, hepatic form;Ornithine aminotransferase, renal form | P07741 | OAT | 1,96052 | 1,13304 | 22,30683 | 21,71209 | 22,35308 | 20,96339 | 21,25603 | 20,75344 |
| | Proteasome activator complex subunit 3 | Q9Y678 | PSME3 | 1,94708 | 4,20564 | 22,20275 | 21,16140 | 20,88569 | 15,50000 | 17,97544 | 18,15749 |
| | Chromobox protein homolog 3 | P08195 | CBX3 | 1,94631 | -1,04403 | 20,65508 | 21,42552 | 20,91510 | 22,16717 | 22,00175 | 21,95886 |
| | M-phase phosphoprotein 8 | P54578 | MPHOSPH8 | 1,94548 | -0,86145 | 21,63436 | 21,26402 | 21,06197 | 22,35617 | 22,01798 | 22,17054 |
| | General transcription factor II-I | O00231 | GTF2I | 1,93359 | 5,42658 | 21,58907 | 22,30805 | 22,48623 | 15,50000 | 19,10362 | 15,50000 |
| | T-complex protein 1 subunit eta | O14818 | CCT7 | 1,92130 | 1,28232 | 23,40626 | 22,75595 | 23,25897 | 22,25517 | 21,81449 | 21,50455 |
| | Tubulin alpha-1B chain;Tubulin alpha-4A chain | P30566 | TUBA1B;TUBA4A | 1,90861 | 1,41444 | 28,05181 | 28,81546 | 28,94679 | 26,86302 | 27,43213 | 27,27560 |
| | Protein LSM12 homolog | Q96KG9 | LSM12 | 1,90853 | -0,58177 | 20,56709 | 20,78245 | 20,92796 | 21,33685 | 21,19748 | 21,48846 |
| | Guanine nucleotide-binding protein G(s) subunit alpha isoforms XLas | O95433 | GNAS | 1,88486 | 1,44935 | 22,06822 | 21,74164 | 21,25632 | 20,72700 | 20,03738 | 19,95376 |
| | Probable ATP-dependent RNA helicase DDX52 | Q96CS3 | DDX52 | 1,87461 | -0,71148 | 22,55614 | 22,95692 | 22,40101 | 23,37994 | 23,29179 | 23,37677 |
| | TAR DNA-binding protein 43 | Q12769 | TARDBP | 1,87352 | 0,94403 | 23,42047 | 24,10021 | 23,61246 | 22,94172 | 22,74129 | 22,61802 |
| | Mitochondrial antiviral-signaling protein | O15269 | MAVS | 1,85631 | 3,78537 | 20,17365 | 20,38174 | 19,99714 | 15,50000 | 18,19643 | 15,50000 |
| | Transcription intermediary factor 1-beta | Q96TC7 | TRIM28 | 1,81956 | 1,02553 | 22,54769 | 21,77913 | 22,10217 | 21,15443 | 20,89903 | 21,29894 |
| | Calcium-binding mitochondrial carrier protein Aralar2 | P30153 | SLC25A13 | 1,81456 | 0,86756 | 23,31631 | 23,14088 | 23,27284 | 22,19430 | 22,78873 | 22,14433 |
| | Importin subunit beta-1 | O43819 | KPNB1 | 1,81195 | 1,71105 | 24,11219 | 23,21010 | 23,91324 | 22,62252 | 21,96356 | 21,51629 |
| | CAD protein;Glutamine-dependent carbamoyl-phosphate synthase;Aspartate carbamoyltransferase;Dihydroorotase | P62873 | CAD | 1,80953 | 1,35576 | 24,84196 | 24,75379 | 24,60145 | 23,94592 | 22,81147 | 23,37252 |
| | Importin subunit alpha-1 | Q9C0E8 | KPNA2 | 1,80094 | 0,84991 | 23,28560 | 22,88457 | 22,96749 | 22,52848 | 22,10789 | 21,95156 |
| | Kelch-like ECH-associated protein 1 | P62834 | KEAP1 | 1,79924 | 0,61851 | 20,50252 | 20,62783 | 20,45663 | 19,97365 | 19,63354 | 20,12427 |
| | Glycine--tRNA ligase | O95292 | GARS | 1,78788 | 1,86383 | 22,55925 | 23,24818 | 22,50135 | 20,19592 | 21,58454 | 20,93682 |
| | Guanine nucleotide-binding protein G(i) subunit alpha-2 | Q9BQE3 | GNAI2 | 1,77469 | 4,49028 | 21,28546 | 21,53564 | 20,39791 | 15,50000 | 18,74816 | 15,50000 |
| | Sideroflexin-1 | P54709 | SFXN1 | 1,77114 | 1,13004 | 22,30444 | 22,45151 | 21,73322 | 21,40047 | 20,81253 | 20,88606 |
| | Fatty acid synthase;[Acyl-carrier-protein] S-acetyltransferase;[Acyl-carrier-protein] S-malonyltransferase;3-oxoacyl-[acyl-carrier-protein] synthase;3-oxoacyl-[acyl-carrier-protein] reductase;3-hydroxyacyl-[acyl-carrier-protein] dehydratase;Enoyl-[acyl-carrier-protein] reductase;Oleoyl-[acyl-carrier-protein] hydrolase | P15924 | FASN | 1,75704 | 1,67169 | 26,04806 | 25,43479 | 25,34571 | 24,66649 | 23,49668 | 23,65031 |
| | Tyrosine--tRNA ligase, cytoplasmic;Tyrosine--tRNA ligase, cytoplasmic, N-terminally processed | O75165 | YARS | 1,75684 | 1,48522 | 22,02181 | 21,86155 | 22,15174 | 20,47060 | 21,19556 | 19,91327 |
| | Tubulin beta-4B chain | Q13190 | TUBB4B | 1,75088 | 1,16365 | 27,89082 | 28,43293 | 28,20401 | 27,40154 | 27,10270 | 26,53257 |
| | E3 ubiquitin-protein ligase UBR4 | Q9BUF5 | UBR4 | 1,74813 | 6,01371 | 23,75460 | 22,91379 | 22,36268 | 15,50000 | 19,98995 | 15,50000 |
| | ATP synthase subunit beta, mitochondrial | P13861 | ATP5B | 1,74545 | 2,12614 | 24,30445 | 24,16300 | 24,74566 | 21,86333 | 21,65905 | 23,31230 |
| | Sodium/potassium-transporting ATPase subunit alpha-1 | P49792 | ATP1A1 | 1,73737 | 2,25676 | 23,90141 | 25,18499 | 24,93049 | 21,56913 | 22,66049 | 23,01698 |
| | 26S proteasome non-ATPase regulatory subunit 1 | P51570 | PSMD1 | 1,73369 | 5,04289 | 22,48517 | 21,48458 | 21,46742 | 15,50000 | 19,30853 | 15,50000 |
| | DNA mismatch repair protein Msh6 | Q96QK1 | MSH6 | 1,73285 | 1,14831 | 21,90947 | 21,82981 | 21,44784 | 20,53848 | 21,05616 | 20,14756 |
| | T-complex protein 1 subunit gamma | P27105 | CCT3 | 1,73044 | 1,61492 | 24,30758 | 24,69179 | 24,81787 | 23,68185 | 22,32237 | 22,96827 |
| | ATP synthase subunit alpha, mitochondrial | Q8NEW0 | ATP5A1 | 1,72059 | 0,98375 | 25,98470 | 26,47800 | 26,31456 | 25,56726 | 25,40051 | 24,85823 |
| | 26S proteasome non-ATPase regulatory subunit 13 | P52306 | PSMD13 | 1,69889 | 3,80965 | 21,62230 | 21,51927 | 20,37124 | 15,50000 | 18,14694 | 18,43693 |
| | ER membrane protein complex subunit 2 | P53992 | EMC2 | 1,69463 | 1,31120 | 21,13740 | 20,92774 | 20,37994 | 19,88189 | 18,98393 | 19,64567 |
| | Heat shock protein 105 kDa | Q8TC12 | HSPH1 | 1,69054 | 1,39323 | 22,21154 | 21,81699 | 21,92405 | 21,06874 | 19,89719 | 20,80696 |
| | U3 small nucleolar RNA-associated protein 15 homolog | P48556 | UTP15 | 1,68310 | -1,47735 | 19,20748 | 20,12540 | 20,11095 | 21,42752 | 21,65626 | 20,79210 |
| | Regulator of microtubule dynamics protein 3 | Q9NUQ9 | RMDN3 | 1,68204 | 3,25651 | 20,07874 | 19,53127 | 19,17874 | 15,50000 | 18,01922 | 15,50000 |
| | 26S proteasome non-ATPase regulatory subunit 12 | P55786 | PSMD12 | 1,67289 | 1,71113 | 21,57279 | 21,22152 | 21,32103 | 20,37496 | 18,81997 | 19,78703 |
| | Cystathionine beta-synthase | P62333 | CBS | 1,66306 | 3,64694 | 20,01318 | 20,82135 | 19,29318 | 15,50000 | 15,50000 | 18,18689 |
| | General transcription factor 3C polypeptide 5 | Q9H936 | GTF3C5 | 1,65641 | 0,44109 | 21,09275 | 21,18453 | 21,07267 | 20,90764 | 20,57432 | 20,54472 |
| | Histone acetyltransferase type B catalytic subunit | P37268 | HAT1 | 1,65102 | 1,17314 | 22,83389 | 22,56774 | 21,88684 | 21,52931 | 20,97694 | 21,26282 |
| | Chloride channel CLIC-like protein 1 | Q04323 | CLCC1 | 1,62770 | 2,32272 | 18,76523 | 18,13260 | 16,57033 | 15,50000 | 15,50000 | 15,50000 |
| | Proliferating cell nuclear antigen | P61923 | PCNA | 1,61521 | 1,41537 | 23,56639 | 23,89319 | 23,54132 | 23,00850 | 21,75324 | 21,99307 |
| | Regulator of nonsense transcripts 3B | Q12800 | UPF3B | 1,59434 | -1,34293 | 21,61720 | 22,39076 | 22,37013 | 23,58348 | 23,90536 | 22,91803 |
| | Sequestosome-1 | P13804 | SQSTM1 | 1,59209 | 5,10105 | 22,37684 | 22,38638 | 21,32025 | 15,50000 | 19,78032 | 15,50000 |
| | Nuclear pore complex protein Nup205 | A1L0T0 | NUP205 | 1,57596 | 4,67674 | 23,16896 | 22,75833 | 21,83723 | 18,24961 | 19,98468 | 15,50000 |
| | HLA class I histocompatibility antigen, A-68 alpha chain | O75436 | HLA-A | 1,57534 | 5,08036 | 22,27884 | 22,32017 | 21,51903 | 15,50000 | 19,87697 | 15,50000 |
| | Small subunit processome component 20 homolog | Q8N2G8 | UTP20 | 1,57145 | -0,75948 | 26,10661 | 25,88124 | 26,27508 | 26,75888 | 27,21252 | 26,56997 |

| Significant | Protein names | UniProtID | Gene names | -LOG(P-value) | Difference | CXCR4_1 | CXCR4_2 | CXCR4_3 | MOCK_1 | MOCK_2 | MOCK_3 |
|---|---|---|---|---|---|---|---|---|---|---|---|
| | Nicalin | P53004 | NCLN | 1,56987 | 1,61165 | 21,71716 | 21,43324 | 21,00949 | 19,43237 | 20,61979 | 19,27278 |
| | Adenine phosphoribosyltransferase | Q7L1Q6 | APRT | 1,56779 | 3,54370 | 21,74193 | 20,81339 | 19,80670 | 15,50000 | 18,27294 | 17,95799 |
| | Ubiquitin-40S ribosomal protein S27a;Ubiquitin;40S ribosomal protein S27a | Q9NTJ5 | RPS27A | 1,56707 | 1,51094 | 26,01472 | 24,71177 | 24,77379 | 23,71931 | 23,84019 | 23,40795 |
| | AP-2 complex subunit mu | Q02790 | AP2M1 | 1,54865 | -1,17308 | 23,82357 | 24,63312 | 24,45432 | 25,06636 | 25,92325 | 25,44063 |
| | Syntaxin-5 | P20618 | STX5 | 1,54660 | 3,04346 | 19,47730 | 19,43854 | 19,43469 | 15,50000 | 15,50000 | 18,22015 |
| | Exportin-2 | Q10471 | CSE1L | 1,53250 | 1,91865 | 22,47316 | 22,75269 | 23,42150 | 20,11108 | 21,85742 | 20,92289 |
| | RNA-binding protein 8A | P30837 | RBM8A | 1,52650 | -1,32498 | 21,16968 | 22,37348 | 21,79654 | 22,84751 | 23,49729 | 22,96983 |
| | Nuclear pore complex protein Nup155 | P43246 | NUP155 | 1,51452 | 4,13878 | 22,31979 | 19,14512 | 19,93258 | 15,50000 | 17,98114 | 15,50000 |
| | Cofilin-1 | P61163 | CFL1 | 1,50554 | 0,56785 | 24,71827 | 24,69530 | 24,64532 | 24,34428 | 24,23279 | 23,77826 |
| | Replication factor C subunit 1 | P60900 | RFC1 | 1,49555 | -0,51231 | 25,94217 | 26,28762 | 26,36528 | 26,54149 | 26,85210 | 26,73841 |
| | Ubiquitin thioesterase OTUB1 | P62879 | OTUB1 | 1,49302 | 1,58347 | 22,46785 | 21,82559 | 21,26671 | 20,95319 | 19,81903 | 20,03752 |
| | Mediator of RNA polymerase II transcription subunit 8 | P78344 | MED8 | 1,48876 | -4,11262 | 15,50000 | 15,50000 | 19,28667 | 21,27742 | 20,77169 | 20,57542 |
| | Hydroxysteroid dehydrogenase-like protein 2 | Q00610 | HSDL2 | 1,48821 | 0,95249 | 20,84797 | 21,41431 | 20,65692 | 20,37857 | 19,73041 | 19,95276 |
| | Sigma non-opioid intracellular receptor 1 | P00387 | SIGMAR1 | 1,48673 | 4,28949 | 21,42029 | 21,33702 | 20,52888 | 15,50000 | 19,41772 | 15,50000 |
| | E3 ubiquitin-protein ligase UBR5 | P31930 | UBR5 | 1,47882 | 4,92210 | 22,74893 | 21,13934 | 21,78875 | 15,50000 | 19,91072 | 15,50000 |
| | Alanine--tRNA ligase, cytoplasmic | P57678 | AARS | 1,47077 | 4,93441 | 22,43990 | 20,99189 | 22,32647 | 15,50000 | 15,50000 | 19,95504 |
| | Stress-induced-phosphoprotein 1 | O75489 | STIP1 | 1,46888 | 3,64055 | 20,31130 | 20,43665 | 20,11108 | 15,50000 | 15,50000 | 18,93739 |
| | Cdc42 effector protein 1 | O00505 | CDC42EP1 | 1,46504 | 2,33499 | 18,47374 | 18,75281 | 18,47987 | 15,50000 | 17,70144 | 15,50000 |
| | Histone-binding protein RBBP4 | P00367 | RBBP4 | 1,45463 | -1,12643 | 22,95040 | 23,72910 | 22,83326 | 24,67998 | 24,30862 | 23,90343 |
| | Phosphatidylinositide phosphatase SAC1 | Q07065 | SACM1L | 1,45099 | 2,77751 | 19,56162 | 19,10628 | 18,73364 | 15,50000 | 18,06902 | 15,50000 |
| | 26S proteasome non-ATPase regulatory subunit 3 | P28074 | PSMD3 | 1,44709 | 1,14593 | 24,30013 | 24,03246 | 23,58314 | 23,43071 | 22,54823 | 22,49899 |
| | Zinc transporter 7 | O15270 | SLC30A7 | 1,44564 | 2,96848 | 20,02012 | 19,35503 | 18,62542 | 15,50000 | 18,09512 | 15,50000 |
| | Ataxin-10 | P36957 | ATXN10 | 1,44436 | 2,03257 | 23,02159 | 22,45513 | 21,64592 | 21,35751 | 19,65725 | 20,01018 |
| | Nuclear pore complex protein Nup160 | Q7Z3B4 | NUP160 | 1,44050 | 3,30369 | 19,85206 | 19,33499 | 20,30908 | 15,50000 | 18,58505 | 15,50000 |
| | Putative protein FAM10A4;Hsc70-interacting protein;Putative protein FAM10A5 | P06280 | ST13P4;ST13;ST13P5 | 1,43938 | 3,90427 | 20,86801 | 20,53877 | 20,57626 | 15,50000 | 19,27023 | 15,50000 |
| | Ras-related protein Rab-6A;Ras-related protein Rab-6B | P61081 | RAB6A;RAB6B | 1,43645 | 1,01062 | 22,78307 | 22,62845 | 22,33060 | 22,08226 | 21,04667 | 21,58132 |
| | Adipocyte plasma membrane-associated protein | Q9UI09 | APMAP | 1,40330 | 3,99899 | 22,75426 | 21,91419 | 21,31008 | 15,50000 | 18,92289 | 19,55867 |
| | Ribose-phosphate pyrophosphokinase 2 | Q5H9R7 | PRPS2 | 1,40152 | 1,07147 | 23,22587 | 22,34530 | 22,56065 | 21,41684 | 22,11518 | 21,38538 |
| | Putative methyltransferase C9orf114 | P19623 | C9orf114 | 1,39869 | -1,41788 | 19,75880 | 21,27906 | 20,40458 | 22,22288 | 21,64161 | 21,83159 |
| | Sphingosine-1-phosphate lyase 1 | O95163 | SGPL1 | 1,38910 | -0,98989 | 22,83545 | 23,79333 | 23,37637 | 24,52880 | 23,95908 | 24,48696 |
| | Pyruvate dehydrogenase E1 component subunit beta, mitochondrial | O75351 | PDHB | 1,38243 | 1,53283 | 23,61054 | 22,37201 | 22,36685 | 21,79776 | 20,72176 | 21,23140 |
| | Alpha-aminoadipic semialdehyde dehydrogenase | O96000 | ALDH7A1 | 1,38003 | 1,61052 | 22,19994 | 21,83731 | 20,92071 | 20,71364 | 20,04860 | 19,36414 |
| | Exosome complex exonuclease RRP44 | P22314 | DIS3 | 1,37909 | 1,48348 | 21,10194 | 19,71549 | 20,51884 | 19,54284 | 18,53526 | 18,80773 |
| | Trifunctional enzyme subunit alpha, mitochondrial;Long-chain enoyl-CoA hydratase;Long chain 3-hydroxyacyl-CoA dehydrogenase | P10809 | HADHA | 1,37471 | 7,00215 | 26,93888 | 27,21058 | 27,54995 | 15,50000 | 22,88686 | 22,30608 |
| | Tubulin beta-2B chain | Q9BZZ5 | TUBB2B | 1,37132 | 1,10305 | 21,64064 | 21,85658 | 21,25275 | 21,03631 | 20,51489 | 19,88962 |
| | Extended synaptotagmin-1 | Q14318 | ESYT1 | 1,36387 | 3,92400 | 21,22205 | 20,13496 | 20,83268 | 15,50000 | 19,41770 | 15,50000 |
| | Proteasome subunit alpha type-7;Proteasome subunit alpha type-7-like | P28331 | PSMA7;PSMA8 | 1,36270 | 3,48832 | 21,34294 | 21,54298 | 21,21945 | 15,50000 | 18,94650 | 19,19393 |
| | Ras-related protein Rab-1B;Putative Ras-related protein Rab-1C | P08754 | RAB1B;RAB1C | 1,36053 | 0,70945 | 22,27660 | 22,44670 | 22,77250 | 21,94424 | 21,40088 | 22,02232 |
| | Importin subunit alpha-7 | O75521 | KPNA6 | 1,35781 | 1,07586 | 21,55267 | 21,92267 | 21,12098 | 20,98197 | 19,98593 | 20,40083 |
| | ATP-dependent zinc metalloprotease YME1L1 | Q5VYK3 | YME1L1 | 1,35529 | 1,57589 | 21,87185 | 22,12272 | 22,36429 | 19,51289 | 20,89052 | 21,22776 |
| | Coatomer subunit delta | Q13445 | ARCN1 | 1,35049 | 0,83921 | 20,34981 | 21,20258 | 21,06494 | 20,26899 | 19,87168 | 19,95901 |
| | Mitochondrial import receptor subunit TOM22 homolog | O76031 | TOMM22 | 1,34738 | 2,32402 | 22,96120 | 22,72084 | 21,19436 | 21,10642 | 19,64635 | 19,15155 |
| | Heat shock protein HSP 90-beta | Q9HCU5 | HSP90AB1 | 1,34153 | 1,22334 | 25,66998 | 27,00496 | 26,83364 | 25,38969 | 25,31941 | 25,12947 |
| | GTPase-activating protein and VPS9 domain-containing protein 1 | Q00587 | GAPVD1 | 1,34086 | 4,19329 | 21,90286 | 20,66371 | 20,73248 | 15,50000 | 19,71918 | 15,50000 |
| | Cytochrome c oxidase subunit 4 isoform 1, mitochondrial | Q9NS69 | COX4I1 | 1,34028 | 3,84659 | 22,88560 | 21,12547 | 21,36917 | 15,50000 | 19,08892 | 19,25153 |
| | UDP-glucose 6-dehydrogenase | Q96S66 | UGDH | 1,33740 | 1,00051 | 22,18510 | 21,98287 | 21,52200 | 21,37326 | 20,94256 | 20,37263 |
| | BUB3-interacting and GLEBS motif-containing protein ZNF207 | Q9P2R3 | ZNF207 | 1,33612 | -0,94184 | 21,68136 | 22,49502 | 22,55756 | 23,18826 | 22,89010 | 23,48112 |
| | C-terminal-binding protein 2 | P29401 | CTBP2 | 1,33174 | 0,92788 | 21,50736 | 20,43654 | 21,08589 | 20,08771 | 19,91224 | 20,24621 |
| | Nuclear migration protein nudC | Q15717 | NUDC | 1,33160 | 0,96621 | 20,97190 | 20,44385 | 20,07718 | 19,91686 | 19,15943 | 19,51801 |
| | DnaJ homolog subfamily C member 11 | O75794 | DNAJC11 | 1,33017 | 0,58846 | 21,02506 | 21,67564 | 21,54312 | 20,78085 | 20,94285 | 20,75475 |
| | Heat shock protein HSP 90-alpha | Q13243 | HSP90AA1 | 1,32782 | 1,23988 | 23,76020 | 24,64818 | 24,75297 | 22,86652 | 23,75277 | 22,82242 |
| | PHD finger protein 6 | Q93009 | PHF6 | 1,32377 | -1,38103 | 21,10961 | 22,20066 | 22,43522 | 22,84424 | 23,27341 | 23,77092 |
| | Tyrosine-protein kinase Yes;Proto-oncogene tyrosine-protein kinase Src | P35998 | YES1;SRC | 1,32032 | 1,44810 | 20,05616 | 19,57804 | 18,42370 | 18,24451 | 17,72833 | 17,74077 |
| | Ubiquitin carboxyl-terminal hydrolase 14 | P05023 | USP14 | 1,31329 | 3,50776 | 20,65491 | 20,49209 | 19,45613 | 15,50000 | 19,07984 | 15,50000 |
| | GH3 domain-containing protein | P21912 | GHDC | 1,30848 | 2,80975 | 19,63433 | 19,33956 | 18,90551 | 15,50000 | 18,45016 | 15,50000 |
| | N-terminal kinase-like protein | P34897 | SCYL1 | 1,30616 | 3,41863 | 20,02052 | 20,05656 | 20,34451 | 15,50000 | 15,50000 | 19,16570 |
| | Methylosome subunit pICln | Q9H6S0 | CLNS1A | 1,29715 | 4,43860 | 21,69060 | 21,20246 | 21,70809 | 15,50000 | 20,28534 | 15,50000 |
| | ATP-dependent RNA helicase DDX19B | Q9BPX3 | DDX19B | 1,29320 | 1,09378 | 21,92332 | 21,34234 | 21,07189 | 20,89297 | 20,33232 | 19,83091 |
| | Adenylosuccinate lyase | P11586 | ADSL | 1,28919 | 3,47835 | 20,70742 | 20,11235 | 19,82983 | 15,50000 | 19,21457 | 15,50000 |
| | Importin subunit alpha-3 | O14681 | KPNA4 | 1,28882 | 1,27580 | 20,66752 | 20,90228 | 19,78362 | 19,75352 | 19,10408 | 18,66841 |
| | U5 small nuclear ribonucleoprotein 40 kDa protein | P48643 | SNRNP40 | 1,27925 | -0,99255 | 21,43812 | 21,70126 | 21,96930 | 22,05444 | 23,15210 | 22,87980 |
| | DNA-directed RNA polymerase I subunit RPA43 | P33176 | TWISTNB | 1,27387 | -1,39302 | 22,01546 | 23,65031 | 22,49231 | 24,37981 | 24,14872 | 23,80863 |

| Significant | Protein names | UniProtID | Gene names | -LOG(P-value) | Difference | CXCR4_1 | CXCR4_2 | CXCR4_3 | MOCK_1 | MOCK_2 | MOCK_3 |
|---|---|---|---|---|---|---|---|---|---|---|---|
| | Transitional endoplasmic reticulum ATPase | P25685 | VCP | 1,27375 | 1,24533 | 23,01483 | 23,42265 | 23,72754 | 22,95945 | 21,67810 | 21,79150 |
| | Protein disulfide-isomerase A6 | P49593 | PDIA6 | 1,27117 | 1,13934 | 21,64610 | 22,06671 | 23,04968 | 21,08362 | 21,23444 | 21,02641 |
| | Annexin A2;Putative annexin A2-like protein | Q12789 | ANXA2;ANXA2P2 | 1,26740 | -0,59501 | 25,55652 | 25,55098 | 25,59713 | 25,87786 | 26,59548 | 26,01633 |
| | Cyclin-dependent kinase 1 | Q15785 | CDK1 | 1,26524 | 1,27705 | 22,01941 | 22,26905 | 21,18918 | 21,22735 | 20,12010 | 20,29905 |
| | 26S proteasome non-ATPase regulatory subunit 6 | P06576 | PSMD6 | 1,26412 | 4,30611 | 21,61540 | 21,27889 | 21,30847 | 15,50000 | 20,28444 | 15,50000 |
| | Zinc finger C3H1 domain-containing protein | Q00535 | ZFC3H1 | 1,26337 | -0,81773 | 22,34762 | 22,19596 | 22,17649 | 23,20029 | 23,48922 | 22,48375 |
| | Basic leucine zipper and W2 domain-containing protein 1 | P07099 | BZW1 | 1,25110 | 2,79638 | 20,04260 | 18,89894 | 18,87565 | 15,50000 | 18,42804 | 15,50000 |
| | 26S proteasome non-ATPase regulatory subunit 8 | O00410 | PSMD8 | 1,23534 | 2,94172 | 20,81785 | 20,77370 | 20,20801 | 15,50000 | 18,35832 | 19,11606 |
| | IQ calmodulin-binding motif-containing protein 1 | Q8NBU5 | IQCB1 | 1,23495 | -1,08303 | 20,75866 | 21,70383 | 21,76166 | 22,36407 | 22,97851 | 22,13066 |
| | Protein HIRA | Q8NBM4 | HIRA | 1,22832 | -0,74315 | 21,03591 | 21,04140 | 20,25200 | 21,72080 | 21,49414 | 21,34381 |
| | Dynamin-1-like protein | Q96L92 | DNM1L | 1,22127 | 4,09694 | 23,14784 | 22,67327 | 22,23809 | 15,50000 | 20,42777 | 19,84061 |
| | Ubinuclein-2 | Q13685 | UBN2 | 1,21455 | -3,42921 | 15,50000 | 15,50000 | 19,33793 | 20,91079 | 19,87767 | 19,83709 |
| | Phosphatidylinositol 5-phosphate 4-kinase type-2 beta | Q9H0U9 | PIP4K2B | 1,20889 | -1,18432 | 19,99162 | 21,25759 | 20,60748 | 21,45844 | 22,35762 | 21,59359 |
| | Mannose-1-phosphate guanyltransferase alpha | P04350 | GMPPA | 1,20720 | 1,43621 | 21,30930 | 20,54858 | 19,81190 | 19,82547 | 18,70209 | 18,83359 |
| | Trifunctional enzyme subunit beta, mitochondrial;3-ketoacyl-CoA thiolase | Q96I99 | HADHB | 1,20511 | 6,11913 | 25,53609 | 26,13012 | 27,20574 | 15,50000 | 22,49614 | 22,51841 |
| | Chromobox protein homolog 5 | Q8NF37 | CBX5 | 1,20171 | -5,33414 | 15,50000 | 15,50000 | 21,44738 | 21,54335 | 23,67820 | 23,22823 |
| | 26S protease regulatory subunit 6A | Q9UBB4 | PSMC3 | 1,19651 | 3,82753 | 21,92416 | 22,17005 | 22,72763 | 15,50000 | 19,60025 | 20,23900 |
| | Tubulin beta-4A chain | P18031 | TUBB4A | 1,18889 | 2,05337 | 20,84115 | 22,15208 | 22,52667 | 20,91503 | 19,71158 | 18,73317 |
| | A-kinase anchor protein 8 | P09622 | AKAP8 | 1,18397 | 0,92816 | 21,05384 | 21,97554 | 21,37459 | 20,09726 | 20,96522 | 20,55703 |
| | Pre-mRNA-splicing factor SLU7 | Q9NSE4 | SLU7 | 1,18344 | -1,52223 | 20,21122 | 22,16362 | 21,75621 | 22,87560 | 22,72182 | 23,10031 |
| | L-lactate dehydrogenase A chain | P55060 | LDHA | 1,17599 | 1,01229 | 23,36145 | 22,05096 | 22,94099 | 21,85361 | 21,53469 | 21,92821 |
| | Peptidyl-prolyl cis-trans isomerase FKBP4;Peptidyl-prolyl cis-trans isomerase FKBP4, N-terminally processed | O94905 | FKBP4 | 1,17254 | 2,77644 | 20,38005 | 18,58498 | 18,72067 | 15,50000 | 18,35638 | 15,50000 |
| | Serine palmitoyltransferase 1 | Q8NDZ4 | SPTLC1 | 1,16346 | 3,29467 | 19,31966 | 20,17719 | 20,69617 | 15,50000 | 19,30899 | 15,50000 |
| | Electron transfer flavoprotein subunit alpha, mitochondrial | P62195 | ETFA | 1,16205 | 2,82281 | 20,82858 | 20,82135 | 19,85240 | 15,50000 | 18,56335 | 18,97054 |
| | Proteasome subunit beta type-1 | P50990 | PSMB1 | 1,16082 | 2,77044 | 20,23585 | 20,41287 | 20,68577 | 15,50000 | 18,33595 | 19,18723 |
| | Eukaryotic translation initiation factor 4 gamma 1 | P67775 | EIF4G1 | 1,14409 | 1,07531 | 24,86844 | 23,71712 | 24,60801 | 22,95718 | 23,15670 | 23,85377 |
| | 4F2 cell-surface antigen heavy chain | P41250 | SLC3A2 | 1,14240 | 3,53422 | 21,59860 | 21,72134 | 22,38377 | 15,50000 | 19,88351 | 19,71755 |
| | Ribose-phosphate pyrophosphokinase 1 | Q9BPX5 | PRPS1 | 1,13823 | 0,90741 | 21,52061 | 21,46159 | 21,13652 | 21,11927 | 20,38015 | 19,89707 |
| | Voltage-dependent anion-selective channel protein 1 | Q9BT78 | VDAC1 | 1,12346 | 0,69998 | 25,33187 | 25,45441 | 25,07484 | 24,26799 | 25,12566 | 24,36753 |
| | Basigin | Q8N7H5 | BSG | 1,12015 | 3,85616 | 22,94639 | 22,11731 | 22,61029 | 15,50000 | 20,31363 | 20,29188 |
| | Sister chromatid cohesion protein PDS5 homolog A | O96008 | PDS5A | 1,11883 | 1,45579 | 21,79206 | 22,08745 | 22,29617 | 19,62072 | 21,67327 | 20,51432 |
| | Bifunctional ATP-dependent dihydroxyacetone kinase/FAD-AMP lyase (cyclizing);ATP-dependent dihydroxyacetone kinase;FAD-AMP lyase (cyclizing) | O14980 | DAK | 1,11736 | 1,68303 | 19,06179 | 18,82910 | 18,52742 | 18,37882 | 15,99440 | 16,99601 |
| | Growth arrest and DNA damage-inducible proteins-interacting protein 1 | P11802 | GADD45GIP1 | 1,11439 | -3,13938 | 15,50000 | 15,50000 | 19,47297 | 20,02296 | 19,96663 | 19,90152 |
| | Actin-like protein 6A | Q53H96 | ACTL6A | 1,10760 | -0,69220 | 23,88120 | 23,45237 | 22,88582 | 24,16883 | 24,14188 | 23,98529 |
| | U3 small nucleolar RNA-interacting protein 2 | Q9UNH7 | RRP9 | 1,10760 | -4,24193 | 15,50000 | 15,50000 | 20,87298 | 21,63710 | 21,16601 | 21,79567 |
| | Glyceraldehyde-3-phosphate dehydrogenase | Q53EL6 | GAPDH | 1,10540 | 1,46563 | 27,32492 | 25,59172 | 26,67438 | 25,43787 | 25,42143 | 24,33484 |
| | Glutamate-rich WD repeat-containing protein 1 | Q16637 | GRWD1 | 1,10327 | -1,28302 | 21,97043 | 22,74774 | 23,60964 | 23,52325 | 24,42426 | 24,22935 |
| | Translocation protein SEC62 | O15160 | SEC62 | 1,10134 | -0,75544 | 23,47951 | 23,96067 | 22,88902 | 24,24438 | 24,32526 | 24,02589 |
| | 14-3-3 protein theta | Q9Y6E2 | YWHAQ | 1,09985 | 1,36999 | 23,79303 | 24,22589 | 22,74729 | 22,93292 | 21,59951 | 22,12383 |
| | Cytoskeleton-associated protein 4 | Q86VP6 | CKAP4 | 1,09076 | 2,58499 | 21,82353 | 23,35097 | 23,55532 | 20,36688 | 18,62242 | 21,98555 |
| | Synaptic vesicle membrane protein VAT-1 homolog | P62191 | VAT1 | 1,09073 | 1,23129 | 21,81839 | 20,96614 | 20,18192 | 20,18435 | 19,34774 | 19,74051 |
| | SNW domain-containing protein 1 | P36776 | SNW1 | 1,08848 | -1,00854 | 20,83923 | 22,05126 | 22,18571 | 22,58699 | 22,65938 | 22,85546 |
| | Dolichyl pyrophosphate Man9GlcNAc2 alpha-1,3-glucosyltransferase | O75822 | ALG6 | 1,08146 | -3,46545 | 15,50000 | 15,50000 | 19,78957 | 21,19472 | 20,43736 | 19,55384 |
| | RNA-binding protein Raly | O00232 | RALY | 1,07151 | -1,52168 | 19,58924 | 20,57838 | 21,52425 | 21,44758 | 22,09623 | 22,71310 |
| | Translocon-associated protein subunit alpha | Q14974 | SSR1 | 1,07127 | 1,16890 | 20,14271 | 20,25038 | 21,66778 | 19,52658 | 19,25958 | 19,76801 |
| | UBX domain-containing protein 1 | Q99518 | UBXN1 | 1,06921 | 2,87867 | 21,18948 | 20,99148 | 20,40530 | 15,50000 | 19,33521 | 19,11505 |
| | RNA exonuclease 4 | Q8WXF1 | REXO4 | 1,06115 | -3,14384 | 15,50000 | 17,76156 | 18,82025 | 20,24540 | 22,33350 | 18,93445 |
| | Exportin-1 | Q9UKV8 | XPO1 | 1,06110 | 1,80636 | 22,73076 | 22,23570 | 22,99151 | 19,69379 | 22,30368 | 20,54142 |
| | Eukaryotic translation initiation factor 2 subunit 1 | Q3LXA3 | EIF2S1 | 1,05969 | -1,13226 | 22,52965 | 22,35644 | 23,86183 | 23,72848 | 24,24213 | 24,17410 |
| | Cullin-3 | P11233 | CUL3 | 1,05898 | 4,29818 | 19,36089 | 23,31008 | 24,01044 | 15,50000 | 19,59536 | 18,69149 |
| | KRR1 small subunit processome component homolog | P49327 | KRR1 | 1,05868 | -1,00344 | 19,73907 | 19,85050 | 20,60576 | 20,37973 | 21,54312 | 21,28280 |
| | AP-2 complex subunit beta | Q9Y2Z0 | AP2B1 | 1,05447 | -1,44214 | 23,29796 | 24,63478 | 25,24409 | 25,37951 | 26,36630 | 25,75743 |
| | Nucleoporin p54 | P46060 | NUP54 | 1,05306 | 2,55075 | 19,59578 | 19,30258 | 18,53321 | 18,77931 | 15,50000 | 15,50000 |
| | Alpha-centractin | Q5TA45 | ACTR1A | 1,04289 | 2,72313 | 20,48807 | 20,68748 | 20,82625 | 15,50000 | 19,10288 | 19,22952 |
| | Actin-related protein 2/3 complex subunit 5 | P49368 | ARPC5 | 1,03615 | -4,53368 | 15,50000 | 15,50000 | 21,53217 | 22,62482 | 21,21869 | 22,28971 |
| | Centrosome-associated protein 350 | Q969V3 | CEP350 | 1,03033 | -3,01799 | 15,50000 | 19,48528 | 15,50000 | 20,38638 | 19,97980 | 19,17307 |
| | NEDD8-conjugating enzyme Ubc12 | P04844 | UBE2M | 1,02566 | 2,54012 | 20,33298 | 20,80664 | 19,09605 | 15,50000 | 18,95708 | 18,15823 |
| | cAMP-dependent protein kinase type II-alpha regulatory subunit | P49419 | PRKAR2A | 1,02135 | 3,03367 | 20,28794 | 20,21549 | 19,12581 | 15,50000 | 19,52822 | 15,50000 |
| | SLIT-ROBO Rho GTPase-activating protein 2 | P63104 | SRGAP2 | 1,01615 | -3,81820 | 15,50000 | 20,76134 | 15,50000 | 20,95006 | 21,43746 | 20,82842 |
| | Endoplasmin | P22695 | HSP90B1 | 1,01531 | 0,80577 | 23,39924 | 23,77394 | 23,74837 | 23,47729 | 22,26296 | 22,76399 |
| | Tubulin beta chain | Q9BRX2 | TUBB | 1,01233 | 0,89574 | 25,51187 | 26,66044 | 26,65990 | 25,19637 | 25,70234 | 25,24627 |

| Significant | Protein names | UniProtID | Gene names | -LOG(P-value) | Difference | CXCR4_1 | CXCR4_2 | CXCR4_3 | MOCK_1 | MOCK_2 | MOCK_3 |
|---|---|---|---|---|---|---|---|---|---|---|---|
| | INO80 complex subunit B | P07195 | INO80B;INO80B-WBP1 | 1,01109 | -2,52781 | 18,95586 | 15,50000 | 18,92202 | 20,75735 | 20,31130 | 19,89266 |
| | Serine/threonine-protein phosphatase 6 regulatory subunit 3 | Q9UJZ1 | PPP6R3 | 1,00864 | 2,51266 | 20,50707 | 20,56113 | 19,80928 | 19,07801 | 18,76150 | 15,50000 |
| | Protein FAM49B | Q16531 | FAM49B | 1,00736 | 2,93305 | 21,52974 | 21,34987 | 20,26831 | 15,50000 | 19,42045 | 19,42833 |
| | U2 small nuclear ribonucleoprotein A | Q8WWY3 | SNRPA1 | 1,00681 | -1,00870 | 20,83838 | 21,99217 | 22,39561 | 22,72875 | 22,66475 | 22,85876 |
| | 26S proteasome non-ATPase regulatory subunit 11 | Q9BRP1 | PSMD11 | 0,99734 | 3,50370 | 22,39430 | 22,24002 | 22,24548 | 15,50000 | 20,28839 | 20,58031 |
| | Heat shock protein 75 kDa, mitochondrial | Q96FW1 | TRAP1 | 0,99718 | 1,20226 | 23,48554 | 24,94226 | 24,29172 | 22,42111 | 23,72234 | 22,96928 |
| | Dolichyl-diphosphooligosaccharide--protein glycosyltransferase 48 kDa subunit | Q15024 | DDOST | 0,99405 | 1,47189 | 22,62464 | 22,93659 | 23,82075 | 20,46991 | 22,16159 | 22,33481 |
| | Vacuolar protein sorting-associated protein 26A | Q9Y312 | VPS26A | 0,99070 | 2,81204 | 20,82361 | 21,20335 | 20,78788 | 15,50000 | 19,73655 | 19,14218 |
| | Glutamate dehydrogenase 1, mitochondrial;Glutamate dehydrogenase 2, mitochondrial | Q96TA2 | GLUD1;GLUD2 | 0,98977 | 2,59615 | 19,39276 | 19,56750 | 18,98067 | 15,50000 | 19,15249 | 15,50000 |
| | Eukaryotic initiation factor 4A-I | Q14165 | EIF4A1 | 0,98932 | 0,83553 | 26,94257 | 27,02781 | 27,48762 | 25,62709 | 26,82748 | 26,49683 |
| | Unconventional myosin-Ic | Q14997 | MYO1C | 0,98766 | 0,43292 | 23,65818 | 23,28912 | 23,25494 | 22,86891 | 23,28065 | 22,75391 |
| | DNA replication licensing factor MCM2 | P11177 | MCM2 | 0,98565 | 3,68391 | 21,68624 | 20,17767 | 20,77692 | 15,50000 | 15,50000 | 20,58912 |
| | Apoptotic chromatin condensation inducer in the nucleus | P62979 | ACIN1 | 0,98410 | -1,69579 | 21,20114 | 21,63702 | 23,30103 | 22,81055 | 23,92796 | 24,48806 |
| | Mitochondrial import inner membrane translocase subunit TIM50 | O14893 | TIMM50 | 0,97915 | 1,06547 | 21,51624 | 22,18513 | 21,16681 | 20,42378 | 21,33047 | 19,91752 |
| | Sorting nexin-27 | Q9UPQ0 | SNX27 | 0,97250 | 2,07777 | 18,94829 | 18,47366 | 18,25012 | 15,50000 | 15,50000 | 18,43876 |
| | Erlin-2 | P54577 | ERLIN2 | 0,97238 | 1,91191 | 23,65163 | 23,39741 | 23,11261 | 20,91532 | 23,25124 | 20,25937 |
| | Mediator of RNA polymerase II transcription subunit 7 | Q9NUL7 | MED7 | 0,97111 | -3,81793 | 15,50000 | 20,82889 | 15,50000 | 21,95539 | 20,26979 | 21,05748 |
| | Serine palmitoyltransferase 2 | Q9Y2L1 | SPTLC2 | 0,96908 | 2,56034 | 20,33604 | 20,67435 | 20,46364 | 15,50000 | 19,51198 | 18,78101 |
| | Proteasome subunit alpha type-6 | P39656 | PSMA6 | 0,96790 | 2,71416 | 20,33429 | 20,50911 | 21,41473 | 15,50000 | 19,22870 | 19,38694 |
| | Squalene synthase | P04406 | FDFT1 | 0,95445 | 2,88726 | 21,40062 | 21,34857 | 20,83738 | 15,50000 | 19,65672 | 19,76807 |
| | U3 small nucleolar RNA-associated protein 14 homolog A | O75616 | UTP14A | 0,95418 | -0,61885 | 20,08213 | 20,20836 | 20,70235 | 21,20389 | 21,17005 | 20,47546 |
| | NADH dehydrogenase [ubiquinone] 1 alpha subcomplex subunit 12 | Q29RF7 | NDUFA12 | 0,95105 | 2,53228 | 20,89984 | 20,30697 | 19,88948 | 15,50000 | 19,56326 | 18,43620 |
| | Dihydrolipoyllysine-residue succinyltransferase component of 2-oxoglutarate dehydrogenase complex, mitochondrial | O43175 | DLST | 0,94962 | 2,55205 | 19,03746 | 19,59062 | 19,27004 | 15,50000 | 19,24198 | 15,50000 |
| | Caspase activity and apoptosis inhibitor 1 | Q5JWF2 | CAAP1 | 0,94780 | -1,19391 | 20,08434 | 21,36874 | 20,47229 | 21,44662 | 22,73358 | 21,32691 |
| | Sorting nexin-5 | P07947 | SNX5 | 0,94546 | 0,61496 | 20,07822 | 19,54248 | 19,70207 | 19,65823 | 18,78561 | 19,03405 |
| | DnaJ homolog subfamily B member 12 | Q6KC79 | DNAJB12 | 0,94445 | 0,89747 | 22,04487 | 21,77205 | 20,89571 | 21,07019 | 20,13696 | 20,81308 |
| | Etoposide-induced protein 2.4 homolog | Q8TDX7 | EI24 | 0,94262 | 2,18984 | 18,84046 | 19,14736 | 18,24289 | 15,50000 | 18,66119 | 15,50000 |
| | Sodium/potassium-transporting ATPase subunit beta-3 | Q96IJ6 | ATP1B3 | 0,94233 | 3,08993 | 21,40831 | 21,53284 | 21,97316 | 15,50000 | 20,10072 | 20,04380 |
| | Putative RNA-binding protein Luc7-like 2 | Q96BW9 | LUC7L2 | 0,94187 | -0,79698 | 22,79654 | 22,42424 | 23,22233 | 23,20364 | 23,38416 | 24,24627 |
| | Ras-related protein Rap-1A | Q99816 | RAP1A | 0,94109 | 3,10393 | 22,10127 | 21,91262 | 20,62355 | 15,50000 | 19,95177 | 19,87388 |
| | Apoptosis-inducing factor 1, mitochondrial | P12004 | AIFM1 | 0,94048 | 0,97043 | 22,95886 | 24,15954 | 23,46750 | 23,22691 | 22,23474 | 22,21297 |
| | Replication factor C subunit 2 | P68363 | RFC2 | 0,93987 | -0,72142 | 23,48001 | 23,77886 | 22,66629 | 24,20202 | 24,12487 | 23,76253 |
| | Cytochrome c oxidase subunit 2 | O75586 | MT-CO2 | 0,93667 | 3,58405 | 23,36786 | 21,19886 | 22,61594 | 15,50000 | 20,04953 | 20,88099 |
| | Coatomer subunit zeta-1 | Q7Z417 | COPZ1 | 0,93303 | 2,86986 | 21,10553 | 21,82749 | 20,21869 | 15,50000 | 19,11614 | 19,92599 |
| | Sarcoplasmic/endoplasmic reticulum calcium ATPase 2 | Q92598 | ATP2A2 | 0,93076 | 3,95452 | 23,80577 | 23,52886 | 22,81047 | 15,50000 | 21,42511 | 21,35644 |
| | 14-3-3 protein eta | P27348 | YWHAH | 0,93023 | 1,27844 | 21,89589 | 21,73554 | 20,61450 | 21,02952 | 19,29948 | 20,08161 |
| | 14-3-3 protein zeta/delta | P62258 | YWHAZ | 0,92949 | 1,60460 | 24,23856 | 24,14996 | 22,94204 | 23,54038 | 21,33129 | 21,64509 |
| | Probable ATP-dependent RNA helicase DDX47 | P40227 | DDX47 | 0,92937 | -1,40245 | 22,71689 | 23,95234 | 24,74719 | 24,63954 | 25,04118 | 25,94304 |
| | Alpha-enolase | Q96DG6 | ENO1 | 0,92921 | 1,01878 | 22,98784 | 22,59462 | 22,76124 | 22,67282 | 20,95056 | 21,66397 |
| | Cytochrome b-c1 complex subunit 1, mitochondrial | P27708 | UQCRC1 | 0,92762 | 2,67374 | 21,27986 | 20,88017 | 20,17219 | 15,50000 | 19,19460 | 19,61641 |
| | 28S ribosomal protein S18b, mitochondrial | P05455 | MRPS18B | 0,92349 | -0,75473 | 22,06714 | 22,45229 | 22,22618 | 22,68357 | 23,73169 | 22,59455 |
| | Microsomal glutathione S-transferase 3 | P16298 | MGST3 | 0,92181 | -4,19051 | 15,50000 | 15,50000 | 21,16429 | 20,56291 | 23,52229 | 20,65062 |
| | Transmembrane emp24 domain-containing protein 1 | Q96KA5 | TMED1 | 0,92069 | 2,36386 | 20,63005 | 20,12439 | 19,90905 | 15,50000 | 19,13856 | 18,93336 |
| | 28S ribosomal protein S27, mitochondrial | P49674 | MRPS27 | 0,91534 | -1,15107 | 22,71293 | 22,80176 | 23,28799 | 23,73335 | 25,18090 | 23,34164 |
| | 28S ribosomal protein S28, mitochondrial | Q15006 | MRPS28 | 0,91507 | -3,95233 | 15,50000 | 15,50000 | 21,18749 | 20,88301 | 22,70233 | 20,45914 |
| | Bromodomain-containing protein 7 | Q9BXS5 | BRD7 | 0,91458 | -2,85214 | 15,50000 | 19,64195 | 19,57052 | 21,33014 | 21,80369 | 20,13508 |
| | Importin subunit alpha-4 | P02786 | KPNA3 | 0,91418 | 2,60566 | 20,77796 | 20,87260 | 20,64130 | 15,50000 | 19,48196 | 19,49293 |
| | Acetolactate synthase-like protein | P78371 | ILVBL | 0,91207 | 2,81308 | 20,55422 | 21,58215 | 21,23596 | 15,50000 | 19,85813 | 19,57496 |
| | Replication factor C subunit 3 | Q99832 | RFC3 | 0,91088 | -0,51039 | 21,63143 | 22,40346 | 22,11911 | 22,71643 | 22,67107 | 22,29768 |
| | Actin-related protein 2/3 complex subunit 4 | Q04917 | ARPC4;ARPC4-TTLL3 | 0,90788 | -0,64804 | 23,64317 | 23,53777 | 24,20613 | 24,32512 | 24,06340 | 24,94266 |
| | Galactokinase | P09211 | GALK1 | 0,90784 | 3,01209 | 21,74546 | 22,10239 | 20,44647 | 20,08408 | 19,67399 | 15,50000 |
| | E3 SUMO-protein ligase RanBP2 | P06493 | RANBP2 | 0,90587 | 3,03077 | 21,96409 | 21,58751 | 20,97582 | 20,60856 | 19,32656 | 15,50000 |
| | Helicase-like transcription factor | O00629 | HLTF | 0,90400 | -0,62796 | 24,22037 | 24,99036 | 25,07778 | 25,14927 | 25,73242 | 25,29070 |
| | Peroxiredoxin-6 | Q8NDV7 | PRDX6 | 0,90367 | 0,71424 | 22,11178 | 22,30811 | 21,96095 | 21,94263 | 20,73901 | 21,55647 |
| | Pre-mRNA-processing factor 19 | P55072 | PRPF19 | 0,90359 | -1,51392 | 23,50494 | 24,10613 | 24,98746 | 24,74540 | 26,95538 | 25,43952 |
| | Integrator complex subunit 11 | Q14204 | CPSF3L | 0,90233 | 1,62727 | 18,70944 | 19,03566 | 18,58432 | 15,50000 | 17,80247 | 18,14515 |
| | Double-stranded RNA-specific adenosine deaminase | P07900 | ADAR | 0,89958 | 0,58545 | 21,58642 | 21,39822 | 22,03574 | 20,76490 | 20,94814 | 21,55098 |
| | 40S ribosomal protein S17 | Q99536 | RPS17 | 0,89798 | -1,70949 | 24,68325 | 24,21934 | 25,90777 | 26,34723 | 28,03549 | 25,55611 |
| | Mitochondrial ribonuclease P protein 1 | Q8WVM0 | TRMT10C | 0,89456 | -1,26872 | 23,11710 | 22,82075 | 22,80676 | 23,09010 | 25,35030 | 24,11036 |
| | Desmoplakin | P08238 | DSP | 0,89417 | 3,07622 | 22,16886 | 21,49390 | 21,55258 | 15,50000 | 19,96169 | 20,52497 |
| | Importin subunit alpha-5;Importin subunit alpha-5, N-terminally processed | P52294 | KPNA1 | 0,89334 | 1,22064 | 20,98669 | 20,94092 | 19,90708 | 20,43634 | 18,71011 | 19,02632 |

| Significant | Protein names | UniProtID | Gene names | -LOG(P-value) | Difference | CXCR4_1 | CXCR4_2 | CXCR4_3 | MOCK_1 | MOCK_2 | MOCK_3 |
|---|---|---|---|---|---|---|---|---|---|---|---|
| | Retinol dehydrogenase 11 | O14654 | RDH11 | 0,89291 | 2,94662 | 21,82784 | 21,88234 | 19,97757 | 15,50000 | 20,01699 | 19,33090 |
| | Mitochondrial glutamate carrier 1 | P23258 | SLC25A22 | 0,88913 | 2,89629 | 21,73645 | 21,39503 | 21,03436 | 15,50000 | 19,71302 | 20,26396 |
| | Vesicle-associated membrane protein-associated protein B/C | Q12931 | VAPB | 0,88767 | 3,10063 | 21,42634 | 22,28991 | 21,65036 | 15,50000 | 19,92924 | 20,63547 |
| | Probable ATP-dependent RNA helicase DHX37 | P49770 | DHX37 | 0,88571 | -1,52033 | 19,93272 | 20,32202 | 21,60404 | 21,67633 | 23,36826 | 21,37518 |
| | Alkyldihydroxyacetonephosphate synthase, peroxisomal | Q9NNW5 | AGPS | 0,88547 | 0,61555 | 21,03382 | 21,88446 | 21,69447 | 21,01624 | 21,20467 | 20,54519 |
| | 26S protease regulatory subunit 10B | Q9H0U3 | PSMC6 | 0,88367 | 2,89669 | 20,61144 | 21,55964 | 21,86007 | 15,50000 | 20,16116 | 19,67993 |
| | Galactosylgalactosylxylosylprotein 3-beta-glucuronosyltransferase 3 | O14929 | B3GAT3 | 0,88131 | -1,32574 | 21,80845 | 23,18089 | 22,23228 | 23,22112 | 24,87343 | 23,10428 |
| | Axin interactor, dorsalization-associated protein | P43307 | AIDA | 0,88108 | -0,72377 | 20,24237 | 20,85650 | 20,11959 | 21,56987 | 21,28161 | 20,53830 |
| | Splicing factor 3B subunit 6 | P68371 | SF3B6 | 0,88018 | -1,35623 | 21,89201 | 22,64048 | 23,41596 | 22,87892 | 24,67896 | 24,45926 |
| | 26S proteasome non-ATPase regulatory subunit 2 | P17987 | PSMD2 | 0,87929 | 3,61404 | 22,02938 | 23,03381 | 23,46973 | 15,50000 | 21,19496 | 20,99583 |
| | pre-rRNA processing protein FTSJ3 | Q15645 | FTSJ3 | 0,87651 | -4,20564 | 15,50000 | 15,50000 | 21,85631 | 20,53886 | 22,98245 | 21,95191 |
| | Transcription initiation factor TFIID subunit 2 | P52701 | TAF2 | 0,87600 | -1,39851 | 19,94092 | 20,27128 | 21,34077 | 20,81706 | 22,93043 | 22,00100 |
| | Protein lunapark | O43242 | LNP | 0,87582 | 3,10481 | 20,81964 | 20,06763 | 19,79211 | 15,50000 | 20,36496 | 15,50000 |
| | Alpha-galactosidase A | P50991 | GLA | 0,87015 | 2,54812 | 20,22952 | 20,39864 | 21,37210 | 15,50000 | 19,20936 | 19,64653 |
| | Thioredoxin-dependent peroxide reductase, mitochondrial | P08559 | PRDX3 | 0,86953 | -2,16941 | 15,50000 | 15,50000 | 18,81334 | 19,38064 | 18,14201 | 18,79893 |
| | Alpha-globin transcription factor CP2 | Q15084 | TFCP2 | 0,86683 | 2,84204 | 18,32672 | 20,41359 | 20,36922 | 15,50000 | 15,50000 | 19,58342 |
| | Ubiquitin carboxyl-terminal hydrolase 16 | P04181 | USP16 | 0,86266 | -1,24342 | 22,42573 | 23,13684 | 22,97993 | 23,66384 | 25,33964 | 23,26928 |
| | 28S ribosomal protein S10, mitochondrial | Q9H9B4 | MRPS10 | 0,85992 | -3,52198 | 15,50000 | 15,50000 | 20,31562 | 19,44908 | 22,66800 | 19,76448 |
| | G patch domain-containing protein 4 | P31946 | GPATCH4 | 0,85661 | -1,17039 | 21,07725 | 22,87963 | 22,79992 | 23,90205 | 23,20401 | 23,16189 |
| | AP-2 complex subunit sigma | Q15392 | AP2S1 | 0,85634 | -3,00786 | 15,50000 | 19,85518 | 20,50766 | 20,95886 | 22,46129 | 21,46628 |
| | Copine-3 | P55010 | CPNE3 | 0,85589 | -0,63321 | 23,43466 | 23,41506 | 24,16630 | 23,82900 | 24,57900 | 24,50767 |
| | Tubulin beta-6 chain | Q9BSD7 | TUBB6 | 0,85429 | 3,03474 | 22,14613 | 21,81652 | 21,41488 | 20,67219 | 20,10111 | 15,50000 |
| | Protein PRRC2C | Q9BVA1 | PRRC2C | 0,85234 | 0,58481 | 20,57311 | 20,22506 | 21,26419 | 20,10137 | 19,94478 | 20,26178 |
| | Pyruvate dehydrogenase E1 component subunit alpha, somatic form, mitochondrial | Q9UMR2 | PDHA1 | 0,85224 | 1,14121 | 21,95748 | 22,79612 | 22,97841 | 20,36997 | 21,85859 | 22,07982 |
| | 60S ribosomal protein L22 | Q5VT25 | RPL22 | 0,84889 | -0,89990 | 27,83237 | 29,02681 | 28,28771 | 29,96568 | 29,05709 | 28,82384 |
| | Succinyl-CoA ligase [GDP-forming] subunit beta, mitochondrial | O60684 | SUCLG2 | 0,84865 | 2,03616 | 20,19748 | 19,31591 | 19,57845 | 15,50000 | 18,87541 | 18,60794 |
| | NADH dehydrogenase [ubiquinone] 1 alpha subcomplex subunit 10, mitochondrial | Q04637 | NDUFA10 | 0,84318 | 0,87148 | 21,17133 | 21,87804 | 20,82423 | 21,15097 | 20,08874 | 20,01944 |
| | Vacuolar protein sorting-associated protein 4B | P11908 | VPS4B | 0,84206 | 2,47681 | 20,15289 | 20,91240 | 20,91086 | 15,50000 | 19,59962 | 19,44609 |
| | OCIA domain-containing protein 1 | Q3ZCQ8 | OCIAD1 | 0,84070 | 0,48538 | 20,59222 | 20,00470 | 20,17316 | 20,11782 | 19,78200 | 19,41413 |
| | Heterogeneous nuclear ribonucleoprotein H2 | Q9HCK8 | HNRNPH2 | 0,84047 | -0,44356 | 22,71773 | 23,24909 | 22,78087 | 23,19945 | 23,71607 | 23,16285 |
| | 14-3-3 protein epsilon | O15027 | YWHAE | 0,83770 | 1,36264 | 24,53825 | 24,56197 | 23,79907 | 24,35528 | 22,11712 | 22,33898 |
| | Guanine nucleotide-binding protein-like 3 | Q13263 | GNL3 | 0,83374 | -1,38582 | 22,27099 | 24,53991 | 24,59486 | 25,35394 | 25,04431 | 25,16496 |
| | Transcription initiation factor TFIID subunit 8 | Q13347 | TAF8 | 0,83331 | -1,81297 | 15,50000 | 18,49779 | 18,15590 | 19,02380 | 19,86644 | 18,70235 |
| | Ribosome biogenesis protein BOP1 | Q9P2R7 | BOP1 | 0,83120 | -3,10001 | 15,50000 | 19,22256 | 21,22158 | 20,90478 | 22,25205 | 22,08735 |
| | Chromatin assembly factor 1 subunit B | P06733 | CHAF1B | 0,83001 | -3,27237 | 15,50000 | 21,08537 | 20,78069 | 22,31116 | 22,83082 | 22,04120 |
| | DnaJ homolog subfamily C member 1 | Q9NRG9 | DNAJC1 | 0,82531 | -0,90980 | 19,82775 | 20,79408 | 20,16607 | 21,35208 | 21,80251 | 20,36271 |
| | Heterogeneous nuclear ribonucleoprotein H;Heterogeneous nuclear ribonucleoprotein H, N-terminally processed | P00338 | HNRNPH1 | 0,82504 | -0,62155 | 25,74940 | 26,68164 | 26,31786 | 26,45565 | 27,20284 | 26,95505 |
| | Guanine nucleotide-binding protein G(I)/G(S)/G(T) subunit beta-2 | P20340 | GNB2 | 0,81871 | 2,71210 | 21,67728 | 20,86106 | 21,19003 | 15,50000 | 19,99493 | 20,09713 |
| | THUMP domain-containing protein 3 | O60701 | THUMPD3 | 0,81643 | 0,58354 | 18,96874 | 19,83618 | 19,66920 | 19,26199 | 18,58038 | 18,88113 |
| | Programmed cell death protein 4 | O75306 | PDCD4 | 0,81563 | 1,78684 | 19,68526 | 19,16322 | 18,95487 | 15,50000 | 18,43925 | 18,50359 |
| | Kelch domain-containing protein 4 | P55735 | KLHDC4 | 0,81151 | -2,27943 | 15,50000 | 19,15021 | 19,54785 | 20,63174 | 20,03241 | 20,37220 |
| | 28S ribosomal protein S7, mitochondrial | Q86U42 | MRPS7 | 0,81109 | -1,30848 | 20,65176 | 21,46399 | 22,52095 | 22,12521 | 23,84680 | 22,59012 |
| | 28S ribosomal protein S25, mitochondrial | P55884 | MRPS25 | 0,80983 | -3,49713 | 15,50000 | 15,50000 | 20,82205 | 19,71664 | 22,60200 | 19,99479 |
| | Isoleucine--tRNA ligase, mitochondrial | P33991 | IARS2 | 0,80883 | 1,94890 | 20,16337 | 19,52200 | 19,02388 | 15,50000 | 18,51545 | 18,84711 |
| | L-lactate dehydrogenase B chain | P25705 | LDHB | 0,80748 | 1,59718 | 24,84910 | 23,13214 | 22,87350 | 23,24632 | 20,92665 | 21,89023 |
| | Actin-related protein 5 | Q9NPF5 | ACTR5 | 0,80649 | -2,60392 | 15,50000 | 19,99921 | 19,86843 | 21,36228 | 21,16011 | 20,65701 |
| | Tubulin alpha-1C chain | O95831 | TUBA1C | 0,80633 | 3,09459 | 21,86718 | 22,38279 | 22,14171 | 15,50000 | 20,88800 | 20,71992 |
| | Biliverdin reductase A | P48426 | BLVRA | 0,80530 | 2,80645 | 21,19406 | 21,93012 | 21,22623 | 15,50000 | 20,66943 | 19,76163 |
| | NADH dehydrogenase [ubiquinone] iron-sulfur protein 2, mitochondrial | Q9Y266 | NDUFS2 | 0,80403 | 0,99396 | 21,68765 | 22,46099 | 22,95849 | 20,72442 | 22,20288 | 21,19797 |
| | Ubiquitin-like modifier-activating enzyme 1 | Q9BQ75 | UBA1 | 0,80387 | 2,44357 | 21,47274 | 19,82766 | 20,42511 | 19,64294 | 15,50000 | 19,25186 |
| | DnaJ homolog subfamily C member 13 | Q15738 | DNAJC13 | 0,80253 | 3,04389 | 22,47951 | 21,64258 | 21,37544 | 15,50000 | 21,31601 | 19,54984 |
| | NADH-cytochrome b5 reductase 3;NADH-cytochrome b5 reductase 3 membrane-bound form;NADH-cytochrome b5 reductase 3 soluble form | Q9NVI1 | CYB5R3 | 0,80242 | 2,67876 | 21,80644 | 20,92412 | 20,75743 | 15,50000 | 19,58954 | 20,36217 |
| | Coatomer subunit gamma-1 | P25789 | COPG1 | 0,80238 | 3,54032 | 23,06273 | 22,98694 | 23,22010 | 15,50000 | 21,98318 | 21,16564 |
| | 28S ribosomal protein S5, mitochondrial | Q9UKX7 | MRPS5 | 0,80185 | -3,38058 | 15,50000 | 19,96226 | 21,19442 | 21,95052 | 23,95456 | 20,89334 |
| | Methylcrotonoyl-CoA carboxylase beta chain, mitochondrial | Q6YN16 | MCCC2 | 0,79934 | 0,83716 | 21,25805 | 21,07712 | 20,60585 | 21,02141 | 19,80837 | 19,59976 |
| | Proline-serine-threonine phosphatase-interacting protein 2 | Q9P289 | PSTPIP2 | 0,79779 | -2,20190 | 15,50000 | 19,60805 | 17,93261 | 20,25026 | 20,41225 | 18,98384 |
| | Pre-mRNA-splicing factor SPF27 | Q13148 | BCAS2 | 0,79719 | -3,70578 | 20,90507 | 15,50000 | 22,47894 | 24,00513 | 23,31368 | 22,68254 |
| | Succinyl-CoA ligase [ADP-forming] subunit beta, mitochondrial | Q9UPQ9 | SUCLA2 | 0,79538 | 1,02031 | 21,33162 | 21,01910 | 20,12187 | 20,71867 | 19,52340 | 19,16957 |
| | A-kinase anchor protein 13 | O43823 | AKAP13 | 0,79208 | -3,11292 | 19,51782 | 15,50000 | 21,09944 | 21,14768 | 23,25322 | 21,05510 |
| | Ras-related protein Rab-5A | P56545 | RAB5A | 0,78812 | -2,81873 | 15,50000 | 15,50000 | 20,25315 | 20,74927 | 19,15469 | 19,80538 |
| | N-alpha-acetyltransferase 40 | P17812 | NAA40 | 0,78772 | -2,71736 | 15,50000 | 18,83334 | 19,86735 | 19,57424 | 22,52781 | 20,25072 |
| | Protein transport protein Sec16A | P32119 | SEC16A | 0,78638 | 1,04013 | 21,95947 | 21,34587 | 20,62658 | 21,21342 | 19,87368 | 19,72444 |

| Significant | Protein names | UniProtID | Gene names | -LOG(P-value) | Difference | CXCR4_1 | CXCR4_2 | CXCR4_3 | MOCK_1 | MOCK_2 | MOCK_3 |
|---|---|---|---|---|---|---|---|---|---|---|---|
| | Nucleolar complex protein 3 homolog | Q8NBI6 | NOC3L | 0,78513 | -1,45360 | 22,83080 | 23,40847 | 25,18529 | 24,30814 | 25,70838 | 25,76883 |
| | Guanine nucleotide-binding protein G(k) subunit alpha | P60981 | GNAI3 | 0,78390 | 2,40370 | 20,88524 | 21,12218 | 19,85740 | 15,50000 | 19,47827 | 19,67547 |
| | FAS-associated factor 2 | P60891 | FAF2 | 0,78386 | 3,32547 | 22,66051 | 22,90599 | 22,39874 | 15,50000 | 21,89671 | 20,59213 |
| | COP9 signalosome complex subunit 4 | P61981 | COPS4 | 0,78383 | 1,83633 | 19,82505 | 19,26027 | 19,28209 | 15,50000 | 18,85830 | 18,50013 |
| | 60S ribosomal protein L28 | Q99497 | RPL28 | 0,78218 | -3,00515 | 20,41132 | 20,82151 | 15,50000 | 21,04840 | 22,64384 | 22,05603 |
| | Protein FAM98A | P55209 | FAM98A | 0,77994 | -0,21407 | 23,67465 | 23,61885 | 23,55836 | 24,01546 | 23,87802 | 23,60060 |
| | U4/U6 small nuclear ribonucleoprotein Prp4 | Q9NXW2 | PRPF4 | 0,77923 | -1,03139 | 21,04427 | 21,21981 | 22,26851 | 21,60368 | 23,14636 | 22,87671 |
| | Y-box-binding protein 3 | Q9UHI6 | YBX3 | 0,77529 | -0,87538 | 21,57358 | 21,77901 | 23,21355 | 23,11300 | 23,14940 | 22,92985 |
| | Far upstream element-binding protein 3 | P07437 | FUBP3 | 0,77504 | 0,83177 | 22,79285 | 22,93168 | 22,35668 | 22,66090 | 21,05643 | 21,86857 |
| | Activator of 90 kDa heat shock protein ATPase homolog 1 | P18085 | AHSA1 | 0,77474 | 3,41421 | 23,13187 | 22,95131 | 22,74335 | 15,50000 | 21,94642 | 21,13746 |
| | Chromodomain-helicase-DNA-binding protein 8 | O95299 | CHD8 | 0,77459 | 1,04909 | 22,55733 | 20,64773 | 20,91532 | 20,31915 | 20,64201 | 20,01195 |
| | 40S ribosomal protein S15a | Q68CQ4 | RPS15A | 0,77276 | -1,34254 | 22,89165 | 23,36880 | 24,88879 | 24,26707 | 26,05778 | 24,85201 |
| | Calnexin | Q9UJS0 | CANX | 0,77057 | 3,63321 | 22,04453 | 23,68014 | 24,13328 | 15,50000 | 21,66757 | 21,79074 |
| | N-acylneuraminate cytidylyltransferase | Q99613 | CMAS | 0,76987 | -0,88273 | 20,15968 | 20,77628 | 21,23567 | 21,48557 | 20,93647 | 22,39778 |
| | Elongator complex protein 1 | P52292 | IKBKAP | 0,76811 | 2,47857 | 21,10169 | 20,77989 | 20,87170 | 20,24725 | 19,57032 | 15,50000 |
| | Poly(A) RNA polymerase, mitochondrial | P48444 | MTPAP | 0,76646 | -0,35181 | 21,59755 | 21,43492 | 21,59391 | 21,57015 | 22,27185 | 21,83981 |
| | Ubiquitin carboxyl-terminal hydrolase 7 | Q9HCC0 | USP7 | 0,76565 | 2,27021 | 20,53127 | 20,34830 | 20,55412 | 15,50000 | 19,28658 | 19,83647 |
| | Splicing factor 3B subunit 1 | P60842 | SF3B1 | 0,76370 | -0,73452 | 26,42865 | 27,23644 | 27,74855 | 27,45568 | 27,96036 | 28,20116 |
| | Mitotic checkpoint protein BUB3 | P49591 | BUB3 | 0,76359 | -0,85707 | 23,60048 | 23,90435 | 23,57808 | 24,92506 | 23,55075 | 25,17832 |
| | Mothers against decapentaplegic homolog 2 | Q96I24 | SMAD2 | 0,76230 | -0,68753 | 18,98251 | 19,48129 | 19,12556 | 19,60473 | 20,64974 | 19,39747 |
| | AP-2 complex subunit alpha-1 | Q9UNQ2 | AP2A1 | 0,76061 | -1,09291 | 23,33906 | 24,24365 | 24,54575 | 24,08002 | 25,94673 | 25,38044 |
| | Guanine nucleotide-binding protein G(I)/G(S)/G(T) subunit beta-1 | P14625 | GNB1 | 0,75860 | 3,14185 | 22,40629 | 22,04930 | 22,75071 | 15,50000 | 20,74895 | 21,53179 |
| | Nuclear export mediator factor NEMF | Q6P4A7 | NEMF | 0,75750 | -0,79761 | 22,93086 | 22,66295 | 23,53017 | 23,28136 | 24,63998 | 23,59548 |
| | Elongation factor Tu, mitochondrial | P07237 | TUFM | 0,75584 | 0,39332 | 25,34935 | 25,31824 | 25,62745 | 25,04718 | 25,41158 | 24,65633 |
| | Eukaryotic initiation factor 4A-III;Eukaryotic initiation factor 4A-III, N-terminally processed | Q99733 | EIF4A3 | 0,75299 | -0,99110 | 24,95109 | 24,70546 | 25,18052 | 24,82760 | 26,14767 | 26,83509 |
| | H/ACA ribonucleoprotein complex subunit 4 | O95714 | DKC1 | 0,75056 | -1,40469 | 21,64394 | 24,25465 | 23,55157 | 25,15869 | 23,90830 | 24,59725 |
| | Sterol-4-alpha-carboxylate 3-dehydrogenase, decarboxylating | Q92616 | NSDHL | 0,75008 | 0,95672 | 21,46289 | 22,27119 | 21,98701 | 22,02208 | 20,41349 | 20,41535 |
| | Transcriptional repressor protein YY1 | Q9NWY4 | YY1 | 0,74931 | -1,30845 | 15,50000 | 17,34776 | 18,07828 | 18,67349 | 17,86605 | 18,31186 |
| | Ribosomal L1 domain-containing protein 1 | Q9Y277 | RSL1D1 | 0,74771 | -2,97502 | 15,50000 | 15,50000 | 20,79241 | 19,55150 | 20,01550 | 21,15047 |
| | Dihydrolipoyl dehydrogenase, mitochondrial | Q7L2H7 | DLD | 0,74607 | 1,98323 | 20,53905 | 19,52424 | 19,02753 | 19,26598 | 18,37518 | 15,50000 |
| | NADH dehydrogenase [ubiquinone] 1 beta subcomplex subunit 10 | Q15437 | NDUFB10 | 0,74467 | 2,44922 | 20,54793 | 21,65110 | 19,99479 | 15,50000 | 20,23911 | 19,10705 |
| | Xyloside xylosyltransferase 1 | P55036 | XXYLT1 | 0,74301 | 0,91264 | 21,89367 | 20,88196 | 20,17950 | 20,60141 | 19,78389 | 19,83192 |
| | General transcription factor IIH subunit 4 | Q14152 | GTF2H4 | 0,74255 | -1,44300 | 18,82547 | 19,68507 | 20,38911 | 19,94921 | 22,54489 | 20,73455 |
| | Succinate dehydrogenase [ubiquinone] iron-sulfur subunit, mitochondrial | P14618 | SDHB | 0,74092 | 2,25038 | 20,06842 | 20,83784 | 20,57976 | 15,50000 | 19,47799 | 19,75690 |
| | 40S ribosomal protein S25 | Q9NUL3 | RPS25 | 0,73960 | -2,00297 | 23,19260 | 23,08963 | 26,19299 | 24,96120 | 27,42550 | 26,09744 |
| | Proteasome subunit alpha type-4 | Q14558 | PSMA4 | 0,73915 | 0,95585 | 21,43293 | 20,96600 | 20,12086 | 20,78413 | 19,50731 | 19,36080 |
| | Transmembrane protein 206 | Q9Y285 | TMEM206 | 0,73616 | -2,33869 | 15,50000 | 19,54146 | 19,50232 | 20,38374 | 21,56626 | 19,60984 |
| | Histone H4 | P30041 | HIST1H4A | 0,73504 | -2,12779 | 22,03440 | 21,43898 | 24,63112 | 23,39663 | 24,61768 | 26,47355 |
| | Cleavage and polyadenylation specificity factor subunit 5 | Q9Y5M8 | NUDT21 | 0,73474 | -3,43705 | 15,50000 | 15,50000 | 21,76975 | 20,70058 | 20,41039 | 21,96993 |
| | Condensin complex subunit 3 | Q9H0U4 | NCAPG | 0,73109 | 2,19572 | 20,40935 | 20,51036 | 20,31904 | 19,21212 | 19,93948 | 15,50000 |
| | Pre-mRNA-splicing factor RBM22 | P61619 | RBM22 | 0,72805 | -2,68275 | 15,50000 | 20,76450 | 20,27878 | 21,21561 | 21,73219 | 21,64372 |
| | 40S ribosomal protein S26;Putative 40S ribosomal protein S26-like 1 | O75955 | RPS26;RPS26P11 | 0,72791 | -1,47194 | 22,07081 | 20,84743 | 23,96270 | 23,56987 | 24,14593 | 23,58095 |
| | Transketolase | Q9UBM7 | TKT | 0,72644 | 2,29484 | 19,61450 | 19,47306 | 18,50839 | 19,71143 | 15,50000 | 15,50000 |
| | ATP-dependent RNA helicase DDX18 | P21796 | DDX18 | 0,72297 | -2,14672 | 20,21502 | 22,68468 | 24,39231 | 23,35703 | 25,08764 | 25,28750 |
| | Heterogeneous nuclear ribonucleoproteins C1/C2 | Q7Z406 | HNRNPC | 0,72274 | -1,10013 | 23,64680 | 24,80985 | 25,44174 | 24,83990 | 26,01180 | 26,34709 |
| | Eukaryotic translation initiation factor 4 gamma 2 | O95573 | EIF4G2 | 0,72202 | 2,70203 | 20,60946 | 21,83384 | 22,05454 | 15,50000 | 20,63041 | 20,26132 |
| | Transferrin receptor protein 1;Transferrin receptor protein 1, serum form | Q9BZF1 | TFRC | 0,72111 | 1,28645 | 22,78740 | 22,97267 | 23,27412 | 20,41875 | 21,56695 | 23,18915 |
| | THO complex subunit 7 homolog | Q15365 | THOC7 | 0,71991 | -1,29583 | 22,62199 | 23,05882 | 22,79642 | 23,04572 | 25,71626 | 23,60275 |
| | Lysophosphatidylcholine acyltransferase 1 | P62826 | LPCAT1 | 0,71810 | 2,03555 | 19,78392 | 20,26121 | 20,22400 | 15,50000 | 19,60735 | 19,05513 |
| | Protein LTV1 homolog | Q08J23 | LTV1 | 0,71686 | -2,22672 | 19,98092 | 24,45331 | 23,74263 | 25,25587 | 25,25198 | 24,34915 |
| | Protein phosphatase 1F | O43865 | PPM1F | 0,71603 | 2,13906 | 20,91481 | 20,35434 | 19,30748 | 19,52808 | 19,13137 | 15,50000 |
| | Protein transport protein Sec24C | P04075 | SEC24C | 0,71477 | 2,95571 | 21,16601 | 20,37380 | 19,24433 | 20,91700 | 15,50000 | 15,50000 |
| | 3-hydroxyacyl-CoA dehydrogenase type-2 | Q86X55 | HSD17B10 | 0,70639 | -0,83612 | 21,10910 | 20,17439 | 21,92897 | 21,57265 | 22,21913 | 21,92904 |
| | Mitochondrial 2-oxoglutarate/malate carrier protein | Q8WVX9 | SLC25A11 | 0,70379 | -0,48099 | 23,85500 | 24,28185 | 23,92705 | 24,68635 | 24,87291 | 23,94762 |
| | Ras GTPase-activating-like protein IQGAP2 | Q9C0C9 | IQGAP2 | 0,70104 | -1,18439 | 23,00146 | 23,89190 | 24,34238 | 24,13540 | 26,24440 | 24,40912 |
| | Spermidine synthase | P13489 | SRM | 0,69642 | 2,51176 | 21,82412 | 20,97197 | 20,94349 | 15,50000 | 20,42859 | 20,27571 |
| | Peptidyl-prolyl cis-trans isomerase-like 4 | Q06210 | PPIL4 | 0,69484 | -2,18259 | 15,50000 | 19,89645 | 19,64672 | 20,40769 | 20,77804 | 20,40520 |
| | Eukaryotic translation initiation factor 2 subunit 2 | Q5JSZ5 | EIF2S2 | 0,69379 | -1,45411 | 21,87837 | 23,33919 | 24,42892 | 23,52013 | 25,56929 | 24,91939 |
| | CTP synthase 1 | O15372 | CTPS1 | 0,69176 | 0,92642 | 22,04173 | 23,90894 | 23,10127 | 21,82481 | 22,65482 | 21,79305 |
| | Tumor susceptibility gene 101 protein | Q15149 | TSG101 | 0,69154 | 1,43292 | 19,27143 | 18,54939 | 18,36735 | 15,50000 | 18,04815 | 18,34126 |
| | DNA mismatch repair protein Msh2 | Q13769 | MSH2 | 0,69070 | 2,72915 | 20,43543 | 22,33336 | 22,02908 | 15,50000 | 20,94399 | 20,16644 |
| | Erythrocyte band 7 integral membrane protein | Q14145 | STOM | 0,68977 | 3,00176 | 22,93560 | 22,22367 | 21,92354 | 15,50000 | 21,86582 | 20,71171 |
| | Testis-specific Y-encoded-like protein 1 | O00116 | TSPYL1 | 0,68953 | 2,06298 | 19,87869 | 20,25857 | 20,54350 | 19,85719 | 19,13463 | 15,50000 |
| | Peroxiredoxin-2 | Q9Y5X3 | PRDX2 | 0,68885 | 0,91297 | 22,47103 | 22,95449 | 21,32548 | 22,00120 | 20,75809 | 21,25280 |

| Significant | Protein names | UniProtID | Gene names | -LOG(P-value) | Difference | CXCR4_1 | CXCR4_2 | CXCR4_3 | MOCK_1 | MOCK_2 | MOCK_3 |
|---|---|---|---|---|---|---|---|---|---|---|---|
| | Sideroflexin-4 | Q96GQ7 | SFXN4 | 0,68842 | 0,80152 | 21,84969 | 21,23929 | 20,73851 | 21,25690 | 20,35380 | 19,81223 |
| | Ribonuclease inhibitor | P12236 | RNH1 | 0,68685 | 0,67352 | 23,92180 | 23,35205 | 22,74846 | 23,23954 | 22,46635 | 22,29586 |
| | UPF0568 protein C14orf166 | Q9NX63 | C14orf166 | 0,68685 | -1,01344 | 21,67133 | 21,57727 | 23,37212 | 22,81580 | 22,96564 | 23,87961 |
| | Protein mago nashi homolog 2 | Q99741 | MAGOHB | 0,68665 | -3,73345 | 15,50000 | 22,82594 | 22,88935 | 23,49277 | 24,43530 | 24,48757 |
| | BAG family molecular chaperone regulator 2 | Q9NVH1 | BAG2 | 0,68270 | -2,50542 | 15,50000 | 20,49013 | 15,50000 | 19,66735 | 19,44053 | 19,89850 |
| | Tubulin gamma-1 chain;Tubulin gamma-2 chain | O14497 | TUBG1;TUBG2 | 0,68246 | 1,21658 | 22,70007 | 23,00088 | 23,79452 | 20,52468 | 23,02169 | 22,29933 |
| | General transcription factor IIH subunit 1 | P55265 | GTF2H1 | 0,68202 | -2,07342 | 15,50000 | 19,65641 | 19,35124 | 20,64192 | 20,53744 | 19,54855 |
| | 28S ribosomal protein S23, mitochondrial | Q9Y520 | MRPS23 | 0,68161 | -3,52243 | 15,50000 | 20,54585 | 22,87497 | 22,15974 | 24,92524 | 22,40312 |
| | Cirhin | Q9BV44 | CIRH1A | 0,68093 | -2,32501 | 15,50000 | 20,17621 | 20,02662 | 21,18362 | 20,43135 | 21,06289 |
| | Epoxide hydrolase 1 | P13797 | EPHX1 | 0,67994 | 2,11482 | 20,72301 | 19,93948 | 20,53924 | 15,50000 | 19,75690 | 19,60038 |
| | Histone H2A type 1-J;Histone H2A type 1-H;Histone H2A.J;Histone H2A type 1-C;Histone H2A type 3;Histone H2A type 1-D;Histone H2A type 1;Histone H2A type 1-B/E | Q16891 | HIST1H2AJ;HIST1H2AH;H2AFJ;HIST1H2AC;HIST3H2A;HIST1H2AD;HIST1H2AG;HIST1H2AB | 0,67937 | -1,50908 | 22,41972 | 23,94726 | 25,23652 | 24,91155 | 24,65775 | 26,56144 |
| | Putative adenosylhomocysteinase 2 | P23528 | AHCYL1 | 0,67782 | 0,68749 | 20,27116 | 19,29585 | 18,79490 | 19,07389 | 18,56257 | 18,66299 |
| | Splicing factor, suppressor of white-apricot homolog | Q969Z0 | SFSWAP | 0,67411 | -0,84511 | 21,73880 | 23,46886 | 23,40873 | 23,74776 | 23,61829 | 23,78566 |
| | Ribosome biogenesis protein BMS1 homolog | P46977 | BMS1 | 0,67102 | -2,67327 | 15,50000 | 20,23468 | 21,04853 | 20,76191 | 22,56739 | 21,47373 |
| | Mediator of RNA polymerase II transcription subunit 27 | Q03252 | MED27 | 0,66857 | -0,71960 | 21,24905 | 22,54376 | 21,73686 | 23,02467 | 21,97481 | 22,68900 |
| | Probable ATP-dependent RNA helicase DDX41 | Q99829 | DDX41 | 0,66689 | -3,52834 | 15,50000 | 21,59550 | 22,30263 | 21,53141 | 25,15252 | 23,29922 |
| | 40S ribosomal protein S23 | P13639 | RPS23 | 0,66592 | -3,88160 | 15,50000 | 21,51017 | 24,17333 | 23,00743 | 24,97609 | 24,84479 |
| | Protein transport protein Sec61 subunit alpha isoform 1 | P49841 | SEC61A1 | 0,66516 | 0,70877 | 21,98801 | 21,96236 | 20,71196 | 21,30697 | 20,70447 | 20,52459 |
| | Splicing factor 45 | Q8WWM7 | RBM17 | 0,66512 | -1,30540 | 24,06085 | 26,86502 | 26,34996 | 26,68378 | 27,05879 | 27,44947 |
| | RalBP1-associated Eps domain-containing protein 1 | P49821 | REPS1 | 0,66505 | -0,73791 | 20,93819 | 22,29025 | 22,45445 | 22,67109 | 22,86863 | 22,35689 |
| | Ubiquitin-associated domain-containing protein 2 | O00148 | UBAC2 | 0,66348 | 2,09439 | 20,65979 | 20,33985 | 20,31396 | 15,50000 | 19,56945 | 19,96099 |
| | Peptidyl-prolyl cis-trans isomerase FKBP8 | Q13557 | FKBP8 | 0,66273 | 2,41583 | 20,95929 | 21,23900 | 21,39645 | 15,50000 | 20,65866 | 20,18858 |
| | Chromodomain-helicase-DNA-binding protein 1-like | Q9NX40 | CHD1L | 0,66204 | -0,69567 | 22,38756 | 23,55321 | 23,90967 | 24,08318 | 23,73200 | 24,12226 |
| | Histone-binding protein RBBP7 | P35579 | RBBP7 | 0,66157 | -0,79819 | 22,81147 | 23,72796 | 23,89328 | 23,60637 | 25,08014 | 24,14079 |
| | Activator of basal transcription 1 | O00303 | ABT1 | 0,66041 | -0,69449 | 20,83792 | 21,01216 | 20,85795 | 22,40213 | 20,76393 | 21,62542 |
| | Pyruvate kinase PKM | P19387 | PKM | 0,65961 | 0,73351 | 25,42624 | 24,40957 | 24,35568 | 24,69024 | 23,84182 | 23,45889 |
| | p21-activated protein kinase-interacting protein 1 | Q15366 | PAK1IP1 | 0,65693 | -1,32907 | 21,06480 | 20,53754 | 22,03278 | 20,94271 | 23,18015 | 23,49948 |
| | 60S ribosomal protein L10a | O75821 | RPL10A | 0,65660 | -1,65641 | 22,32053 | 23,07795 | 25,85757 | 24,71167 | 25,47590 | 26,03771 |
| | Translation machinery-associated protein 16 | P12277 | TMA16 | 0,64964 | -1,16612 | 20,86869 | 20,58344 | 22,36172 | 21,24696 | 23,05142 | 23,01381 |
| | 28S ribosomal protein S26, mitochondrial | P67936 | MRPS26 | 0,64910 | -1,38552 | 19,77208 | 21,86514 | 21,82625 | 22,35353 | 23,78776 | 21,47872 |
| | DnaJ homolog subfamily B member 1 | Q14254 | DNAJB1 | 0,64876 | 2,14626 | 20,52640 | 20,76846 | 20,60359 | 15,50000 | 19,92997 | 20,02972 |
| | Ribosome biogenesis protein WDR12 | P07197 | WDR12 | 0,64787 | -0,68066 | 21,57948 | 20,26877 | 20,90727 | 21,10303 | 22,09632 | 21,59814 |
| | Probable ATP-dependent RNA helicase DDX6 | O15371 | DDX6 | 0,64670 | -0,48982 | 22,65005 | 22,58183 | 23,45538 | 23,22688 | 23,15448 | 23,77535 |
| | Diacylglycerol kinase epsilon | Q9Y5Q8 | DGKE | 0,64644 | -0,59733 | 19,29923 | 19,88618 | 20,21691 | 20,07078 | 21,03765 | 20,08589 |
| | Rabankyrin-5 | Q4VCS5 | ANKFY1 | 0,64573 | 2,31176 | 22,06812 | 19,81606 | 20,30341 | 20,11832 | 15,50000 | 19,63398 |
| | Destrin | P21333 | DSTN | 0,64524 | 0,90981 | 22,61675 | 23,01635 | 21,71297 | 22,54774 | 21,12206 | 20,94685 |
| | Protein SEC13 homolog | Q9Y6G9 | SEC13 | 0,64401 | 0,98684 | 22,08076 | 22,02682 | 21,02985 | 21,88450 | 19,86505 | 20,42737 |
| | NADH dehydrogenase [ubiquinone] iron-sulfur protein 3, mitochondrial | O75369 | NDUFS3 | 0,64378 | 2,62120 | 21,41116 | 22,09458 | 21,81636 | 15,50000 | 21,10361 | 20,85491 |
| | Tyrosine-protein phosphatase non-receptor type 1 | O00159 | PTPN1 | 0,64345 | 2,00147 | 20,68927 | 20,31108 | 19,76341 | 15,50000 | 19,50159 | 19,75778 |
| | Cilia- and flagella-associated protein 20 | O95425 | CFAP20 | 0,64210 | -1,10467 | 20,89970 | 22,79769 | 22,92094 | 22,55431 | 23,37849 | 23,99955 |
| | Actin-related protein 2/3 complex subunit 3 | P54727 | ARPC3 | 0,64076 | -0,55316 | 21,18641 | 21,42829 | 21,90779 | 22,24432 | 21,42542 | 22,51222 |
| | Enoyl-CoA delta isomerase 2, mitochondrial | P57088 | ECI2 | 0,64062 | 2,37959 | 21,23497 | 21,46842 | 20,80144 | 15,50000 | 20,88472 | 19,98134 |
| | Regulator of nonsense transcripts 2 | P78406 | UPF2 | 0,64012 | -0,84451 | 23,86089 | 23,55532 | 23,79045 | 23,88995 | 25,75001 | 24,10021 |
| | Pescadillo homolog | P33993 | PES1 | 0,63767 | -2,87571 | 15,50000 | 21,25084 | 21,84946 | 22,44448 | 22,73822 | 22,04473 |
| | ATPase family AAA domain-containing protein 1 | P50402 | ATAD1 | 0,63764 | 2,11085 | 20,50300 | 20,76174 | 20,06789 | 15,50000 | 20,48778 | 19,01231 |
| | H/ACA ribonucleoprotein complex subunit 1 | Q9Y3F4 | GAR1 | 0,63666 | -2,95958 | 15,50000 | 21,32113 | 21,89796 | 22,47298 | 21,75287 | 23,37199 |
| | Mediator of RNA polymerase II transcription subunit 19 | Q9UQ88 | MED19 | 0,63606 | -2,01261 | 15,50000 | 19,88804 | 19,23963 | 20,19160 | 19,53269 | 20,94120 |
| | Poly(rC)-binding protein 1 | P49411 | PCBP1 | 0,63551 | 0,69104 | 23,67142 | 24,70946 | 23,76769 | 24,04968 | 22,82912 | 23,19666 |
| | Superkiller viralicidic activity 2-like 2 | P24666 | SKIV2L2 | 0,63531 | -0,59495 | 22,82001 | 24,18571 | 23,89319 | 24,30222 | 24,30716 | 24,07439 |
| | RuvB-like 2 | P35637 | RUVBL2 | 0,63437 | -0,22593 | 25,00920 | 24,69200 | 24,51424 | 24,83928 | 24,97496 | 25,07900 |
| | Signal recognition particle subunit SRP72 | Q15007 | SRP72 | 0,63230 | -1,80599 | 21,32317 | 24,08950 | 23,73769 | 23,59889 | 26,71581 | 24,25364 |
| | 60S ribosomal protein L27 | Q9Y5K5 | RPL27 | 0,63129 | -4,12998 | 15,50000 | 22,00761 | 25,39940 | 24,16077 | 25,86481 | 25,27138 |
| | Interferon-induced protein with tetratricopeptide repeats 5 | P20073 | IFIT5 | 0,63062 | -1,13953 | 19,41749 | 20,03537 | 20,06618 | 20,24400 | 22,55018 | 20,14345 |
| | Basic leucine zipper and W2 domain-containing protein 2 | Q96EP5 | BZW2 | 0,63022 | 1,75190 | 19,67903 | 19,17823 | 19,84512 | 15,50000 | 19,72216 | 18,22454 |
| | Serine/threonine-protein phosphatase 2A catalytic subunit alpha isoform;Serine/threonine-protein phosphatase 2A catalytic subunit beta isoform | P19474 | PPP2CA;PPP2CB | 0,62895 | 1,86871 | 20,14942 | 20,01045 | 19,96268 | 15,50000 | 19,43742 | 19,57900 |
| | RNA polymerase II elongation factor ELL | P11021 | ELL | 0,62779 | -2,31856 | 15,50000 | 20,76361 | 19,92686 | 21,42562 | 20,42931 | 21,29120 |
| | Serine/threonine-protein kinase 26 | Q15436 | STK26 | 0,62762 | 0,94624 | 21,76118 | 22,15542 | 21,19376 | 20,71062 | 21,85129 | 19,70974 |
| | 40S ribosomal protein S3a | Q9Y490 | RPS3A | 0,62736 | -0,83509 | 26,80845 | 26,03149 | 27,07572 | 27,05662 | 28,48987 | 26,87444 |
| | 60S ribosomal protein L27a | Q9NYL9 | RPL27A | 0,62715 | -1,28857 | 21,61239 | 22,42470 | 24,57005 | 23,63943 | 24,58869 | 24,24474 |

| Significant | Protein names | UniProtID | Gene names | -LOG(P-value) | Difference | CXCR4_1 | CXCR4_2 | CXCR4_3 | MOCK_1 | MOCK_2 | MOCK_3 |
|---|---|---|---|---|---|---|---|---|---|---|---|
| | Nucleosome assembly protein 1-like 4 | P01834 | NAP1L4 | 0,62692 | 0,78971 | 19,79839 | 20,20670 | 21,53901 | 20,12805 | 19,40520 | 19,64171 |
| | Phosphoribosyl pyrophosphate synthase-associated protein 1 | P22234 | PRPSAP1 | 0,62343 | 0,72594 | 21,47719 | 22,06178 | 20,80609 | 21,45332 | 20,52267 | 20,19123 |
| | 14-3-3 protein gamma;14-3-3 protein gamma, N-terminally processed | Q9Y262 | YWHAG | 0,62317 | 0,90616 | 21,40904 | 20,53991 | 20,07835 | 20,74156 | 18,93988 | 19,62737 |
| | Serine-threonine kinase receptor-associated protein | P60228 | STRAP | 0,62138 | 0,39631 | 22,55075 | 22,61890 | 22,45171 | 22,65379 | 21,67762 | 22,10101 |
| | Signal recognition particle receptor subunit beta | Q16643 | SRPRB | 0,61911 | 0,71299 | 23,27739 | 23,78417 | 22,55328 | 21,96325 | 23,21589 | 22,29673 |
| | Transcription initiation factor TFIID subunit 4 | Q96J01 | TAF4 | 0,61878 | -2,18344 | 15,50000 | 20,35068 | 19,99879 | 21,15159 | 20,99148 | 20,25673 |
| | Flotillin-1 | P08670 | FLOT1 | 0,61873 | 0,70686 | 23,15097 | 22,31932 | 22,05996 | 21,91900 | 21,07025 | 22,42042 |
| | 60S ribosomal protein L9 | P50454 | RPL9 | 0,61826 | -1,56698 | 23,28546 | 23,75297 | 26,33125 | 24,78915 | 26,88040 | 26,40106 |
| | Puromycin-sensitive aminopeptidase | P30876 | NPEPPS | 0,61811 | 2,90114 | 18,76439 | 22,88891 | 23,02036 | 15,50000 | 20,34093 | 20,12931 |
| | Translation initiation factor eIF-2B subunit delta | P51148 | EIF2B4 | 0,61762 | -0,83381 | 21,74669 | 22,99664 | 22,82328 | 23,08560 | 24,25947 | 22,72297 |
| | Protein numb homolog | P0DMV9 | NUMB | 0,61491 | -2,52544 | 15,50000 | 21,13182 | 20,87770 | 21,87358 | 21,85498 | 21,35730 |
| | Signal recognition particle 14 kDa protein | Q9BU76 | SRP14 | 0,61479 | -1,10449 | 20,69532 | 22,42573 | 23,25465 | 22,67420 | 23,38350 | 23,63145 |
| | 60S ribosomal protein L30 | Q92900 | RPL30 | 0,61235 | -1,56778 | 22,62192 | 22,20275 | 25,16603 | 23,54262 | 25,46880 | 25,68263 |
| | Coiled-coil domain-containing protein 47 | Q13247 | CCDC47 | 0,61201 | -0,93020 | 23,28912 | 24,63467 | 24,34015 | 24,45029 | 26,11048 | 24,49375 |
| | Fragile X mental retardation syndrome-related protein 1 | Q15233 | FXR1 | 0,61040 | -1,05135 | 21,30986 | 23,12155 | 23,79996 | 23,37385 | 24,03465 | 23,97692 |
| | Protein pelota homolog | Q52LJ0 | PELO | 0,61015 | 1,59765 | 19,52742 | 19,82930 | 18,66557 | 15,50000 | 18,88950 | 18,83984 |
| | H/ACA ribonucleoprotein complex non-core subunit NAF1 | Q53GQ0 | NAF1 | 0,60827 | -2,60715 | 15,50000 | 20,79011 | 21,26917 | 22,11009 | 20,76531 | 22,50531 |
| | Vacuolar protein sorting-associated protein 35 | Q16543 | VPS35 | 0,60826 | 3,00499 | 21,64046 | 23,28884 | 23,35877 | 15,50000 | 22,31141 | 21,46169 |
| | CCAAT/enhancer-binding protein zeta | O95429 | CEBPZ | 0,60742 | -1,06176 | 22,08080 | 22,97348 | 22,90334 | 23,10561 | 25,16676 | 22,87053 |
| | Ribonuclease P protein subunit p38 | O43324 | RPP38 | 0,60310 | -2,00999 | 19,65372 | 15,50000 | 20,23503 | 20,34906 | 20,60982 | 20,45984 |
| | Replication factor C subunit 4 | Q92945 | RFC4 | 0,60226 | -0,45858 | 22,61666 | 23,38271 | 23,27113 | 23,10290 | 23,60286 | 23,94047 |
| | Emerin | P51648 | EMD | 0,60077 | 0,39752 | 22,08777 | 22,08923 | 21,32789 | 21,71179 | 21,18198 | 21,41854 |
| | Cell division cycle protein 123 homolog | Q6P1M0 | CDC123 | 0,59882 | 2,28979 | 21,24528 | 21,19418 | 21,13615 | 15,50000 | 20,92180 | 20,28444 |
| | Partitioning defective 3 homolog | Q13813 | PARD3 | 0,59576 | -1,30140 | 18,90839 | 20,11540 | 20,37687 | 20,18713 | 22,83260 | 20,28512 |
| | Prolactin regulatory element-binding protein | Q5T9A4 | PREB | 0,59399 | 2,34731 | 21,52535 | 21,45643 | 21,13464 | 15,50000 | 20,74755 | 20,82695 |
| | ADP-ribosylation factor 4 | Q9NZB2 | ARF4 | 0,59221 | 0,87592 | 23,02540 | 23,50118 | 22,12266 | 22,68915 | 22,35289 | 20,97946 |
| | Nucleolar protein 56 | A0A1W2PNV4 | NOP56 | 0,59108 | -1,55522 | 21,84827 | 25,22159 | 24,78850 | 26,12076 | 24,49777 | 25,90550 |
| | RNA-binding protein with serine-rich domain 1 | P45880 | RNPS1 | 0,58801 | -2,76617 | 15,50000 | 20,66249 | 21,97980 | 21,25425 | 21,64051 | 23,54604 |
| | Transcriptional repressor p66-beta | O60762 | GATAD2B | 0,58755 | -2,17251 | 15,50000 | 20,29041 | 19,88580 | 20,43002 | 21,90591 | 19,85781 |
| | A-kinase anchor protein 17A | P25398 | AKAP17A | 0,58740 | -2,86439 | 15,50000 | 21,21437 | 21,89008 | 20,82648 | 23,37981 | 22,99132 |
| | Apoptosis inhibitor 5 | P53621 | API5 | 0,58404 | 2,42779 | 21,41581 | 21,03879 | 22,22359 | 15,50000 | 20,70708 | 21,18773 |
| | 40S ribosomal protein SA | Q9P0K7 | RPSA | 0,58261 | -0,70952 | 23,79174 | 23,39611 | 25,02064 | 24,31624 | 24,92936 | 25,09144 |
| | SRSF protein kinase 2;SRSF protein kinase 2 N-terminal;SRSF protein kinase 2 C-terminal | Q9BWF3 | SRPK2 | 0,58229 | -2,25384 | 15,50000 | 20,08057 | 21,01652 | 21,56267 | 20,62435 | 21,17158 |
| | Nuclease-sensitive element-binding protein 1 | O00566 | YBX1 | 0,58131 | -0,86017 | 24,00204 | 24,48075 | 25,71853 | 24,97946 | 25,41361 | 26,38877 |
| | WD repeat-containing protein 5 | O75431 | WDR5 | 0,58051 | -0,43702 | 22,36864 | 22,74259 | 22,29922 | 23,17023 | 22,29707 | 23,25422 |
| | 28S ribosomal protein S34, mitochondrial | O14979 | MRPS34 | 0,58025 | -2,79024 | 15,50000 | 20,78373 | 22,02249 | 21,32125 | 23,75868 | 21,59701 |
| | TBC1 domain family member 10B | Q9P2J5 | TBC1D10B | 0,57948 | -1,84684 | 19,20460 | 22,38184 | 23,72661 | 22,73329 | 23,79064 | 24,32965 |
| | 28S ribosomal protein S11, mitochondrial | Q6IAN0 | MRPS11 | 0,57916 | -2,49287 | 15,50000 | 20,24284 | 20,97841 | 20,80884 | 23,08571 | 20,30530 |
| | Cyclin-dependent kinase 4 | P47756 | CDK4 | 0,57696 | 1,80623 | 20,19868 | 20,29480 | 19,60152 | 15,50000 | 19,24158 | 19,93474 |
| | 28S ribosomal protein S22, mitochondrial | P42166 | MRPS22 | 0,57630 | -0,86468 | 22,80900 | 22,93200 | 23,21028 | 23,42842 | 25,13766 | 22,97925 |
| | Pentatricopeptide repeat domain-containing protein 3, mitochondrial | P11142 | PTCD3 | 0,57475 | -1,30336 | 21,44859 | 21,50300 | 23,53493 | 22,41274 | 24,89291 | 23,09096 |
| | ATP-dependent RNA helicase DDX55 | O95347 | DDX55 | 0,57366 | -3,02181 | 15,50000 | 22,52578 | 22,51644 | 23,13874 | 23,00501 | 23,46389 |
| | Transgelin-2 | Q9NVI7 | TAGLN2 | 0,57191 | -1,93140 | 15,50000 | 15,50000 | 19,12502 | 20,42358 | 17,67975 | 17,81591 |
| | Angio-associated migratory cell protein | Q15907 | AAMP | 0,57127 | 2,06595 | 21,45914 | 20,40260 | 19,99369 | 15,50000 | 20,52860 | 19,62899 |
| | CXXC-type zinc finger protein 1 | Q9NU22 | CXXC1 | 0,57095 | -2,76149 | 15,50000 | 22,01117 | 21,68265 | 21,75752 | 22,70961 | 23,01115 |
| | General transcription factor 3C polypeptide 1 | P52272 | GTF3C1 | 0,57045 | 2,13453 | 21,32108 | 20,40613 | 20,82873 | 15,50000 | 20,90375 | 19,74862 |
| | WD repeat-containing protein 82 | Q01082 | WDR82 | 0,57002 | -0,46323 | 21,60386 | 21,26728 | 21,98235 | 21,71762 | 21,85692 | 22,66865 |
| | Lysine--tRNA ligase | Q9BXP5 | KARS | 0,56898 | -3,32611 | 15,50000 | 22,73746 | 22,22058 | 23,01158 | 25,66058 | 21,76422 |
| | Sorting nexin-6;Sorting nexin-6, N-terminally processed | O00186 | SNX6 | 0,56773 | 1,79998 | 19,52772 | 20,32389 | 20,25442 | 15,50000 | 19,99092 | 19,21516 |
| | RNA-binding protein 39 | Q6WCQ1 | RBM39 | 0,56768 | -1,42608 | 23,68089 | 26,39203 | 26,93146 | 26,66343 | 26,55741 | 28,06178 |
| | Mediator of RNA polymerase II transcription subunit 4 | P68104 | MED4 | 0,56724 | -0,64357 | 21,43532 | 22,50368 | 21,91379 | 22,96190 | 21,79844 | 23,02315 |
| | Small nuclear ribonucleoprotein-associated protein N;Small nuclear ribonucleoprotein-associated proteins B and B | O43707 | SNRPN;SNRPB | 0,56660 | -2,75413 | 15,50000 | 20,94027 | 22,57994 | 21,88997 | 22,49140 | 22,90123 |
| | 28S ribosomal protein S15, mitochondrial | Q9Y512 | MRPS15 | 0,56565 | -2,54094 | 15,50000 | 21,30480 | 21,28743 | 21,75927 | 22,83327 | 21,12250 |
| | G protein-coupled receptor kinase 6 | P38646 | GRK6 | 0,56542 | -1,77529 | 15,50000 | 19,31935 | 19,94592 | 19,78708 | 20,16349 | 20,14058 |
| | Zinc finger CCCH domain-containing protein 4 | Q96HS1 | ZC3H4 | 0,56330 | -1,32032 | 19,03314 | 20,94399 | 22,18077 | 21,50698 | 21,58394 | 23,02795 |
| | SWI/SNF-related matrix-associated actin-dependent regulator of chromatin subfamily E member 1 | Q13595 | SMARCE1 | 0,56325 | -0,90183 | 22,23532 | 21,85802 | 20,85392 | 22,51711 | 23,57045 | 21,56519 |
| | Bystin | P25205 | BYSL | 0,56279 | -0,76306 | 19,80477 | 21,37565 | 21,40665 | 21,12679 | 22,12225 | 21,62721 |
| | Chromodomain-helicase-DNA-binding protein 7 | Q9H2U1 | CHD7 | 0,56063 | -2,50951 | 20,54868 | 15,50000 | 21,79990 | 20,84514 | 22,45854 | 22,07343 |
| | N-alpha-acetyltransferase 15, NatA auxiliary subunit | P53985 | NAA15 | 0,55737 | -2,92034 | 15,50000 | 20,87418 | 21,59596 | 20,39874 | 24,76056 | 21,57187 |
| | 60S ribosomal protein L11 | P63151 | RPL11 | 0,55635 | -0,38515 | 24,33899 | 24,46214 | 25,04018 | 24,64147 | 25,39326 | 24,96204 |
| | Cell cycle and apoptosis regulator protein 2 | Q9GZR7 | CCAR2 | 0,55531 | -0,41080 | 20,96754 | 20,29446 | 21,28923 | 21,45694 | 20,97365 | 21,35305 |

| Significant | Protein names | UniProtID | Gene names | -LOG(P-value) | Difference | CXCR4_1 | CXCR4_2 | CXCR4_3 | MOCK_1 | MOCK_2 | MOCK_3 |
|---|---|---|---|---|---|---|---|---|---|---|---|
| | ATP synthase subunit gamma, mitochondrial | O15144 | ATP5C1 | 0,55421 | -1,29340 | 24,49179 | 23,69381 | 25,00273 | 24,26699 | 27,52066 | 25,28090 |
| | Chromatin assembly factor 1 subunit A | Q9BS26 | CHAF1A | 0,55333 | -0,76912 | 21,82268 | 21,84713 | 20,84759 | 22,43606 | 23,08463 | 21,30408 |
| | Brain-specific angiogenesis inhibitor 1-associated protein 2-like protein 1 | Q8WWI1 | BAIAP2L1 | 0,55175 | -2,03236 | 15,50000 | 20,78165 | 19,66570 | 20,36922 | 21,23421 | 20,44101 |
| | HBS1-like protein | P20719 | HBS1L | 0,55063 | -3,49508 | 15,50000 | 23,97552 | 23,79600 | 24,12731 | 25,09067 | 24,53878 |
| | Nucleolar GTP-binding protein 1 | Q13045 | GTPBP4 | 0,54979 | -3,33128 | 15,50000 | 23,15553 | 23,64889 | 23,22808 | 24,88985 | 24,18033 |
| | Pyrroline-5-carboxylate reductase 3 | Q9GZS3 | PYCRL | 0,54891 | 1,80305 | 20,57552 | 20,41452 | 19,29370 | 15,50000 | 19,61723 | 19,75734 |
| | Mediator of RNA polymerase II transcription subunit 20 | P62269 | MED20 | 0,54688 | -0,81222 | 21,77398 | 23,18739 | 21,75499 | 23,92234 | 22,39221 | 22,83848 |
| | Voltage-dependent anion-selective channel protein 3 | O14757 | VDAC3 | 0,54657 | 0,75714 | 24,85605 | 24,68073 | 23,61054 | 22,95697 | 24,53925 | 23,37968 |
| | 14-3-3 protein beta/alpha;14-3-3 protein beta/alpha, N-terminally processed | P51610 | YWHAB | 0,54643 | 1,12146 | 22,30265 | 22,39979 | 21,64302 | 22,68637 | 19,75708 | 20,53763 |
| | WD repeat-containing protein 3 | P12268 | WDR3 | 0,54631 | -2,93532 | 15,50000 | 20,94928 | 22,35391 | 20,45322 | 24,33381 | 22,82211 |
| | rRNA 2-O-methyltransferase fibrillarin | O94832 | FBL | 0,54610 | -3,17979 | 15,50000 | 23,06296 | 23,25811 | 24,05333 | 23,14342 | 24,16369 |
| | Probable ATP-dependent RNA helicase DDX17 | Q9BZF9 | DDX17 | 0,54336 | -0,66032 | 24,55145 | 25,54478 | 26,17775 | 25,59665 | 26,21035 | 26,44795 |
| | Mitochondrial import receptor subunit TOM34 | Q92734 | TOMM34 | 0,54228 | 2,12752 | 21,15697 | 19,96860 | 21,54326 | 15,50000 | 19,79659 | 20,98968 |
| | Chromodomain-helicase-DNA-binding protein 4 | P52907 | CHD4 | 0,54134 | -1,22346 | 21,01318 | 21,41854 | 22,12165 | 21,66210 | 24,62271 | 21,93894 |
| | 28S ribosomal protein S31, mitochondrial | P35580 | MRPS31 | 0,54081 | -1,05610 | 21,10047 | 21,56491 | 22,44655 | 22,65676 | 24,13657 | 21,48689 |
| | 40S ribosomal protein S7 | O14974 | RPS7 | 0,53974 | -1,14654 | 22,93749 | 22,80520 | 24,76162 | 23,27014 | 25,47423 | 25,19955 |
| | Elongation factor 2 | Q6P1J9 | EEF2 | 0,53863 | 0,49770 | 25,12762 | 25,65581 | 26,35510 | 25,05234 | 24,98046 | 25,61263 |
| | Centrosomal protein of 55 kDa | P24534 | CEP55 | 0,53812 | -0,18072 | 19,41809 | 19,07467 | 19,28238 | 19,60525 | 19,23248 | 19,47957 |
| | Zinc finger CCCH domain-containing protein 14 | Q12904 | ZC3H14 | 0,53772 | -2,81390 | 15,50000 | 19,96056 | 22,27884 | 19,76600 | 23,62611 | 22,78899 |
| | 60S ribosomal protein L24 | Q08211 | RPL24 | 0,53654 | -3,45153 | 15,50000 | 21,83121 | 24,88465 | 22,90331 | 24,84470 | 24,82245 |
| | Protein disulfide-isomerase | Q9P035 | P4HB | 0,53602 | 0,79102 | 20,63759 | 19,75812 | 19,48148 | 20,09545 | 19,21556 | 18,19311 |
| | Alpha-taxilin | Q01085 | TXLNA | 0,53599 | -2,19474 | 15,50000 | 21,37087 | 19,66626 | 21,01393 | 21,86737 | 20,24005 |
| | Mediator of RNA polymerase II transcription subunit 28 | P51572 | MED28 | 0,53484 | -1,91565 | 15,50000 | 20,48188 | 19,69401 | 20,90331 | 19,84501 | 20,67452 |
| | DnaJ homolog subfamily B member 6 | Q9UBS4 | DNAJB6 | 0,53429 | -0,40090 | 20,90154 | 21,29013 | 20,23468 | 21,35305 | 21,30502 | 20,97099 |
| | Low molecular weight phosphotyrosine protein phosphatase | Q96SB3 | ACP1 | 0,53377 | 0,38233 | 22,32295 | 23,11288 | 22,43321 | 22,52636 | 22,33202 | 21,86367 |
| | Protein-L-isoaspartate(D-aspartate) O-methyltransferase | P61221 | PCMT1 | 0,53275 | -0,82932 | 22,05451 | 23,83277 | 23,84000 | 24,59138 | 23,42355 | 24,20030 |
| | Fragile X mental retardation protein 1 | Q12965 | FMR1 | 0,53229 | -1,10192 | 20,99321 | 21,54609 | 23,65829 | 22,39553 | 23,28842 | 23,81939 |
| | Zinc finger protein ubi-d4 | Q08380 | DPF2 | 0,53229 | -0,54823 | 21,12067 | 20,56579 | 20,07966 | 21,05563 | 21,76203 | 20,59314 |
| | U5 small nuclear ribonucleoprotein 200 kDa helicase | Q13838 | SNRNP200 | 0,53199 | -0,66510 | 24,71586 | 24,41300 | 25,41893 | 24,60682 | 26,12995 | 25,80632 |
| | Deleted in autism protein 1 | P62263 | C3orf58 | 0,53174 | 1,91082 | 20,46901 | 20,40499 | 20,68765 | 15,50000 | 19,67028 | 20,65892 |
| | Myosin light chain 6B | P19105 | MYL6B | 0,53063 | -0,41856 | 19,00454 | 19,24272 | 19,17816 | 19,88535 | 18,88047 | 19,91528 |
| | Pleiotropic regulator 1 | O14579 | PLRG1 | 0,53057 | -1,72246 | 19,50953 | 21,26659 | 22,85371 | 21,03006 | 24,67147 | 23,09568 |
| | Zinc finger CCHC domain-containing protein 3 | Q6PKG0 | ZCCHC3 | 0,52993 | -2,26082 | 15,50000 | 20,08200 | 21,59751 | 20,49726 | 21,69761 | 21,76709 |
| | Rab11 family-interacting protein 2 | O43795 | RAB11FIP2 | 0,52952 | -2,24688 | 15,50000 | 21,36645 | 20,79003 | 21,59259 | 21,33364 | 21,47090 |
| | Scaffold attachment factor B1 | P42677 | SAFB | 0,52701 | -2,58646 | 15,50000 | 21,03926 | 22,59170 | 22,42270 | 21,94792 | 22,51970 |
| | Heterogeneous nuclear ribonucleoprotein F;Heterogeneous nuclear ribonucleoprotein F, N-terminally processed | Q12830 | HNRNPF | 0,52633 | -0,70786 | 24,56634 | 24,17881 | 24,96512 | 24,42265 | 25,11648 | 26,29470 |
| | RNA-binding protein 14 | Q92804 | RBM14 | 0,52456 | -0,94769 | 22,35950 | 24,86623 | 22,97462 | 24,56448 | 24,62879 | 23,85015 |
| | Transcription activator BRG1 | Q01844 | SMARCA4 | 0,52435 | -0,34268 | 23,43097 | 23,54109 | 22,88764 | 24,03322 | 23,48173 | 23,37279 |
| | Nucleolar GTP-binding protein 2 | P26640 | GNL2 | 0,52425 | -2,42605 | 15,50000 | 19,88712 | 21,69655 | 19,81814 | 22,78357 | 21,76012 |
| | PAX3- and PAX7-binding protein 1 | Q9Y6B6 | PAXBP1 | 0,52395 | -2,94106 | 15,50000 | 22,51133 | 23,13751 | 22,88945 | 23,15235 | 23,93022 |
| | 40S ribosomal protein S11 | Q99615 | RPS11 | 0,52388 | -0,71659 | 22,28269 | 22,97002 | 23,87540 | 23,00878 | 24,29095 | 23,97815 |
| | Luc7-like protein 3 | P27694 | LUC7L3 | 0,52292 | -0,81909 | 21,08297 | 21,26957 | 22,89449 | 21,85198 | 22,71465 | 23,13768 |
| | Pre-mRNA 3-end-processing factor FIP1 | P81605 | FIP1L1 | 0,52180 | -2,38233 | 15,50000 | 21,24458 | 21,63971 | 21,45101 | 22,42864 | 21,65163 |
| | Proteasome-associated protein ECM29 homolog | Q00325 | ECM29 | 0,52125 | 2,36676 | 20,13934 | 19,01892 | 20,30318 | 15,50000 | 21,36116 | 15,50000 |
| | Protein deglycase DJ-1 | P23526 | PARK7 | 0,52018 | 0,90238 | 21,13646 | 23,03803 | 21,60896 | 21,08076 | 20,12641 | 21,86914 |
| | 40S ribosomal protein S9 | P07814 | RPS9 | 0,52016 | -1,86183 | 21,13057 | 23,96049 | 26,18314 | 24,46656 | 26,12591 | 26,26721 |
| | 7-dehydrocholesterol reductase | P26368 | DHCR7 | 0,51948 | 0,70492 | 22,27239 | 22,57427 | 21,52970 | 21,76247 | 22,07946 | 20,41967 |
| | 26S proteasome non-ATPase regulatory subunit 4 | P31942 | PSMD4 | 0,51850 | 0,73951 | 23,27555 | 23,89763 | 22,38759 | 22,52203 | 23,18226 | 21,63795 |
| | BMP-2-inducible protein kinase | Q10570 | BMP2K | 0,51767 | -2,25931 | 15,50000 | 21,15178 | 21,30341 | 21,85548 | 21,54680 | 21,33085 |
| | Nucleolar pre-ribosomal-associated protein 1 | P41252 | URB1 | 0,51763 | -0,91086 | 21,45583 | 20,96402 | 20,94514 | 21,30145 | 23,54050 | 21,25563 |
| | Tyrosine-protein kinase JAK1 | P63244 | JAK1 | 0,51750 | -1,81593 | 15,50000 | 19,77666 | 18,87529 | 18,98031 | 21,51191 | 19,10753 |
| | Splicing factor 3A subunit 3 | Q96K37 | SF3A3 | 0,51611 | -0,82074 | 20,41782 | 21,45719 | 22,47234 | 21,87144 | 21,93409 | 23,00403 |
| | Protein AAR2 homolog | Q14444 | AAR2 | 0,51561 | 1,57611 | 19,90730 | 19,85767 | 19,38886 | 15,50000 | 19,74361 | 19,18188 |
| | Serine/threonine-protein kinase Nek7 | Q13547 | NEK7 | 0,51552 | 1,44183 | 19,02210 | 19,95631 | 18,94086 | 15,50000 | 19,11502 | 18,97874 |
| | 40S ribosomal protein S20 | Q13155 | RPS20 | 0,51535 | -1,39237 | 22,49351 | 21,64764 | 25,39842 | 23,91333 | 24,76663 | 25,03670 |
| | Protein arginine N-methyltransferase 5;Protein arginine N-methyltransferase 5, N-terminally processed | O94813 | PRMT5 | 0,51511 | -1,24244 | 23,36132 | 26,24274 | 26,26076 | 27,02538 | 25,65905 | 26,90770 |
| | SWI/SNF complex subunit SMARCC1 | O15047 | SMARCC1 | 0,51504 | -0,26477 | 23,91141 | 23,76486 | 23,36413 | 24,23425 | 23,89763 | 23,70282 |
| | 60S ribosomal protein L15 | Q86YS7 | RPL15 | 0,51268 | -3,35258 | 15,50000 | 22,85041 | 24,86207 | 23,74222 | 24,63267 | 24,89532 |
| | Serine hydroxymethyltransferase, mitochondrial | Q92499 | SHMT2 | 0,51227 | 2,24086 | 21,50693 | 21,11889 | 21,89438 | 15,50000 | 20,65648 | 21,64112 |
| | Stromal cell-derived factor 2 | P06396 | SDF2 | 0,51227 | -1,45479 | 15,50000 | 19,45083 | 18,92758 | 19,65433 | 19,29116 | 19,29728 |
| | tRNA-splicing ligase RtcB homolog | Q12792 | RTCB | 0,51195 | -0,32016 | 25,08493 | 24,34861 | 24,58589 | 25,02855 | 25,26410 | 24,68726 |
| | Myb-binding protein 1A | O00571 | MYBBP1A | 0,51029 | -1,00310 | 25,77407 | 24,87108 | 26,06491 | 25,41480 | 28,06389 | 26,24067 |

| Significant | Protein names | UniProtID | Gene names | -LOG(P-value) | Difference | CXCR4_1 | CXCR4_2 | CXCR4_3 | MOCK_1 | MOCK_2 | MOCK_3 |
|---|---|---|---|---|---|---|---|---|---|---|---|
| | Retinoblastoma-binding protein 5 | O60884 | RBBP5 | 0,50944 | -0,75920 | 21,09397 | 22,51304 | 22,21963 | 22,63354 | 21,89067 | 23,58003 |
| | RNA-binding motif protein, X-linked 2 | P36873 | RBMX2 | 0,50914 | -2,03429 | 15,50000 | 20,79836 | 19,88219 | 21,47586 | 19,51867 | 21,28889 |
| | 28S ribosomal protein S9, mitochondrial | P26641 | MRPS9 | 0,50719 | -0,72686 | 22,19520 | 22,65491 | 22,35523 | 22,74999 | 24,32718 | 22,30875 |
| | Protein RCC2 | P46782 | RCC2 | 0,50631 | -0,28999 | 22,85064 | 22,91245 | 23,32828 | 23,56407 | 22,92209 | 23,47519 |
| | Round spermatid basic protein 1 | P62140 | RSBN1 | 0,50549 | -2,66713 | 15,50000 | 22,80361 | 21,77937 | 23,09063 | 22,01499 | 22,97874 |
| | N-acetyltransferase 10 | P50995 | NAT10 | 0,50487 | -0,79193 | 23,98971 | 25,35454 | 25,82231 | 25,78009 | 25,17234 | 26,58992 |
| | Ataxin-2-like protein | P62701 | ATXN2L | 0,50261 | 0,49273 | 22,61419 | 21,70548 | 21,98874 | 22,23792 | 21,09822 | 21,49409 |
| | Plectin | P31689 | PLEC | 0,50259 | 0,64240 | 24,49234 | 23,85034 | 23,53445 | 24,07447 | 22,42034 | 23,45513 |
| | Suppressor of G2 allele of SKP1 homolog | Q9HCE1 | SUGT1 | 0,50250 | 1,65913 | 20,36239 | 19,99314 | 19,28071 | 20,21229 | 15,50000 | 18,94655 |
| | Lamin-B receptor | P29692 | LBR | 0,50204 | -0,87142 | 21,23707 | 21,18174 | 21,45909 | 21,45136 | 23,67120 | 21,36959 |
| | ESF1 homolog | Q9UQE7 | ESF1 | 0,49996 | -0,57307 | 22,28238 | 23,48148 | 23,24932 | 24,03783 | 22,91280 | 23,78177 |
| | Serine/threonine-protein phosphatase 2B catalytic subunit beta isoform;Serine/threonine-protein phosphatase 2B catalytic subunit gamma isoform | Q14683 | PPP3CB;PPP3CC | 0,49964 | 1,33129 | 19,63828 | 18,93413 | 18,68490 | 15,50000 | 18,79592 | 18,96753 |
| | Transcription elongation factor B polypeptide 3 | Q15459 | TCEB3 | 0,49886 | -2,60387 | 15,50000 | 21,89390 | 22,01744 | 21,04900 | 23,64196 | 22,53198 |
| | Protein LYRIC | Q00839 | MTDH | 0,49804 | -3,38627 | 15,50000 | 23,78616 | 24,71597 | 23,76051 | 25,13806 | 25,26238 |
| | Ubinuclein-1 | P20700 | UBN1 | 0,49775 | -2,05492 | 15,50000 | 20,30997 | 20,64342 | 22,21146 | 20,54670 | 19,85998 |
| | 28S ribosomal protein S29, mitochondrial | P62316 | DAP3 | 0,49607 | -0,86334 | 22,58976 | 22,78429 | 22,91845 | 23,49363 | 24,99425 | 22,39464 |
| | Copine-1 | P63261 | CPNE1 | 0,49532 | 0,51436 | 20,73760 | 21,06131 | 21,18489 | 20,37729 | 19,78708 | 21,27634 |
| | Cancer/testis antigen family 45 member A10;Cancer/testis antigen family 45 member A3;Cancer/testis antigen family 45 member A1;Cancer/testis antigen family 45 member A2;Cancer/testis antigen family 45 member A9;Cancer/testis antigen family 45 member A8;Cancer/testis antigen family 45 member A7;Cancer/testis antigen family 45 member A5;Cancer/testis antigen family 45 member A6 | Q06830 | CT45A10;CT45A3;CT45A1;CT45A2;CT45A9;CT45A8;CT45A7;CT45A5;CT45A6 | 0,49519 | -3,08976 | 15,50000 | 24,13594 | 22,92991 | 24,53255 | 23,33838 | 23,96420 |
| | Translation factor GUF1, mitochondrial | Q14566 | GUF1 | 0,49485 | -2,11247 | 15,50000 | 19,38104 | 21,04780 | 19,37585 | 22,37936 | 20,51104 |
| | 116 kDa U5 small nuclear ribonucleoprotein component | P29375 | EFTUD2 | 0,49484 | -0,50591 | 24,17988 | 23,88651 | 24,64960 | 23,97281 | 25,08780 | 25,17310 |
| | Single-stranded DNA-binding protein, mitochondrial | Q96GM8 | SSBP1 | 0,49479 | -1,93908 | 20,73455 | 15,50000 | 20,45914 | 21,17767 | 20,71381 | 20,61944 |
| | RalA-binding protein 1 | Q53EZ4 | RALBP1 | 0,49354 | -1,99200 | 15,50000 | 20,81909 | 20,45302 | 20,23760 | 21,60340 | 20,90713 |
| | Chromatin complexes subunit BAP18 | P40937 | BAP18;RNASEK-C17orf49 | 0,49346 | -2,68776 | 15,50000 | 22,75959 | 22,09703 | 23,61796 | 21,83596 | 22,96600 |
| | Bromodomain adjacent to zinc finger domain protein 1A | Q99873 | BAZ1A | 0,49274 | -2,32534 | 21,00333 | 15,50000 | 19,02451 | 19,82333 | 23,38007 | 19,30046 |
| | Splicing factor 3B subunit 3 | Q9ULV4 | SF3B3 | 0,49248 | -0,66970 | 25,91668 | 26,41099 | 27,47777 | 26,66980 | 27,19009 | 27,95466 |
| | Cleavage and polyadenylation specificity factor subunit 6 | Q9Y6C9 | CPSF6 | 0,49193 | -1,68111 | 15,50000 | 19,05298 | 20,50048 | 19,82552 | 20,07835 | 20,19292 |
| | AP-3 complex subunit sigma-2 | P62277 | AP3S2;C15orf38-AP3S2 | 0,49129 | -2,22034 | 15,50000 | 21,72130 | 20,46533 | 22,32658 | 20,52420 | 21,49687 |
| | Protein LSM14 homolog A | Q96IZ0 | LSM14A | 0,49051 | -1,60474 | 15,50000 | 19,56947 | 19,92434 | 20,11159 | 19,67931 | 20,01713 |
| | Putative ribosome-binding factor A, mitochondrial | P05141 | RBFA | 0,49030 | -2,36140 | 15,50000 | 21,32438 | 21,63227 | 21,93467 | 22,92778 | 20,67840 |
| | Estradiol 17-beta-dehydrogenase 11 | Q15654 | HSD17B11 | 0,48705 | -1,85177 | 20,86559 | 15,50000 | 19,78633 | 20,73983 | 20,85696 | 20,11044 |
| | U3 small nucleolar RNA-associated protein 6 homolog | Q13162 | UTP6 | 0,48584 | -1,69216 | 15,50000 | 20,07025 | 19,55733 | 20,18374 | 20,79186 | 19,22844 |
| | Malectin | P53350 | MLEC | 0,48493 | 1,56185 | 19,91191 | 19,74084 | 19,92851 | 15,50000 | 19,75867 | 19,63704 |
| | ATP-binding cassette sub-family D member 3 | P23246 | ABCD3 | 0,48403 | -0,59308 | 22,75452 | 24,15329 | 23,71398 | 23,84938 | 24,80449 | 23,74714 |
| | RAF proto-oncogene serine/threonine-protein kinase | Q8NCA5 | RAF1 | 0,48233 | -1,93080 | 15,50000 | 20,59459 | 20,68046 | 20,73455 | 21,43761 | 20,39530 |
| | A-kinase anchor protein 8-like | Q13242 | AKAP8L | 0,48207 | -1,87298 | 15,50000 | 20,81644 | 20,22423 | 20,92673 | 20,80041 | 20,43247 |
| | Fatty acyl-CoA reductase 1 | P62995 | FAR1 | 0,48206 | 0,67955 | 22,47093 | 22,95893 | 21,84506 | 21,44935 | 22,75870 | 21,02823 |
| | Pre-mRNA-processing-splicing factor 8 | Q7L2E3 | PRPF8 | 0,47834 | -0,54496 | 24,56674 | 24,46725 | 25,27601 | 24,48824 | 25,89222 | 25,56442 |
| | PIN2/TERF1-interacting telomerase inhibitor 1 | Q71U36 | PINX1 | 0,47798 | -1,88467 | 15,50000 | 21,04700 | 18,86804 | 21,29480 | 20,45192 | 19,32232 |
| | Gem-associated protein 4 | Q9Y230 | GEMIN4 | 0,47708 | 2,66793 | 23,46089 | 22,60935 | 22,61268 | 15,50000 | 23,41183 | 21,76729 |
| | Cytoplasmic dynein 1 heavy chain 1 | Q9H0D6 | DYNC1H1 | 0,47674 | 1,24490 | 25,51204 | 24,84336 | 24,86868 | 23,14642 | 26,00404 | 22,33892 |
| | 60S ribosomal protein L18 | P56537 | RPL18 | 0,47655 | -3,50335 | 15,50000 | 23,15174 | 26,00776 | 24,00427 | 25,16062 | 26,00466 |
| | Interferon-induced, double-stranded RNA-activated protein kinase | Q9UQ80 | EIF2AK2 | 0,47632 | -2,26892 | 15,50000 | 20,56113 | 22,10348 | 20,57274 | 22,21481 | 22,18383 |
| | Probable ATP-dependent RNA helicase DDX28 | P11388 | DDX28 | 0,47516 | 1,48391 | 19,75378 | 19,33447 | 19,89224 | 15,50000 | 19,29390 | 19,73488 |
| | WD repeat-containing protein 48 | Q9NTJ3 | WDR48 | 0,47462 | -1,96414 | 15,50000 | 20,60820 | 20,76959 | 20,73703 | 20,20681 | 21,82637 |
| | Transcription factor A, mitochondrial | P24928 | TFAM | 0,47297 | -1,99462 | 15,50000 | 20,37146 | 21,33593 | 21,35019 | 20,49989 | 21,34115 |
| | ATP-binding cassette sub-family F member 1 | Q9Y4P3 | ABCF1 | 0,47256 | -0,46876 | 25,50237 | 25,09603 | 26,31259 | 25,66123 | 26,17727 | 26,47877 |
| | Splicing factor U2AF 35 kDa subunit;Splicing factor U2AF 26 kDa subunit | P47755 | U2AF1;U2AF1L4 | 0,47089 | -2,44622 | 15,50000 | 20,73612 | 22,91656 | 21,42101 | 22,03527 | 23,03507 |
| | Ras-associated and pleckstrin homology domains-containing protein 1 | Q3YEC7 | RAPH1 | 0,46803 | -1,26650 | 15,50000 | 19,04303 | 18,22322 | 19,78011 | 18,57633 | 18,20929 |
| | Cyclin-dependent-like kinase 5 | Q8TAQ2 | CDK5 | 0,46783 | 2,11726 | 21,61787 | 21,74997 | 21,12755 | 15,50000 | 20,99859 | 21,64500 |
| | THO complex subunit 1 | Q14103 | THOC1 | 0,46733 | -1,55551 | 22,44529 | 23,70651 | 25,13469 | 22,99524 | 27,08568 | 25,87211 |
| | Actin-related protein 3 | I3L0E3 | ACTR3 | 0,46589 | -0,30223 | 25,23338 | 25,16081 | 24,85534 | 25,27088 | 25,01125 | 25,87408 |
| | 60 kDa SS-A/Ro ribonucleoprotein | P04843 | TROVE2 | 0,46476 | -0,29910 | 21,53460 | 21,41214 | 21,55983 | 21,25373 | 22,10965 | 22,04049 |
| | Treacle protein | Q9HB71 | TCOF1 | 0,46461 | -1,06661 | 21,11902 | 24,16123 | 22,85306 | 24,68368 | 23,21798 | 23,43148 |
| | Flotillin-2 | Q9BVI4 | FLOT2 | 0,46435 | 0,45304 | 23,27980 | 22,65674 | 22,17813 | 22,29978 | 21,75156 | 22,70421 |
| | ATP-binding cassette sub-family B member 7, mitochondrial | P78527 | ABCB7 | 0,46395 | -1,81456 | 15,50000 | 20,75573 | 20,11223 | 20,45744 | 21,26132 | 20,09288 |
| | Probable ATP-dependent RNA helicase YTHDC2 | Q92922 | YTHDC2 | 0,46349 | 2,19760 | 21,66258 | 20,65377 | 22,57101 | 15,50000 | 21,10175 | 21,69281 |
| | Plastin-3 | O14617 | PLS3 | 0,46322 | 0,57869 | 22,14799 | 22,37989 | 21,85403 | 22,44306 | 20,64852 | 21,55426 |

| Significant | Protein names | UniProtID | Gene names | -LOG(P-value) | Difference | CXCR4_1 | CXCR4_2 | CXCR4_3 | MOCK_1 | MOCK_2 | MOCK_3 |
|---|---|---|---|---|---|---|---|---|---|---|---|
| | Histone H2B type 1-L;Histone H2B type 1-M;Histone H2B type 1-N;Histone H2B type 1-H;Histone H2B type 2-F;Histone H2B type 1-C/E/F/G/I;Histone H2B type 1-D;Histone H2B type F-S;Histone H2B type 1-K | O15355 | HIST1H2BL;HIST1H2BM;HIST1H2BN;HIST1H2BH;HIST2H2BF;HIST1H2BC;HIST1H2BD;H2BFS;HIST1H2BK | 0,46177 | -1,43611 | 21,74919 | 22,52967 | 25,54593 | 23,41596 | 24,98863 | 25,72853 |
| | Calcium homeostasis endoplasmic reticulum protein | Q9UM54 | CHERP | 0,46117 | -0,78555 | 24,09547 | 26,30947 | 26,22293 | 26,47633 | 26,06735 | 26,44085 |
| | 40S ribosomal protein S8 | P09543 | RPS8 | 0,46044 | -1,14008 | 23,87446 | 24,55215 | 27,07521 | 25,47166 | 26,47837 | 26,97202 |
| | 60S ribosomal protein L19 | Q9P258 | RPL19 | 0,45736 | -3,10855 | 15,50000 | 22,53830 | 25,21004 | 23,46936 | 24,06027 | 25,04435 |
| | DNA/RNA-binding protein KIN17 | O43615 | KIN | 0,45699 | -0,87441 | 19,99742 | 20,69498 | 21,63936 | 20,30396 | 22,27549 | 22,37555 |
| | LIM domain and actin-binding protein 1 | P14866 | LIMA1 | 0,45693 | -0,70603 | 22,97474 | 24,20650 | 24,24452 | 24,42873 | 23,66014 | 25,45497 |
| | 60S ribosomal export protein NMD3 | Q96EY1 | NMD3 | 0,45645 | -1,14171 | 15,50000 | 18,75055 | 18,40761 | 19,27831 | 18,18810 | 18,61687 |
| | Transcription elongation regulator 1 | P10155 | TCERG1 | 0,45630 | -1,21029 | 20,39069 | 20,75914 | 22,16079 | 20,49062 | 23,97386 | 22,47699 |
| | Peptidyl-prolyl cis-trans isomerase-like 3 | P61158 | PPIL3 | 0,45603 | -1,97795 | 15,50000 | 21,52526 | 20,48079 | 21,38290 | 20,73471 | 21,32229 |
| | Long-chain-fatty-acid--CoA ligase 3 | Q9BUJ2 | ACSL3 | 0,45575 | 0,69891 | 21,91936 | 23,18083 | 22,38885 | 21,19940 | 22,89460 | 21,29832 |
| | Pumilio domain-containing protein KIAA0020 | Q9H4M9 | KIAA0020 | 0,45499 | -3,10530 | 15,50000 | 23,71440 | 24,49747 | 24,37763 | 23,26284 | 25,38729 |
| | 60S ribosomal protein L23 | P61160 | RPL23 | 0,45444 | -0,49254 | 24,49869 | 24,09515 | 25,36302 | 24,58583 | 25,45234 | 25,39630 |
| | Bromodomain-containing protein 4 | Q96AG4 | BRD4 | 0,45385 | -2,06755 | 15,50000 | 21,47605 | 20,90749 | 22,16026 | 20,51007 | 21,41586 |
| | Cell division cycle 5-like protein | Q9Y3I0 | CDC5L | 0,45341 | -3,10649 | 15,50000 | 22,63843 | 24,07692 | 21,32328 | 25,63359 | 24,57796 |
| | Phenylalanine--tRNA ligase alpha subunit | Q9UBQ5 | FARSA | 0,45327 | 0,71816 | 23,06054 | 22,92122 | 23,39846 | 22,00041 | 23,71398 | 21,51133 |
| | RuvB-like 1 | Q07021 | RUVBL1 | 0,45078 | -0,43545 | 23,73397 | 23,93778 | 24,36038 | 23,97579 | 25,18219 | 24,18048 |
| | Nuclear pore complex protein Nup50 | P62424 | NUP50 | 0,45045 | 0,95534 | 18,69207 | 17,76059 | 17,83152 | 18,42394 | 17,49420 | 15,50000 |
| | Serine/arginine repetitive matrix protein 1 | P05388 | SRRM1 | 0,45042 | -2,43467 | 15,50000 | 21,77016 | 22,78929 | 22,36086 | 21,69183 | 23,31077 |
| | Rab11 family-interacting protein 1 | P51991 | RAB11FIP1 | 0,45019 | -1,07664 | 20,75434 | 23,46214 | 23,22245 | 22,63034 | 24,55701 | 23,48148 |
| | DNA replication licensing factor MCM4 | P51532 | MCM4 | 0,44954 | 0,98446 | 19,62487 | 22,28994 | 22,56071 | 20,50727 | 20,32817 | 20,68671 |
| | 1-phosphatidylinositol 4,5-bisphosphate phosphodiesterase beta-4 | P43243 | PLCB4 | 0,44919 | -1,39900 | 15,50000 | 19,46210 | 19,40572 | 19,85343 | 18,97306 | 19,73831 |
| | Glycogen synthase kinase-3 beta | Q9NZM4 | GSK3B | 0,44849 | 0,49407 | 20,28455 | 19,49303 | 19,08825 | 19,62044 | 18,53283 | 19,23034 |
| | Heterogeneous nuclear ribonucleoprotein U-like protein 1 | Q13435 | HNRNPUL1 | 0,44515 | -0,30386 | 25,14332 | 25,74714 | 25,65521 | 25,55385 | 25,63616 | 26,26724 |
| | THO complex subunit 6 homolog | Q9NVV4 | THOC6 | 0,44366 | -1,10823 | 24,21653 | 23,29768 | 24,68854 | 23,44481 | 26,88180 | 25,20083 |
| | Phenylalanine--tRNA ligase beta subunit | Q86XI2 | FARSB | 0,44331 | -1,32981 | 15,50000 | 19,18839 | 19,48211 | 19,37607 | 19,62669 | 19,15716 |
| | NADH dehydrogenase [ubiquinone] flavoprotein 1, mitochondrial | Q12905 | NDUFV1 | 0,43974 | 0,49191 | 22,46797 | 21,92738 | 22,49870 | 21,39210 | 22,69068 | 21,33554 |
| | Mitochondrial import receptor subunit TOM40 homolog | P05386 | TOMM40 | 0,43788 | 1,83007 | 21,42281 | 20,75344 | 20,09313 | 15,50000 | 21,26081 | 20,01835 |
| | AT-rich interactive domain-containing protein 2 | P61978 | ARID2 | 0,43727 | -0,90564 | 21,47100 | 20,46503 | 19,19573 | 21,28613 | 22,31310 | 20,24945 |
| | SWI/SNF-related matrix-associated actin-dependent regulator of chromatin subfamily B member 1 | O60264 | SMARCB1 | 0,43719 | -0,63457 | 22,87405 | 21,71423 | 20,92340 | 22,64434 | 22,80546 | 21,96557 |
| | Abl interactor 1 | Q15427 | ABI1 | 0,43717 | -1,50823 | 15,50000 | 20,04540 | 19,28864 | 20,07496 | 20,39498 | 18,88878 |
| | Brain-specific angiogenesis inhibitor 1-associated protein 2 | P22626 | BAIAP2 | 0,43692 | -1,90425 | 15,50000 | 20,04220 | 21,00840 | 19,27048 | 21,08881 | 21,90408 |
| | Mediator of RNA polymerase II transcription subunit 17 | P62913 | MED17 | 0,43671 | -2,86847 | 15,50000 | 24,24757 | 23,47296 | 24,21771 | 23,28546 | 24,32279 |
| | Phosphatidylinositol 4-phosphate 5-kinase type-1 alpha | O94906 | PIP5K1A | 0,43671 | -1,35301 | 15,50000 | 18,98915 | 19,66395 | 19,58382 | 18,79398 | 19,83433 |
| | DNA polymerase delta subunit 3 | Q14247 | POLD3 | 0,43504 | -1,48254 | 15,50000 | 20,01522 | 19,41870 | 20,28467 | 19,99051 | 19,10638 |
| | 60S ribosomal protein L6 | O75190 | RPL6 | 0,43464 | -1,43885 | 21,17035 | 22,86015 | 25,76398 | 23,82542 | 24,91027 | 25,37534 |
| | RRP12-like protein | Q8N163 | RRP12 | 0,43411 | -1,23429 | 20,70438 | 20,66787 | 22,65818 | 21,23210 | 24,58938 | 21,91181 |
| | Heterogeneous nuclear ribonucleoprotein K | P07196 | HNRNPK | 0,43293 | -0,36575 | 25,88138 | 25,66582 | 26,64966 | 26,49267 | 26,05134 | 26,75009 |
| | Dolichyl-diphosphooligosaccharide--protein glycosyltransferase subunit STT3A | P14649 | STT3A | 0,43267 | 0,56322 | 21,67603 | 21,68949 | 21,19129 | 20,93675 | 21,88740 | 20,04300 |
| | SRSF protein kinase 1 | O95793 | SRPK1 | 0,43210 | -3,17665 | 15,50000 | 24,53647 | 24,98859 | 24,54469 | 24,01878 | 25,99155 |
| | Fructose-bisphosphate aldolase A | Q92925 | ALDOA | 0,43085 | 0,68160 | 22,56973 | 21,14967 | 20,55112 | 20,95837 | 21,14725 | 20,12010 |
| | THO complex subunit 2 | Q07955 | THOC2 | 0,43026 | -1,12177 | 24,74124 | 24,14367 | 25,52999 | 24,24598 | 27,82905 | 25,70520 |
| | Programmed cell death protein 2-like | Q9NZ01 | PDCD2L | 0,42863 | 1,58441 | 20,29939 | 20,45131 | 19,93833 | 15,50000 | 20,24179 | 20,19400 |
| | Rho GTPase-activating protein 10 | Q9Y265 | ARHGAP10 | 0,42736 | -2,00069 | 15,50000 | 21,60562 | 20,60928 | 21,48714 | 20,00717 | 22,22267 |
| | Protein transport protein Sec23B | Q86UP2 | SEC23B | 0,42646 | 0,74102 | 20,92998 | 21,13558 | 19,46949 | 20,73422 | 18,92694 | 19,65085 |
| | WD repeat-containing protein 34 | Q92769 | WDR34 | 0,42570 | -1,23896 | 18,30915 | 19,89427 | 19,47029 | 18,41237 | 22,38627 | 20,59195 |
| | 60S ribosomal protein L3 | P61964 | RPL3 | 0,42556 | -1,06931 | 23,22350 | 24,27299 | 26,34262 | 24,56790 | 26,29539 | 26,18374 |
| | tRNA (cytosine(34)-C(5))-methyltransferase | P55795 | NSUN2 | 0,42503 | 0,68816 | 21,77816 | 23,41377 | 23,87849 | 21,86759 | 22,34055 | 22,79778 |
| | Actin-related protein 2/3 complex subunit 5-like protein | Q9UN81 | ARPC5L | 0,42411 | 1,84293 | 20,70776 | 21,33533 | 21,04753 | 15,50000 | 20,87972 | 21,18211 |
| | 28S ribosomal protein S35, mitochondrial | Q9UHX1 | MRPS35 | 0,42288 | -2,24243 | 15,50000 | 21,64720 | 21,36688 | 20,58646 | 23,84230 | 20,81261 |
| | Probable ATP-dependent RNA helicase DDX49 | P56192 | DDX49 | 0,42247 | -1,93650 | 15,50000 | 21,15986 | 21,30964 | 20,67625 | 22,04680 | 21,05596 |
| | Protein DEK | Q96RS0 | DEK | 0,42222 | -2,04907 | 21,08518 | 26,59662 | 26,98055 | 26,38471 | 25,89113 | 28,53371 |
| | Probable ATP-dependent RNA helicase DDX20 | P35249 | DDX20 | 0,42169 | 0,89696 | 23,08604 | 22,80882 | 23,15188 | 20,63795 | 23,74827 | 21,96965 |
| | Band 4.1-like protein 5 | Q96GM5 | EPB41L5 | 0,42075 | -2,25644 | 15,50000 | 22,61858 | 21,47398 | 23,11518 | 21,05159 | 22,19511 |
| | Heparan sulfate 2-O-sulfotransferase 1 | Q9NQ66 | HS2ST1 | 0,42006 | -0,79040 | 19,67166 | 20,95525 | 19,95787 | 20,19748 | 22,38416 | 20,37433 |
| | Coilin | Q6UXN9 | COIL | 0,41813 | -2,51432 | 15,50000 | 23,06431 | 22,83076 | 22,50870 | 22,20934 | 24,22000 |
| | AP-3 complex subunit sigma-1 | Q8NE71 | AP3S1 | 0,41700 | -2,07732 | 15,50000 | 22,26991 | 21,03106 | 22,48640 | 21,16521 | 21,38132 |
| | Splicing factor 1 | P09874 | SF1 | 0,41649 | -0,81215 | 19,76396 | 21,42193 | 22,37798 | 21,84770 | 21,52826 | 22,62435 |
| | Testis-expressed sequence 10 protein | Q8WVM8 | TEX10 | 0,41594 | -1,71485 | 21,04414 | 23,53481 | 24,33749 | 22,18386 | 27,19892 | 24,67820 |

| Significant | Protein names | UniProtID | Gene names | -LOG(P-value) | Difference | CXCR4_1 | CXCR4_2 | CXCR4_3 | MOCK_1 | MOCK_2 | MOCK_3 |
|---|---|---|---|---|---|---|---|---|---|---|---|
| | DNA-directed RNA polymerases I, II, and III subunit RPABC3 | Q02978 | POLR2H | 0,41513 | -2,14586 | 15,50000 | 22,44068 | 21,36437 | 22,67687 | 21,18338 | 21,88237 |
| | Protein RER1 | Q9NR50 | RER1 | 0,41360 | -2,31596 | 15,50000 | 22,78554 | 22,02520 | 21,70632 | 23,54439 | 22,00789 |
| | NADH-ubiquinone oxidoreductase 75 kDa subunit, mitochondrial | Q99623 | NDUFS1 | 0,41350 | 2,41558 | 22,23725 | 22,47093 | 23,66709 | 15,50000 | 23,11818 | 22,51034 |
| | 60S ribosomal protein L36 | P26196 | RPL36 | 0,41327 | -2,29866 | 21,10840 | 15,50000 | 23,34638 | 21,61984 | 22,28074 | 22,95020 |
| | Leucine-rich repeat-containing protein 47 | O60244 | LRRC47 | 0,41196 | -1,89981 | 15,50000 | 21,09803 | 21,60585 | 21,06677 | 21,48262 | 21,35391 |
| | Small nuclear ribonucleoprotein Sm D3 | P62829 | SNRPD3 | 0,41171 | -2,80750 | 23,43262 | 15,50000 | 24,47877 | 22,74357 | 24,46083 | 24,62951 |
| | Nucleolar RNA helicase 2 | Q9BQA1 | DDX21 | 0,41109 | -1,55692 | 20,06184 | 22,31645 | 24,71025 | 22,17085 | 24,52821 | 25,06023 |
| | DNA replication licensing factor MCM5 | Q15029 | MCM5 | 0,41080 | -1,10046 | 19,98008 | 21,47284 | 23,75338 | 22,67157 | 22,41372 | 23,42240 |
| | 60S ribosomal protein L17 | P60660 | RPL17-C18orf32;RPL17 | 0,41065 | -1,01285 | 21,68011 | 24,73903 | 24,41022 | 24,09869 | 25,40237 | 24,36686 |
| | Calcium/calmodulin-dependent protein kinase type II subunit delta | Q93074 | CAMK2D | 0,40845 | 0,48547 | 21,08304 | 21,01202 | 20,62489 | 20,90925 | 19,45312 | 20,90117 |
| | Ubiquitin carboxyl-terminal hydrolase isozyme L5 | Q92614 | UCHL5 | 0,40751 | 0,36099 | 20,67883 | 21,36025 | 20,58426 | 20,80711 | 20,79090 | 19,94235 |
| | Talin-1 | P40938 | TLN1 | 0,40736 | 0,34528 | 20,53155 | 20,35154 | 21,43120 | 20,35864 | 20,68372 | 20,23608 |
| | Eukaryotic translation initiation factor 2 subunit 3;Putative eukaryotic translation initiation factor 2 subunit 3-like protein | P35251 | EIF2S3;EIF2S3L | 0,40641 | -0,83724 | 23,47544 | 24,40341 | 25,67522 | 24,16883 | 25,82355 | 26,07341 |
| | LINE-1 retrotransposable element ORF1 protein | P53396 | L1RE1 | 0,40543 | -0,44702 | 20,11133 | 20,25223 | 20,47971 | 21,11216 | 19,82151 | 21,25067 |
| | Peptidyl-prolyl cis-trans isomerase B | P17844 | PPIB | 0,40540 | -1,29098 | 23,29039 | 21,81061 | 24,53321 | 22,66299 | 24,38528 | 26,45886 |
| | Methionine aminopeptidase 2 | P15880 | METAP2 | 0,40351 | -1,58036 | 15,50000 | 20,52917 | 20,12174 | 20,99424 | 20,25811 | 19,63964 |
| | 60S ribosomal protein L13a | E9PAV3 | RPL13A | 0,40334 | -0,60629 | 23,63079 | 24,46818 | 25,64526 | 24,81176 | 25,08091 | 25,67042 |
| | GTP-binding nuclear protein Ran | Q9BZK7 | RAN | 0,40274 | 0,69023 | 23,88920 | 24,20948 | 22,62948 | 23,95500 | 22,18873 | 22,51374 |
| | Leucine-rich repeat flightless-interacting protein 2 | O94973 | LRRFIP2 | 0,40080 | -1,82782 | 15,50000 | 20,39373 | 21,36682 | 21,06309 | 19,70929 | 21,97162 |
| | Protein RRP5 homolog | Q6P2Q9 | PDCD11 | 0,40076 | -0,85305 | 22,48139 | 19,98454 | 21,11731 | 21,44253 | 23,12354 | 21,57630 |
| | Nucleosome assembly protein 1-like 1 | Q92785 | NAP1L1 | 0,40031 | 0,89765 | 21,21898 | 21,95904 | 24,19520 | 20,92673 | 21,90364 | 21,84992 |
| | Transmembrane protein 33 | P18583 | TMEM33 | 0,39844 | 0,40717 | 22,56651 | 23,28616 | 23,78816 | 22,47117 | 23,28898 | 22,65916 |
| | 40S ribosomal protein S3 | Q12923 | RPS3 | 0,39815 | -0,74176 | 25,56628 | 25,63364 | 27,58039 | 26,15093 | 27,32260 | 27,53208 |
| | Mediator of RNA polymerase II transcription subunit 14 | Q16629 | MED14 | 0,39663 | -0,49016 | 22,65888 | 24,18646 | 23,67702 | 24,39159 | 23,48873 | 24,11251 |
| | Flap endonuclease 1 | O15145 | FEN1 | 0,39578 | -0,85589 | 23,18643 | 23,63920 | 25,04356 | 23,37955 | 25,36132 | 25,69602 |
| | DNA-directed RNA polymerases I, II, and III subunit RPABC1 | Q8TEQ6 | POLR2E | 0,39507 | -0,70907 | 19,84970 | 22,00398 | 21,54411 | 22,27512 | 21,08031 | 22,16956 |
| | 40S ribosomal protein S24 | Q9H501 | RPS24 | 0,39506 | -0,71080 | 21,26167 | 22,12146 | 23,50179 | 22,22653 | 23,36813 | 23,42265 |
| | EH domain-containing protein 4 | Q3MHD2 | EHD4 | 0,39436 | -0,75145 | 20,31529 | 21,90003 | 21,37814 | 21,85003 | 23,13181 | 20,86597 |
| | Glutathione S-transferase P | Q13310 | GSTP1 | 0,39287 | 1,27729 | 19,60948 | 19,29165 | 18,43774 | 20,09520 | 17,91181 | 15,50000 |
| | Breakpoint cluster region protein | Q9UL25 | BCR | 0,39123 | -1,64916 | 21,06815 | 15,50000 | 19,56162 | 20,27481 | 21,49067 | 19,31177 |
| | Nascent polypeptide-associated complex subunit alpha, muscle-specific form | Q02543 | NACA | 0,38900 | -0,53151 | 21,80475 | 20,03012 | 20,90925 | 20,92100 | 21,67861 | 21,73905 |
| | F-box-like/WD repeat-containing protein TBL1XR1 | O00425 | TBL1XR1 | 0,38891 | -0,53937 | 19,49207 | 20,66301 | 19,44291 | 21,25718 | 19,91606 | 20,04287 |
| | Neuroguidin | P35232 | NGDN | 0,38868 | -0,74032 | 20,19256 | 19,87007 | 20,86181 | 21,67827 | 19,55884 | 21,90830 |
| | Nuclear valosin-containing protein-like | P28288 | NVL | 0,38833 | -2,00998 | 15,50000 | 22,08398 | 21,94703 | 21,86257 | 22,19349 | 21,50489 |
| | Protein argonaute-2 | P42285 | AGO2 | 0,38643 | 1,69179 | 20,45724 | 21,10936 | 20,98057 | 15,50000 | 20,77000 | 21,20180 |
| | Polymerase delta-interacting protein 3 | P07355 | POLDIP3 | 0,38518 | -0,79856 | 22,01570 | 22,33811 | 23,74561 | 22,61908 | 23,01144 | 24,86457 |
| | FAST kinase domain-containing protein 2 | Q8NC51 | FASTKD2 | 0,38481 | -1,15538 | 15,50000 | 19,16091 | 19,29134 | 18,89968 | 19,59851 | 18,92021 |
| | LIM and calponin homology domains-containing protein 1 | P52429 | LIMCH1 | 0,38437 | 1,49465 | 20,43797 | 20,27674 | 20,06342 | 20,45543 | 15,50000 | 20,33876 |
| | U2 snRNP-associated SURP motif-containing protein | Q14008 | U2SURP | 0,38405 | -0,76851 | 24,59520 | 26,82736 | 27,28039 | 26,94022 | 26,79452 | 27,27373 |
| | Lamin-B2 | P30050 | LMNB2 | 0,38217 | 0,54025 | 20,19075 | 20,69268 | 20,61387 | 20,39613 | 18,82177 | 20,65866 |
| | Nucleophosmin | O60841 | NPM1 | 0,38123 | -0,78880 | 22,02722 | 23,45287 | 24,52964 | 24,04252 | 24,99632 | 23,33729 |
| | Serine/arginine-rich splicing factor 5 | P40429 | SRSF5 | 0,38098 | 2,27864 | 23,38311 | 20,43074 | 23,39401 | 15,50000 | 22,48456 | 22,38737 |
| | Serine/arginine repetitive matrix protein 2 | P62249 | SRRM2 | 0,38085 | -1,99433 | 15,50000 | 19,60684 | 22,54644 | 21,41792 | 19,70577 | 22,51258 |
| | 60S ribosomal protein L8 | Q9NX58 | RPL8 | 0,38079 | -2,62229 | 15,50000 | 22,71549 | 24,91780 | 22,61419 | 24,03339 | 24,35259 |
| | Histone H1.4 | Q9BVJ6 | HIST1H1E | 0,38058 | -1,18512 | 24,84297 | 25,48745 | 27,97010 | 26,79340 | 26,03836 | 29,02411 |
| | Ras-related protein Rab-14 | O15226 | RAB14 | 0,37995 | -1,79508 | 15,50000 | 20,78213 | 21,92358 | 21,41550 | 20,89297 | 21,28246 |
| | Arginine--tRNA ligase, cytoplasmic | P31943 | RARS | 0,37926 | -1,13499 | 22,64957 | 24,51569 | 24,15460 | 24,44550 | 27,03674 | 23,24258 |
| | 40S ribosomal protein S6 | Q96C57 | RPS6 | 0,37877 | -0,73201 | 23,91907 | 23,95677 | 25,80462 | 24,27412 | 25,60995 | 25,99242 |
| | Gem-associated protein 2 | Q14527 | GEMIN2 | 0,37642 | 1,49544 | 20,79066 | 19,93776 | 20,03456 | 15,50000 | 20,77676 | 19,99990 |
| | Ribonuclease P protein subunit p30 | P62937 | RPP30 | 0,37592 | -1,57286 | 15,50000 | 20,86944 | 20,24563 | 21,33767 | 19,86216 | 20,13383 |
| | Methylosome protein 50 | O75131 | WDR77 | 0,37537 | -0,49698 | 24,47203 | 25,61285 | 25,38634 | 25,29775 | 25,15039 | 26,51400 |
| | Signal recognition particle 54 kDa protein | Q12824 | SRP54 | 0,37445 | -0,97815 | 21,75218 | 22,07653 | 22,52217 | 21,69850 | 25,20247 | 22,38435 |
| | Mediator of RNA polymerase II transcription subunit 1 | Q14331 | MED1 | 0,37421 | -0,90406 | 21,05431 | 24,21712 | 23,43822 | 24,42547 | 23,22513 | 23,77122 |
| | Pre-mRNA-splicing factor ATP-dependent RNA helicase DHX15 | P11940 | DHX15 | 0,37395 | -1,17292 | 22,84430 | 25,41222 | 26,74227 | 25,10621 | 26,06850 | 27,34282 |
| | Kinesin-1 heavy chain | O75396 | KIF5B | 0,37372 | 2,15610 | 22,49599 | 21,54142 | 23,09420 | 15,50000 | 23,03867 | 22,12465 |
| | DNA-directed RNA polymerases I and III subunit RPAC1 | Q9NPJ6 | POLR1C | 0,37371 | 1,75381 | 21,25908 | 21,22917 | 20,98036 | 15,50000 | 21,75434 | 20,95284 |
| | FACT complex subunit SPT16 | Q02880 | SUPT16H | 0,37265 | -1,69012 | 20,86514 | 25,46516 | 26,58305 | 25,63572 | 24,92922 | 27,41877 |
| | Poly [ADP-ribose] polymerase 2 | P59998 | PARP2 | 0,37211 | -2,15766 | 15,50000 | 22,57059 | 22,46171 | 23,05303 | 21,02925 | 22,92299 |
| | Staphylococcal nuclease domain-containing protein 1 | P48047 | SND1 | 0,37088 | -1,04619 | 22,58068 | 21,27872 | 24,50585 | 22,42580 | 24,29544 | 24,78257 |
| | Cleft lip and palate transmembrane protein 1-like protein | Q92841 | CLPTM1L | 0,36993 | 1,32936 | 19,80934 | 20,02120 | 19,59752 | 15,50000 | 20,25200 | 19,68799 |
| | PRA1 family protein 3 | P26373 | ARL6IP5 | 0,36902 | -1,81058 | 15,50000 | 21,74119 | 21,51735 | 21,02432 | 21,58957 | 21,57639 |
| | Trinucleotide repeat-containing gene 6A protein | O75643 | TNRC6A | 0,36846 | 1,27106 | 23,49570 | 22,81886 | 20,70877 | 23,27753 | 20,65744 | 19,27517 |

| Significant | Protein names | UniProtID | Gene names | -LOG(P-value) | Difference | CXCR4_1 | CXCR4_2 | CXCR4_3 | MOCK_1 | MOCK_2 | MOCK_3 |
|---|---|---|---|---|---|---|---|---|---|---|---|
| | Protein FRG1 | Q9Y2W1 | FRG1 | 0,36757 | -0,63537 | 21,44784 | 22,43919 | 22,87641 | 22,10438 | 22,52860 | 24,03658 |
| | Mediator of RNA polymerase II transcription subunit 23 | Q15393 | MED23 | 0,36689 | -1,17278 | 15,50000 | 19,68839 | 19,10774 | 19,57777 | 18,74924 | 19,48744 |
| | 60S ribosomal protein L23a | Q07666 | RPL23A | 0,36448 | -2,68728 | 15,50000 | 23,63854 | 25,32059 | 23,21567 | 24,24540 | 25,05990 |
| | Ribosome-binding protein 1 | P50914 | RRBP1 | 0,36400 | -0,77329 | 21,61423 | 21,82310 | 22,71331 | 22,13499 | 24,45776 | 21,87778 |
| | Ras GTPase-activating-like protein IQGAP1 | Q9GZL7 | IQGAP1 | 0,36342 | -0,95252 | 23,52934 | 22,95656 | 24,85572 | 23,23576 | 26,46175 | 24,50167 |
| | 60S ribosomal protein L10 | P26599 | RPL10 | 0,36257 | -2,49325 | 15,50000 | 23,07428 | 24,26821 | 22,02523 | 24,91442 | 23,38258 |
| | Unconventional myosin-VI | Q15796 | MYO6 | 0,36250 | -0,27203 | 22,05059 | 21,91758 | 22,12263 | 21,77728 | 22,84159 | 22,28802 |
| | Survival motor neuron protein | Q53H12 | SMN1 | 0,36221 | 1,76305 | 21,06592 | 21,40468 | 21,47946 | 15,50000 | 21,59072 | 21,57020 |
| | Exosome complex component RRP42 | O96019 | EXOSC7 | 0,36069 | 1,57806 | 19,94321 | 19,97953 | 21,66544 | 15,50000 | 20,39005 | 20,96395 |
| | ADP/ATP translocase 3;ADP/ATP translocase 3, N-terminally processed | Q9ULW3 | SLC25A6 | 0,36037 | 0,60608 | 23,89523 | 23,92751 | 23,62500 | 21,87609 | 24,21327 | 23,54014 |
| | Pre-mRNA-processing factor 40 homolog A | Q86WJ1 | PRPF40A | 0,36018 | -2,70407 | 15,50000 | 24,96062 | 24,28016 | 25,21893 | 22,87421 | 24,75984 |
| | Histone-arginine methyltransferase CARM1 | Q9ULH0 | CARM1 | 0,36012 | 0,68037 | 20,99576 | 20,95844 | 19,87854 | 21,17719 | 18,76613 | 19,84830 |
| | Round spermatid basic protein 1-like protein | Q9UHB6 | RSBN1L | 0,35822 | -2,36641 | 15,50000 | 23,72151 | 22,90378 | 21,38221 | 23,68485 | 24,15746 |
| | Bcl-2-associated transcription factor 1 | P52597 | BCLAF1 | 0,35710 | -2,43119 | 15,50000 | 23,17035 | 24,37226 | 23,51785 | 22,44109 | 24,37723 |
| | Protein TBRG4 | P19388 | TBRG4 | 0,35614 | 0,56712 | 20,45443 | 19,99383 | 19,42263 | 19,45876 | 20,37867 | 18,33211 |
| | ELAV-like protein 1 | P47897 | ELAVL1 | 0,35609 | 2,29223 | 23,10851 | 22,42301 | 23,58187 | 15,50000 | 22,62140 | 24,11529 |
| | Calcineurin-binding protein cabin-1 | P08865 | CABIN1 | 0,35388 | -1,61957 | 19,94192 | 21,09359 | 15,50000 | 22,15295 | 19,55710 | 19,68418 |
| | Lysine-rich nucleolar protein 1 | P18124 | KNOP1 | 0,35340 | -1,03633 | 19,48408 | 22,78109 | 21,83677 | 23,60885 | 21,10322 | 22,49885 |
| | Acylglycerol kinase, mitochondrial | P62847 | AGK | 0,35337 | -0,69130 | 22,87378 | 22,90866 | 23,16864 | 22,59717 | 25,25663 | 23,17118 |
| | Probable ATP-dependent RNA helicase DDX5 | P19338 | DDX5 | 0,35306 | -0,52791 | 24,18828 | 25,11644 | 26,08881 | 25,30821 | 25,43011 | 26,23896 |
| | Nuclear cap-binding protein subunit 1 | Q9Y2R4 | NCBP1 | 0,35090 | -2,28366 | 15,50000 | 22,83150 | 23,95863 | 22,18444 | 22,96298 | 23,99369 |
| | ATP synthase subunit O, mitochondrial | P62280 | ATP5O | 0,34920 | -0,65101 | 22,50974 | 23,42085 | 24,68368 | 23,36226 | 24,29999 | 24,90504 |
| | Putative RNA polymerase II subunit B1 CTD phosphatase RPAP2 | Q6P2C8 | RPAP2 | 0,34917 | -1,77462 | 15,50000 | 22,26296 | 20,41318 | 21,74283 | 19,93876 | 21,81839 |
| | 60S ribosomal protein L12 | P35250 | RPL12 | 0,34910 | -0,60059 | 24,02673 | 24,03633 | 25,89583 | 24,55215 | 25,64763 | 25,56087 |
| | Actin-binding LIM protein 1 | Q96BJ3 | ABLIM1 | 0,34829 | -1,37875 | 15,50000 | 20,75678 | 19,50992 | 20,76401 | 19,30070 | 19,83824 |
| | DNA topoisomerase 2-beta | P82933 | TOP2B | 0,34613 | -0,64427 | 22,60123 | 20,00525 | 21,47956 | 21,67491 | 22,26376 | 22,08018 |
| | Casein kinase II subunit beta | P62753 | CSNK2B | 0,34364 | -1,68526 | 15,50000 | 19,99286 | 22,15515 | 20,57330 | 20,19003 | 21,94045 |
| | DNA methyltransferase 1-associated protein 1 | O75533 | DMAP1 | 0,34210 | 0,97439 | 19,07971 | 18,79785 | 18,53047 | 18,82842 | 19,15642 | 15,50000 |
| | Protein ENL | Q96D71 | MLLT1 | 0,34165 | -2,09808 | 15,50000 | 23,09941 | 22,40722 | 22,38047 | 21,15301 | 23,76739 |
| | RNA polymerase II-associated factor 1 homolog | Q8NEJ9 | PAF1 | 0,34122 | 1,83345 | 22,10390 | 21,36901 | 21,75405 | 15,50000 | 22,44253 | 21,78409 |
| | Lon protease homolog, mitochondrial | P23396 | LONP1 | 0,34077 | 1,73653 | 18,78692 | 21,33123 | 21,63321 | 15,50000 | 22,09552 | 18,94624 |
| | Mediator of RNA polymerase II transcription subunit 30 | P54198 | MED30 | 0,33870 | -1,86694 | 15,50000 | 22,55089 | 20,90110 | 23,01130 | 20,23163 | 21,30986 |
| | Sister chromatid cohesion protein PDS5 homolog B | P05387 | PDS5B | 0,33854 | -0,87516 | 21,80069 | 21,15382 | 21,98767 | 21,14289 | 24,55555 | 21,86922 |
| | X-ray repair cross-complementing protein 5 | Q9H223 | XRCC5 | 0,33841 | -3,21798 | 15,50000 | 25,15287 | 27,92547 | 24,88311 | 25,03175 | 28,31743 |
| | Plasminogen receptor (KT) | Q9Y676 | PLGRKT | 0,33806 | -1,45052 | 15,50000 | 20,33364 | 20,35380 | 20,83923 | 18,71877 | 20,98099 |
| | Filamin-A | Q99442 | FLNA | 0,33763 | 0,43959 | 25,98637 | 25,91670 | 25,29109 | 26,15979 | 25,25278 | 24,46283 |
| | E3 ubiquitin-protein ligase MYCBP2 | Q15291 | MYCBP2 | 0,33665 | -1,16608 | 21,46274 | 20,17329 | 20,81041 | 20,88733 | 24,72458 | 20,33276 |
| | MICOS complex subunit MIC19 | O75691 | CHCHD3 | 0,33660 | 0,60514 | 22,62250 | 23,52659 | 21,83329 | 21,80053 | 23,12458 | 21,24185 |
| | Tropomyosin alpha-3 chain | Q13895 | TPM3 | 0,33648 | -1,98489 | 15,50000 | 22,59662 | 22,34028 | 22,44983 | 20,74024 | 23,20152 |
| | HMG box transcription factor BBX | O75151 | BBX | 0,33635 | -1,34275 | 15,50000 | 20,90323 | 19,67062 | 20,04006 | 19,66990 | 20,39215 |
| | Transcriptional regulator ATRX | O15042 | ATRX | 0,33545 | -1,09968 | 19,46425 | 22,90097 | 23,40950 | 23,43593 | 21,94839 | 23,68945 |
| | Protein polybromo-1 | Q13111 | PBRM1 | 0,33349 | -1,15844 | 23,44910 | 20,52095 | 19,28636 | 22,74390 | 23,17536 | 20,81245 |
| | Ras-related protein Rab-18 | Q9P2E9 | RAB18 | 0,33241 | -1,36499 | 15,50000 | 20,98774 | 19,86742 | 20,64174 | 19,84486 | 19,96353 |
| | Ras-related protein Ral-A | Q8IWX8 | RALA | 0,33197 | 1,67205 | 20,42532 | 20,65097 | 22,23710 | 15,50000 | 20,80719 | 21,99006 |
| | Protein SON | Q9Y3D8 | SON | 0,33062 | -0,55021 | 22,18550 | 24,15831 | 23,56058 | 24,19370 | 23,13475 | 24,22656 |
| | Interleukin enhancer-binding factor 3 | P06748 | ILF3 | 0,33055 | -2,56363 | 15,50000 | 21,80648 | 25,61983 | 21,57163 | 23,26642 | 25,77914 |
| | SWI/SNF-related matrix-associated actin-dependent regulator of chromatin subfamily A member 5 | Q7LGA3 | SMARCA5 | 0,32964 | -0,36598 | 23,52683 | 23,76040 | 23,07821 | 24,61986 | 23,25465 | 23,58887 |
| | Heterogeneous nuclear ribonucleoprotein Q | Q9H0A0 | SYNCRIP | 0,32864 | -2,50330 | 15,50000 | 21,39530 | 25,46236 | 21,44167 | 22,86431 | 25,56157 |
| | SWI/SNF-related matrix-associated actin-dependent regulator of chromatin subfamily D member 2 | Q9Y383 | SMARCD2 | 0,32679 | -0,42566 | 22,29067 | 21,50978 | 20,98454 | 22,52724 | 22,25312 | 21,28161 |
| | U4/U6 small nuclear ribonucleoprotein Prp31 | O60524 | PRPF31 | 0,32641 | 1,58573 | 20,62952 | 20,33090 | 21,86034 | 15,50000 | 20,87635 | 21,68723 |
| | Chromodomain-helicase-DNA-binding protein 1 | Q16576 | CHD1 | 0,32272 | -1,93664 | 15,50000 | 22,83711 | 22,46949 | 22,77233 | 20,99141 | 22,85278 |
| | Unconventional myosin-IXb | Q9BY77 | MYO9B | 0,32247 | -2,08111 | 23,58909 | 15,50000 | 21,84441 | 22,08440 | 24,23396 | 20,85848 |
| | Oxysterol-binding protein-related protein 8 | Q15637 | OSBPL8 | 0,32114 | 0,69169 | 22,67487 | 21,66978 | 21,53602 | 20,27401 | 22,86650 | 20,66510 |
| | Protein SDA1 homolog | Q9H944 | SDAD1 | 0,32111 | -2,51505 | 15,50000 | 24,34881 | 25,01912 | 24,11251 | 22,49882 | 25,80174 |
| | Microsomal glutathione S-transferase 1 | Q9Y2T2 | MGST1 | 0,31947 | -1,01270 | 22,43125 | 21,94360 | 24,07300 | 21,66097 | 25,45513 | 24,36986 |
| | 1-phosphatidylinositol 4,5-bisphosphate phosphodiesterase beta-1 | O60293 | PLCB1 | 0,31942 | -0,46041 | 19,78673 | 21,01495 | 20,34960 | 21,47016 | 19,91890 | 21,14345 |
| | GTPase Era, mitochondrial | O95232 | ERAL1 | 0,31817 | 1,46507 | 20,94971 | 20,62934 | 20,53374 | 15,50000 | 21,47991 | 20,73769 |
| | Casein kinase I isoform epsilon | Q12874 | CSNK1E | 0,31336 | 1,31431 | 19,62115 | 19,89211 | 20,73355 | 15,50000 | 19,71671 | 21,08719 |
| | Serine/arginine-rich splicing factor 11 | Q9Y2X3 | SRSF11 | 0,31207 | -1,49932 | 19,43659 | 23,37584 | 25,21534 | 23,94548 | 22,61722 | 25,96303 |
| | Creatine kinase B-type | P22061 | CKB | 0,31140 | 0,45649 | 23,28687 | 21,84900 | 21,91269 | 22,63070 | 21,44096 | 21,60743 |
| | Small nuclear ribonucleoprotein Sm D2 | Q9UI10 | SNRPD2 | 0,31114 | -0,17096 | 22,28709 | 21,95248 | 21,55117 | 22,12717 | 22,20958 | 21,96688 |
| | Exonuclease 3-5 domain-containing protein 2 | P61247 | EXD2 | 0,31083 | -1,40839 | 15,50000 | 21,06769 | 20,99092 | 20,90522 | 20,45634 | 20,42224 |

| Significant | Protein names | UniProtID | Gene names | -LOG(P-value) | Difference | CXCR4_1 | CXCR4_2 | CXCR4_3 | MOCK_1 | MOCK_2 | MOCK_3 |
|---|---|---|---|---|---|---|---|---|---|---|---|
| | U1 small nuclear ribonucleoprotein 70 kDa | Q99714 | SNRNP70 | 0,30990 | -1,77576 | 15,50000 | 21,98912 | 22,81677 | 22,02073 | 21,22958 | 22,38287 |
| | Trinucleotide repeat-containing gene 6B protein | P41091 | TNRC6B | 0,30532 | 0,94240 | 26,11555 | 25,60961 | 23,34760 | 25,93193 | 23,15586 | 23,15778 |
| | 60S ribosomal protein L18a | Q9HAU5 | RPL18A | 0,30524 | -0,58731 | 21,97474 | 23,11218 | 24,41177 | 23,08105 | 23,96526 | 24,21430 |
| | Dimethylaniline monooxygenase [N-oxide-forming] 2 | Q12872 | FMO2 | 0,30480 | 1,70093 | 21,96215 | 21,83931 | 21,21478 | 15,50000 | 21,63130 | 22,78215 |
| | Nck-associated protein 1 | Q14690 | NCKAP1 | 0,30470 | -1,30106 | 15,50000 | 19,09587 | 19,32764 | 18,39268 | 21,68543 | 17,74859 |
| | RNA-binding protein 4 | P39748 | RBM4 | 0,30428 | 0,19570 | 22,80648 | 23,11740 | 22,95246 | 22,85232 | 22,29964 | 23,13729 |
| | 60S ribosomal protein L13 | Q9UHB9 | RPL13 | 0,30369 | -0,66383 | 23,58955 | 24,13743 | 26,01259 | 24,27177 | 25,50516 | 25,95414 |
| | Zinc finger CCCH domain-containing protein 18 | O43684 | ZC3H18 | 0,30358 | -1,53516 | 15,50000 | 21,59687 | 21,18241 | 21,83819 | 19,80273 | 21,24383 |
| | FACT complex subunit SSRP1 | P67809 | SSRP1 | 0,30349 | -1,50522 | 21,13477 | 25,83786 | 26,95238 | 24,90761 | 25,52943 | 28,00362 |
| | YTH domain-containing family protein 2 | Q99549 | YTHDF2 | 0,30344 | -1,04795 | 15,50000 | 19,42128 | 19,81359 | 19,52854 | 18,73691 | 19,61328 |
| | Serine/threonine-protein kinase RIO1 | P51398 | RIOK1 | 0,30270 | -1,28990 | 20,45232 | 25,34868 | 25,32553 | 24,60637 | 24,22413 | 26,16574 |
| | 60S ribosomal protein L4 | P82650 | RPL4 | 0,30247 | -0,92404 | 24,68110 | 25,04135 | 27,71581 | 25,17653 | 27,27950 | 27,75436 |
| | Filamin-B | Q14739 | FLNB | 0,30227 | 0,43699 | 22,73659 | 22,37462 | 21,64970 | 22,70282 | 20,99735 | 21,74977 |
| | Aspartate--tRNA ligase, cytoplasmic | O60870 | DARS | 0,30226 | -0,96523 | 24,61997 | 24,84709 | 24,31879 | 24,67309 | 28,10134 | 23,90711 |
| | SWI/SNF-related matrix-associated actin-dependent regulator of chromatin subfamily D member 1 | Q9NTI5 | SMARCD1 | 0,30207 | -0,45993 | 22,54538 | 23,89078 | 22,31066 | 24,08869 | 22,81540 | 23,22251 |
| | Serine/threonine-protein kinase PRP4 homolog | P16989 | PRPF4B | 0,30173 | -1,69673 | 15,50000 | 21,52440 | 22,41111 | 21,33827 | 20,36196 | 22,82547 |
| | Lamina-associated polypeptide 2, isoform alpha;Thymopoietin;Thymopentin | Q9UGU5 | TMPO | 0,30171 | 0,17712 | 21,99341 | 22,12758 | 22,47625 | 21,74004 | 22,38461 | 21,94124 |
| | Carboxymethylenebutenolidase homolog | Q8NFW8 | CMBL | 0,29939 | 1,35952 | 20,26396 | 21,03932 | 17,96268 | 21,00655 | 15,50000 | 18,68085 |
| | Succinate dehydrogenase [ubiquinone] flavoprotein subunit, mitochondrial | P35268 | SDHA | 0,29938 | -1,52815 | 15,50000 | 21,72030 | 21,20884 | 22,12177 | 20,27139 | 20,62042 |
| | Voltage-dependent anion-selective channel protein 2 | Q969G3 | VDAC2 | 0,29753 | 0,22559 | 25,39473 | 25,54578 | 25,23125 | 25,16054 | 25,67632 | 24,65813 |
| | Eukaryotic translation initiation factor 3 subunit J | Q15648 | EIF3J | 0,29644 | 1,73622 | 23,75797 | 19,57994 | 20,32948 | 15,50000 | 21,26648 | 21,69226 |
| | Phosphatidate cytidylyltransferase, mitochondrial | O94822 | TAMM41 | 0,29633 | 1,43449 | 20,75987 | 20,80625 | 20,89630 | 15,50000 | 21,80786 | 20,85110 |
| | Cyclin-dependent kinase 11A;Cyclin-dependent kinase 11B | Q68CP9 | CDK11A;CDK11B | 0,29432 | 0,39607 | 22,34679 | 21,76243 | 23,38179 | 22,00120 | 22,61097 | 21,69064 |
| | Cyclin-dependent kinase 9 | Q96KC8 | CDK9 | 0,29410 | -1,31771 | 15,50000 | 21,04893 | 18,66011 | 19,65877 | 18,29118 | 21,21223 |
| | Myosin light polypeptide 6 | O60287 | MYL6 | 0,29108 | -0,50929 | 25,68370 | 27,30287 | 27,10969 | 27,68545 | 26,22784 | 27,71083 |
| | Nipped-B-like protein | Q9Y6M1 | NIPBL | 0,28809 | 1,44467 | 21,82015 | 19,91230 | 20,09159 | 15,50000 | 22,05993 | 19,93010 |
| | HEAT repeat-containing protein 1;HEAT repeat-containing protein 1, N-terminally processed | O60831 | HEATR1 | 0,28776 | -0,97015 | 22,34519 | 21,84004 | 21,77398 | 21,75377 | 25,65067 | 21,46523 |
| | DnaJ homolog subfamily A member 3, mitochondrial | P36578 | DNAJA3 | 0,28755 | -0,29395 | 22,91216 | 23,27966 | 22,48306 | 22,90093 | 23,86788 | 22,78794 |
| | Regulator of nonsense transcripts 1 | Q96A33 | UPF1 | 0,28683 | 0,26182 | 23,00970 | 22,98968 | 23,87568 | 22,80225 | 22,80807 | 23,47927 |
| | Splicing regulatory glutamine/lysine-rich protein 1 | O43670 | SREK1 | 0,28572 | -2,09038 | 15,50000 | 22,79608 | 24,56500 | 22,50865 | 21,62729 | 24,99628 |
| | Multifunctional protein ADE2;Phosphoribosylaminoimidazole-succinocarboxamide synthase;Phosphoribosylaminoimidazole carboxylase | Q9UEW8 | PAICS | 0,28517 | 0,33717 | 22,77481 | 21,61648 | 21,62373 | 22,22738 | 21,44380 | 21,33232 |
| | Heat shock 70 kDa protein 1B;Heat shock 70 kDa protein 1A | Q96PK6 | HSPA1B;HSPA1A | 0,28385 | 0,27501 | 26,09772 | 26,69139 | 27,09607 | 26,09067 | 26,09271 | 26,87678 |
| | Tuftelin-interacting protein 11 | P46940 | TFIP11 | 0,28236 | -2,15493 | 15,50000 | 23,43250 | 24,12013 | 20,57358 | 23,89208 | 25,05175 |
| | H/ACA ribonucleoprotein complex subunit 2 | P14868 | NHP2 | 0,27878 | -1,02234 | 20,21810 | 23,94762 | 21,72376 | 24,89989 | 21,52373 | 22,53286 |
| | 60S ribosomal protein L7 | Q9H583 | RPL7 | 0,27875 | -0,70999 | 25,27170 | 24,74258 | 27,20247 | 25,04922 | 27,11515 | 27,18234 |
| | Serine/arginine-rich splicing factor 1 | P61011 | SRSF1 | 0,27751 | -0,42706 | 23,24731 | 22,50978 | 24,27824 | 23,09590 | 23,98650 | 24,23411 |
| | Dimethyladenosine transferase 1, mitochondrial | P49916 | TFB1M | 0,27478 | 1,22617 | 20,03322 | 20,43553 | 20,38459 | 15,50000 | 21,11610 | 20,55871 |
| | Serpin H1 | O95470 | SERPINH1 | 0,27455 | 0,29541 | 20,81425 | 21,71498 | 22,00316 | 21,67848 | 21,10111 | 20,86658 |
| | Eukaryotic translation initiation factor 3 subunit C;Eukaryotic translation initiation factor 3 subunit C-like protein | P38919 | EIF3C;EIF3CL | 0,27396 | 0,86656 | 28,49602 | 24,92379 | 24,68624 | 24,97334 | 25,76294 | 24,77012 |
| | Ras-related protein Rab-5C | Q96DI7 | RAB5C | 0,27256 | 0,29324 | 22,08868 | 23,10482 | 23,07267 | 22,91847 | 21,97278 | 22,49521 |
| | DAZ-associated protein 1 | Q9BQG0 | DAZAP1 | 0,27245 | 0,36032 | 22,06806 | 22,95709 | 23,16028 | 22,85114 | 21,55154 | 22,70178 |
| | Polypyrimidine tract-binding protein 1 | Q13601 | PTBP1 | 0,27237 | -0,68435 | 23,20528 | 24,05142 | 25,88651 | 24,18155 | 24,74689 | 26,26783 |
| | Mediator of RNA polymerase II transcription subunit 6 | P84077 | MED6 | 0,27130 | 1,41311 | 20,33484 | 21,80511 | 20,64685 | 15,50000 | 20,91736 | 22,13009 |
| | E2/E3 hybrid ubiquitin-protein ligase UBE2O | Q13573 | UBE2O | 0,27099 | 0,67510 | 23,44922 | 20,91867 | 21,32795 | 20,77402 | 22,44359 | 20,45292 |
| | Poly(rC)-binding protein 2 | P09661 | PCBP2 | 0,26801 | 0,47136 | 25,50513 | 25,78152 | 24,51334 | 25,93910 | 23,98155 | 24,46526 |
| | Paraspeckle component 1 | P10620 | PSPC1 | 0,26752 | 1,69920 | 20,64878 | 22,65550 | 22,88452 | 22,48498 | 15,50000 | 23,10621 |
| | Ras GTPase-activating protein-binding protein 2 | A0A0A6YYL6 | G3BP2 | 0,26551 | -1,41015 | 15,50000 | 20,12855 | 22,33587 | 19,40618 | 21,27418 | 21,51451 |
| | Protein NRDE2 homolog | Q9Y224 | NRDE2 | 0,26394 | -1,95619 | 15,50000 | 24,41584 | 22,85219 | 24,48971 | 20,79281 | 23,35407 |
| | PRA1 family protein 2 | Q9NX24 | PRAF2 | 0,26337 | -0,91510 | 15,50000 | 19,93229 | 18,73536 | 19,29437 | 19,46523 | 18,15333 |
| | Polyadenylate-binding protein 1 | O43172 | PABPC1 | 0,26318 | -0,63934 | 24,84498 | 26,44240 | 27,00121 | 25,29432 | 27,54642 | 27,36586 |
| | Insulin-like growth factor 2 mRNA-binding protein 1 | P68032 | IGF2BP1 | 0,26240 | -1,05067 | 20,58325 | 22,24127 | 25,18196 | 22,66405 | 23,06794 | 25,42650 |
| | Protein FAM98B | Q1ED39 | FAM98B | 0,26136 | 0,25647 | 21,71511 | 21,33789 | 21,32729 | 21,34803 | 20,50329 | 21,75955 |
| | AP-1 complex subunit mu-1 | Q13185 | AP1M1 | 0,26116 | 1,29313 | 20,42972 | 21,08771 | 20,63529 | 15,50000 | 21,52745 | 21,24586 |
| | Nuclear fragile X mental retardation-interacting protein 2 | Q7KZF4 | NUFIP2 | 0,26103 | 1,40907 | 20,23806 | 21,61144 | 21,48311 | 15,50000 | 21,54165 | 22,06375 |
| | Protein arginine N-methyltransferase 1 | Q9Y5A9 | PRMT1 | 0,26086 | -0,18161 | 23,32828 | 23,91762 | 24,09241 | 23,70504 | 23,94253 | 24,23557 |
| | Ras GTPase-activating protein-binding protein 1 | Q9NZI8 | G3BP1 | 0,25847 | -1,82050 | 15,50000 | 20,83345 | 24,37921 | 20,33789 | 21,74370 | 24,09257 |
| | STE20/SPS1-related proline-alanine-rich protein kinase | P51114 | STK39 | 0,25508 | -0,94671 | 15,50000 | 20,14768 | 19,35791 | 19,85162 | 19,30519 | 18,68890 |
| | RNA polymerase-associated protein RTF1 homolog | Q92665 | RTF1 | 0,25452 | -1,77798 | 15,50000 | 23,24436 | 23,08963 | 23,03409 | 20,28275 | 23,85110 |
| | Nucleolar protein 58 | Q03701 | NOP58 | 0,25303 | -0,82366 | 23,88418 | 27,42358 | 26,05459 | 27,65928 | 25,08817 | 27,08588 |
| | Tyrosine-protein phosphatase non-receptor type 13 | Q13428 | PTPN13 | 0,25251 | -0,55108 | 20,80719 | 23,17075 | 21,68910 | 23,21275 | 21,44127 | 22,66629 |

| Significant | Protein names | UniProtID | Gene names | -LOG(P-value) | Difference | CXCR4_1 | CXCR4_2 | CXCR4_3 | MOCK_1 | MOCK_2 | MOCK_3 |
|---|---|---|---|---|---|---|---|---|---|---|---|
| | Angiomotin | P39023 | AMOT | 0,25224 | 0,44029 | 24,49644 | 24,32327 | 23,62433 | 24,61549 | 22,47454 | 24,03313 |
| | Double-stranded RNA-binding protein Staufen homolog 1 | Q6WKZ4 | STAU1 | 0,25206 | -0,41861 | 21,10418 | 20,71934 | 22,72750 | 21,75169 | 21,65268 | 22,40247 |
| | 60S acidic ribosomal protein P1 | Q15051 | RPLP1 | 0,25069 | -0,35683 | 23,22831 | 24,12873 | 24,53202 | 25,14398 | 23,90003 | 23,91552 |
| | Delta(24)-sterol reductase | P19404 | DHCR24 | 0,25048 | 1,11909 | 20,10993 | 19,93445 | 19,74041 | 15,50000 | 21,54076 | 19,38678 |
| | X-ray repair cross-complementing protein 6 | O95782 | XRCC6 | 0,25022 | -1,28166 | 22,28740 | 25,66590 | 28,39873 | 25,90095 | 25,57632 | 28,71973 |
| | Neurofilament medium polypeptide | Q9H3K2 | NEFM | 0,24975 | 0,44363 | 24,97482 | 25,16557 | 24,26456 | 25,61866 | 23,46613 | 23,98928 |
| | Matrin-3 | P46100 | MATR3 | 0,24968 | -0,34309 | 23,81870 | 23,11580 | 24,67616 | 23,64438 | 24,30904 | 24,68651 |
| | Membrane-associated progesterone receptor component 1 | P07910 | PGRMC1 | 0,24857 | -1,18590 | 15,50000 | 20,85575 | 21,37925 | 20,57699 | 20,67685 | 20,03885 |
| | Developmentally-regulated GTP-binding protein 1 | P33992 | DRG1 | 0,24754 | -1,32569 | 18,06366 | 20,52564 | 23,22532 | 18,91777 | 23,41041 | 23,46351 |
| | Leucine-rich repeat-containing protein 40 | Q06787 | LRRC40 | 0,24709 | -1,98921 | 15,50000 | 25,14736 | 24,32347 | 24,23535 | 24,60524 | 22,09787 |
| | Signal recognition particle subunit SRP68 | P37108 | SRP68 | 0,24533 | -0,85706 | 20,90117 | 23,05121 | 23,00436 | 21,39346 | 25,41967 | 22,71479 |
| | Histone deacetylase 2 | Q9Y6A4 | HDAC2 | 0,24404 | -0,43700 | 21,20455 | 21,47724 | 21,56774 | 20,80829 | 23,17963 | 21,57261 |
| | HMG domain-containing protein 4 | Q86W42 | HMGXB4 | 0,24219 | -0,87672 | 19,62553 | 23,20577 | 20,87837 | 23,92488 | 20,60856 | 21,80640 |
| | Neurofilament light polypeptide | Q8NI27 | NEFL | 0,24008 | -0,41348 | 23,23929 | 24,60953 | 23,43326 | 25,21582 | 23,51665 | 23,79005 |
| | Kinectin | Q09028 | KTN1 | 0,23852 | -0,43580 | 23,00982 | 23,39519 | 23,87258 | 22,98383 | 25,18843 | 23,41274 |
| | DNA ligase 3 | P05198 | LIG3 | 0,23701 | -0,98925 | 19,36211 | 22,94457 | 24,01895 | 21,48861 | 23,45149 | 24,35326 |
| | 40S ribosomal protein S19 | P54136 | RPS19 | 0,23695 | -1,90845 | 22,16656 | 15,50000 | 25,35468 | 20,68517 | 22,97726 | 25,08416 |
| | Mediator of RNA polymerase II transcription subunit 26 | Q13325 | MED26 | 0,23527 | -1,49157 | 15,50000 | 23,53172 | 21,40374 | 21,86128 | 20,39362 | 22,65528 |
| | Thyroid hormone receptor-associated protein 3 | P62241 | THRAP3 | 0,23410 | -0,66750 | 22,17173 | 23,42944 | 24,87277 | 23,93490 | 22,89247 | 25,64905 |
| | Glutamine--tRNA ligase | Q96D46 | QARS | 0,23297 | -0,70941 | 23,53077 | 24,80331 | 24,46712 | 24,05432 | 27,23050 | 23,64460 |
| | MICOS complex subunit MIC60 | P62081 | IMMT | 0,23261 | 0,57155 | 23,66438 | 24,91105 | 25,00770 | 22,59903 | 25,55546 | 23,71398 |
| | Plasminogen activator inhibitor 1 RNA-binding protein | Q92552 | SERBP1 | 0,23131 | -0,59617 | 21,19382 | 22,32042 | 24,25314 | 22,47828 | 22,99386 | 24,08375 |
| | Translation initiation factor eIF-2B subunit beta | Q9NYY8 | EIF2B2 | 0,23058 | 1,19950 | 19,55469 | 20,83121 | 20,82042 | 15,50000 | 22,34186 | 19,76597 |
| | E3 ubiquitin-protein ligase HERC2 | Q86U86 | HERC2 | 0,22837 | 0,78824 | 24,21608 | 20,66614 | 21,32937 | 20,33930 | 22,87228 | 20,63529 |
| | RNA-binding protein 25 | O75592 | RBM25 | 0,22793 | -1,24866 | 19,89353 | 26,17722 | 25,06706 | 26,23153 | 23,16789 | 25,48438 |
| | Mitochondrial carrier homolog 2 | Q96EY4 | MTCH2 | 0,22727 | -0,18402 | 22,11867 | 22,20446 | 22,87648 | 22,42719 | 22,99478 | 22,32970 |
| | Bromodomain-containing protein 9 | Q9H8M2 | BRD9 | 0,22643 | -1,16976 | 15,50000 | 21,87237 | 20,20920 | 20,24458 | 19,27662 | 21,56964 |
| | Long-chain fatty acid transport protein 4 | Q5T3I0 | SLC27A4 | 0,22635 | 0,23330 | 21,74111 | 22,76401 | 22,33418 | 22,10399 | 22,48810 | 21,54731 |
| | Eukaryotic translation initiation factor 3 subunit B | Q9ULK4 | EIF3B | 0,22572 | 0,98574 | 29,09245 | 24,03440 | 24,65223 | 24,09233 | 26,13518 | 24,59434 |
| | Insulin-like growth factor 2 mRNA-binding protein 3 | O43143 | IGF2BP3 | 0,22536 | -0,58977 | 20,68808 | 21,92445 | 23,63777 | 21,73707 | 22,60413 | 23,67842 |
| | Hepatoma-derived growth factor-related protein 2 | Q96CW1 | HDGFRP2 | 0,22474 | -1,74449 | 15,50000 | 23,74745 | 23,30897 | 24,31251 | 19,77308 | 23,70430 |
| | RNA-binding protein FUS | P78356 | FUS | 0,22351 | 0,37994 | 27,37468 | 27,45803 | 26,98261 | 27,79749 | 25,63755 | 27,24045 |
| | Metaxin-2 | Q13576 | MTX2 | 0,22336 | 0,19322 | 22,07205 | 22,36941 | 22,24272 | 21,86393 | 22,66629 | 21,57432 |
| | NADH dehydrogenase [ubiquinone] flavoprotein 2, mitochondrial | P10412 | NDUFV2 | 0,22105 | -1,08912 | 15,50000 | 20,89785 | 21,40940 | 19,73461 | 20,91991 | 20,42008 |
| | DNA replication licensing factor MCM7 | O00264 | MCM7 | 0,21993 | 0,39769 | 22,52344 | 24,33354 | 24,69912 | 23,04467 | 23,57831 | 23,74006 |
| | AP-3 complex subunit beta-1 | Q9H8G2 | AP3B1 | 0,21967 | -1,40589 | 19,74141 | 25,92255 | 26,09083 | 25,77382 | 22,77290 | 27,42575 |
| | Calcyclin-binding protein | O14776 | CACYBP | 0,21905 | -0,25988 | 22,24153 | 22,77628 | 22,02283 | 22,55787 | 23,33019 | 21,93222 |
| | Eukaryotic translation initiation factor 3 subunit I | Q14839 | EIF3I | 0,21854 | 1,02527 | 27,75449 | 22,62346 | 23,49351 | 22,13912 | 25,26317 | 23,39336 |
| | Eukaryotic translation initiation factor 6 | Q5JTH9 | EIF6 | 0,21833 | -0,22747 | 22,04810 | 21,43227 | 21,68320 | 22,63498 | 21,39462 | 21,81640 |
| | Proteasome activator complex subunit 4 | Q96EX3 | PSME4 | 0,21828 | 1,55101 | 19,92519 | 19,77517 | 19,75026 | 15,50000 | 23,79759 | 15,50000 |
| | 60S acidic ribosomal protein P2 | O14744 | RPLP2 | 0,21819 | -0,74743 | 22,94733 | 22,83152 | 26,20792 | 23,25638 | 25,54998 | 25,42268 |
| | Constitutive coactivator of PPAR-gamma-like protein 1 | Q9Y5T5 | FAM120A | 0,21697 | 0,23138 | 21,63790 | 21,34657 | 22,52991 | 21,28980 | 21,51845 | 22,01199 |
| | 60S ribosomal protein L14 | P49756 | RPL14 | 0,21681 | -0,68027 | 22,59971 | 23,41893 | 26,03999 | 23,81421 | 24,33545 | 25,94978 |
| | Growth hormone-inducible transmembrane protein | Q13503 | GHITM | 0,21635 | -1,09402 | 15,50000 | 21,76842 | 20,85901 | 20,09674 | 20,81198 | 20,50077 |
| | DNA topoisomerase 2-alpha | Q70E73 | TOP2A | 0,21560 | -0,22910 | 24,33940 | 23,16153 | 23,63212 | 24,01035 | 23,50918 | 24,30083 |
| | Serine/threonine-protein kinase MRCK alpha | Q7L0Y3 | CDC42BPA | 0,21487 | 1,08099 | 20,65770 | 20,07091 | 20,30508 | 15,50000 | 22,01352 | 20,27719 |
| | Magnesium transporter protein 1 | P12956 | MAGT1 | 0,21482 | 1,17667 | 20,95816 | 19,97169 | 19,69513 | 15,50000 | 22,74520 | 18,84977 |
| | Supervillin | Q9BQ67 | SVIL | 0,21161 | 0,41642 | 24,11378 | 23,65731 | 22,89408 | 24,15097 | 21,85855 | 23,40639 |
| | ADP-ribosylation factor 1;ADP-ribosylation factor 3 | P46776 | ARF1;ARF3 | 0,20880 | -1,00598 | 15,50000 | 21,35971 | 20,52420 | 20,27662 | 20,67685 | 19,44839 |
| | Very-long-chain enoyl-CoA reductase | Q9BRS2 | TECR | 0,20840 | -0,43385 | 23,18132 | 22,43066 | 22,84496 | 21,85011 | 24,52952 | 23,37888 |
| | Cell division control protein 6 homolog | P23284 | CDC6 | 0,20751 | 0,60199 | 18,40528 | 18,48911 | 18,09435 | 18,72297 | 18,95980 | 15,50000 |
| | Glutamine--fructose-6-phosphate aminotransferase [isomerizing] 1 | P36542 | GFPT1 | 0,20725 | 0,66135 | 20,72584 | 23,82794 | 23,60513 | 20,71364 | 23,20071 | 22,26049 |
| | Serine/threonine-protein phosphatase PP1-beta catalytic subunit | Q6I9Y2 | PPP1CB | 0,20721 | -0,12414 | 23,01805 | 22,98111 | 23,49020 | 23,61482 | 23,13149 | 23,11548 |
| | Translational activator GCN1 | Q9Y2A7 | GCN1L1 | 0,20704 | 0,76873 | 23,43377 | 22,70698 | 22,61187 | 20,70126 | 24,97325 | 20,77193 |
| | 78 kDa glucose-regulated protein | Q8TEW0 | HSPA5 | 0,20689 | 0,35120 | 23,46613 | 25,54442 | 24,90362 | 23,85443 | 24,36185 | 24,64427 |
| | AT-rich interactive domain-containing protein 1A | Q96EY7 | ARID1A | 0,20668 | 0,58747 | 21,47264 | 20,80373 | 19,64241 | 21,23532 | 20,77072 | 18,15031 |
| | Adenylate kinase isoenzyme 6 | Q96I25 | AK6 | 0,20584 | -0,78627 | 15,50000 | 20,37316 | 19,26879 | 19,26323 | 19,17165 | 19,06587 |
| | Serine/threonine-protein kinase PLK1 | P25490 | PLK1 | 0,20574 | -0,20388 | 20,37846 | 21,31805 | 20,89401 | 21,46034 | 20,55028 | 21,19153 |
| | Phosphatidylinositol 5-phosphate 4-kinase type-2 alpha | Q9Y2R9 | PIP4K2A | 0,20561 | 0,96974 | 20,13609 | 20,27481 | 19,85552 | 15,50000 | 21,24016 | 20,61702 |
| | Lupus La protein | P50750 | SSB | 0,20508 | 1,34145 | 18,99037 | 18,92199 | 23,70641 | 15,50000 | 19,87388 | 22,22055 |
| | Mediator of RNA polymerase II transcription subunit 21 | Q9UPT8 | MED21 | 0,20490 | -1,25364 | 15,50000 | 23,01236 | 20,02810 | 22,57145 | 19,77944 | 19,95049 |
| | Protein CMSS1 | Q9Y5S9 | CMSS1 | 0,20428 | 0,96030 | 20,45573 | 19,24551 | 20,30719 | 15,50000 | 21,00498 | 20,62257 |
| | Vimentin | Q9Y295 | VIM | 0,20404 | 0,30340 | 25,41064 | 25,33316 | 24,31181 | 25,59278 | 24,08950 | 24,46314 |
| | Eukaryotic translation initiation factor 5B | O94766 | EIF5B | 0,20279 | -0,60202 | 26,41646 | 29,13095 | 28,95983 | 29,54725 | 27,30042 | 29,46564 |

| Significant | Protein names | UniProtID | Gene names | -LOG(P-value) | Difference | CXCR4_1 | CXCR4_2 | CXCR4_3 | MOCK_1 | MOCK_2 | MOCK_3 |
|---|---|---|---|---|---|---|---|---|---|---|---|
| | Actin-related protein 2 | Q9NWT1 | ACTR2 | 0,20163 | -0,31502 | 24,27668 | 24,71602 | 23,53362 | 25,14133 | 23,52205 | 24,80799 |
| | Gem-associated protein 5 | Q9NSD9 | GEMIN5 | 0,20158 | -0,55340 | 21,45689 | 22,06520 | 21,75062 | 21,51326 | 24,38027 | 21,03939 |
| | Complement component 1 Q subcomponent-binding protein, mitochondrial | P62244 | C1QBP | 0,20025 | -0,32983 | 22,19442 | 22,03086 | 23,35030 | 22,14376 | 23,76820 | 22,65312 |
| | Eukaryotic translation initiation factor 5 | Q8WY36 | EIF5 | 0,20024 | 1,10674 | 20,67633 | 20,63724 | 21,02154 | 15,50000 | 22,46451 | 21,05039 |
| | NF-kappa-B-repressing factor | Q9BZI7 | NKRF | 0,19995 | -0,62039 | 19,49906 | 21,73744 | 22,70582 | 20,59851 | 22,11302 | 23,09194 |
| | Src substrate cortactin | Q99755 | CTTN | 0,19945 | -0,38976 | 20,13119 | 21,74505 | 20,57469 | 21,89416 | 20,05775 | 21,66830 |
| | Leucine-rich repeat-containing protein 59 | Q9Y3B4 | LRRC59 | 0,19867 | -0,31740 | 27,22977 | 27,52903 | 28,63803 | 27,56289 | 27,79829 | 28,98785 |
| | Kinase D-interacting substrate of 220 kDa | Q9NP72 | KIDINS220 | 0,19853 | -0,69936 | 20,56913 | 19,50554 | 20,29177 | 19,43770 | 23,45538 | 19,57143 |
| | Tropomodulin-3 | O14639 | TMOD3 | 0,19708 | 0,33912 | 22,54307 | 22,48910 | 21,75467 | 22,46136 | 20,70455 | 22,60356 |
| | Cell growth-regulating nucleolar protein | Q8IWS0 | LYAR | 0,19688 | -0,61808 | 25,52453 | 27,88443 | 27,02189 | 27,93005 | 25,51794 | 28,83711 |
| | UV excision repair protein RAD23 homolog B | Q9BYN8 | RAD23B | 0,19532 | 0,41219 | 19,73724 | 19,05674 | 17,72227 | 19,50704 | 18,09620 | 17,67645 |
| | Myosin-14 | Q9BVP2 | MYH14 | 0,19422 | 0,69953 | 23,57762 | 25,56401 | 25,40931 | 25,06077 | 21,72355 | 25,66804 |
| | Eukaryotic translation initiation factor 3 subunit M | Q9UHB7 | EIF3M | 0,19257 | 0,74367 | 27,65745 | 23,98129 | 23,74714 | 23,27625 | 25,85804 | 24,02056 |
| | AF4/FMR2 family member 4 | P60866 | AFF4 | 0,19183 | -1,38765 | 15,50000 | 23,37372 | 22,38256 | 23,44190 | 19,35973 | 22,61758 |
| | Actin, alpha cardiac muscle 1;Actin, aortic smooth muscle;Actin, gamma-enteric smooth muscle;Actin, alpha skeletal muscle | Q3B726 | ACTC1;ACTA2;ACTG2;ACTA1 | 0,19040 | -1,03242 | 22,06016 | 26,52075 | 26,94592 | 23,70757 | 26,50191 | 28,41462 |
| | Uncharacterized protein C12orf43 | Q6P1X5 | C12orf43 | 0,18878 | -0,62353 | 20,25718 | 22,76871 | 21,83526 | 22,22767 | 20,47001 | 24,03406 |
| | ATP-dependent RNA helicase DDX39A | Q15147 | DDX39A | 0,18777 | 0,48777 | 20,87927 | 22,73314 | 20,98015 | 22,58461 | 20,57015 | 19,97449 |
| | Mediator of RNA polymerase II transcription subunit 12 | Q9H0S4 | MED12 | 0,18742 | -0,50974 | 21,57182 | 20,11312 | 20,32553 | 19,70730 | 22,91276 | 20,91962 |
| | Heterogeneous nuclear ribonucleoprotein L | O60832 | HNRNPL | 0,18588 | -0,29258 | 23,05489 | 23,29922 | 24,33368 | 23,89680 | 23,04629 | 24,62243 |
| | Ras-related protein Rab-11B | O00203 | RAB11B | 0,18543 | 0,15574 | 22,23322 | 23,28023 | 22,62916 | 22,46792 | 22,75416 | 22,45330 |
| | Elongation factor 1-alpha 1;Putative elongation factor 1-alpha-like 3 | Q9NVH0 | EEF1A1;EEF1A1P5 | 0,18375 | 0,11925 | 27,17741 | 27,32071 | 27,93961 | 27,20387 | 27,40202 | 27,47406 |
| | Heat shock cognate 71 kDa protein | Q9UN86 | HSPA8 | 0,18362 | 0,17183 | 26,73660 | 27,38574 | 27,77881 | 26,77944 | 27,18754 | 27,41869 |
| | Serine/arginine-rich splicing factor 7 | Q5T280 | SRSF7 | 0,18222 | -0,55196 | 23,21290 | 21,53602 | 24,75292 | 22,45189 | 23,90573 | 24,80011 |
| | Spectrin alpha chain, non-erythrocytic 1 | Q14498 | SPTAN1 | 0,18211 | 0,23301 | 23,26170 | 22,37870 | 22,16192 | 22,93251 | 21,71963 | 22,45116 |
| | Translation initiation factor eIF-2B subunit gamma | Q99880 | EIF2B3 | 0,18065 | -0,48924 | 21,72121 | 21,89914 | 21,85232 | 21,67526 | 24,32718 | 20,93797 |
| | Eukaryotic translation initiation factor 3 subunit A | Q02878 | EIF3A | 0,17967 | 0,73887 | 29,17234 | 24,67082 | 25,33929 | 24,55906 | 26,93405 | 25,47274 |
| | 40S ribosomal protein S2 | P63010 | RPS2 | 0,17940 | -0,52804 | 24,04185 | 24,14779 | 26,74343 | 24,23922 | 25,67807 | 26,59989 |
| | 40S ribosomal protein S16 | Q92759 | RPS16 | 0,17898 | -0,61789 | 23,97989 | 23,50422 | 26,41809 | 23,35111 | 26,09444 | 26,31032 |
| | Splicing factor 3B subunit 2 | Q9HBL7 | SF3B2 | 0,17883 | -0,34999 | 23,59912 | 23,65524 | 25,21908 | 23,50240 | 24,77374 | 25,24728 |
| | Fatty aldehyde dehydrogenase | Q8WTT2 | ALDH3A2 | 0,17833 | 0,23448 | 21,15128 | 21,71456 | 21,04240 | 20,56412 | 21,97446 | 20,66622 |
| | Poly(U)-binding-splicing factor PUF60 | P20042 | PUF60 | 0,17820 | -0,44740 | 26,22205 | 27,48984 | 28,15952 | 27,49595 | 26,54847 | 29,16919 |
| | Trimethylguanosine synthase | Q99470 | TGS1 | 0,17797 | -0,45388 | 20,73892 | 21,12666 | 20,18277 | 19,94106 | 22,96652 | 20,50242 |
| | Cytoskeleton-associated protein 5 | P62854 | CKAP5 | 0,17707 | -0,59807 | 22,92441 | 22,79947 | 22,14976 | 22,41013 | 25,69384 | 21,56388 |
| | Protein transport protein Sec23A | Q8TED0 | SEC23A | 0,17697 | 0,34728 | 21,85022 | 22,08456 | 21,35536 | 22,74160 | 20,29692 | 21,20979 |
| | Hsp90 co-chaperone Cdc37;Hsp90 co-chaperone Cdc37, N-terminally processed | Q15054 | CDC37 | 0,17343 | 0,24856 | 18,80483 | 20,08822 | 20,15906 | 20,06039 | 19,02364 | 19,22242 |
| | Double-stranded RNA-binding protein Staufen homolog 2 | O95402 | STAU2 | 0,17315 | 0,72763 | 19,37176 | 18,87634 | 19,83576 | 15,50000 | 20,22317 | 20,17779 |
| | Protein PRRC2B | Q05519 | PRRC2B | 0,17224 | 0,65982 | 23,64746 | 19,47596 | 19,82792 | 19,82100 | 21,44859 | 19,70230 |
| | Heterogeneous nuclear ribonucleoprotein A3 | Q08945 | HNRNPA3 | 0,17110 | -0,33968 | 22,59115 | 22,42642 | 24,23389 | 23,32978 | 22,64253 | 24,29817 |
| | DNA topoisomerase 1 | Q8IZP0 | TOP1 | 0,17076 | -1,55464 | 15,50000 | 22,31493 | 24,41235 | 20,80978 | 19,55266 | 26,52876 |
| | Drebrin | Q99878 | DBN1 | 0,16956 | 0,30635 | 26,25712 | 25,44177 | 25,58781 | 25,99449 | 24,19114 | 26,18200 |
| | Target of EGR1 protein 1 | Q9UMS4 | TOE1 | 0,16925 | -0,18052 | 20,06158 | 20,75597 | 20,46533 | 21,10284 | 19,93460 | 20,78700 |
| | EH domain-containing protein 1 | Q8IY37 | EHD1 | 0,16860 | -0,30486 | 22,58777 | 22,72721 | 22,08835 | 22,55614 | 23,99886 | 21,76292 |
| | Putative helicase MOV-10 | Q9UKM9 | MOV10 | 0,16737 | -0,13671 | 21,34224 | 21,42834 | 21,83923 | 21,69511 | 21,19994 | 22,12487 |
| | Prohibitin | O95391 | PHB | 0,16707 | -0,59123 | 25,72567 | 25,43425 | 25,53249 | 24,73423 | 28,81696 | 24,91493 |
| | 2,3-cyclic-nucleotide 3-phosphodiesterase | P31040 | CNP | 0,16705 | -0,28114 | 22,79529 | 23,26456 | 23,46176 | 22,47936 | 24,55713 | 23,32855 |
| | KH domain-containing, RNA-binding, signal transduction-associated protein 1 | Q86VM9 | KHDRBS1 | 0,16558 | -0,67708 | 20,43288 | 22,11886 | 24,22494 | 21,99186 | 21,72217 | 25,09390 |
| | AP-3 complex subunit mu-1 | P11387 | AP3M1 | 0,16395 | -0,81448 | 19,68839 | 23,11997 | 24,20471 | 20,92441 | 23,16795 | 25,36416 |
| | Stress-70 protein, mitochondrial | O00567 | HSPA9 | 0,16359 | 0,11423 | 24,11458 | 24,85096 | 24,14009 | 24,41002 | 24,29551 | 24,05738 |
| | Putative ATP-dependent RNA helicase DHX30 | Q96FV9 | DHX30 | 0,16222 | -0,22351 | 23,51617 | 23,04459 | 24,50555 | 23,33743 | 24,22920 | 24,17020 |
| | Eukaryotic translation initiation factor 3 subunit D | Q9NR30 | EIF3D | 0,16106 | 0,44344 | 27,54782 | 24,68126 | 24,79328 | 24,53499 | 26,03578 | 25,12127 |
| | Probable dimethyladenosine transferase | P32969 | DIMT1 | 0,16058 | 0,83102 | 19,95617 | 19,60820 | 20,84261 | 15,50000 | 21,46399 | 20,94992 |
| | DNA-directed RNA polymerase II subunit RPB3 | P62888 | POLR2C | 0,16050 | 0,47651 | 20,81041 | 22,75165 | 21,38374 | 22,93882 | 19,66287 | 20,91459 |
| | Fanconi anemia group I protein | P78346 | FANCI | 0,16027 | 0,95672 | 19,44645 | 20,01127 | 19,61775 | 15,50000 | 23,02315 | 17,68216 |
| | F-actin-capping protein subunit alpha-2 | P50579 | CAPZA2 | 0,16011 | -0,24949 | 22,62589 | 22,45646 | 21,71708 | 23,30271 | 21,54948 | 22,69570 |
| | ATPase family AAA domain-containing protein 3B | Q8ND56 | ATAD3B | 0,15999 | 0,23158 | 24,52199 | 25,81558 | 25,68129 | 24,75364 | 25,81941 | 24,75108 |
| | ADP/ATP translocase 2;ADP/ATP translocase 2, N-terminally processed | Q9Y6J0 | SLC25A5 | 0,15921 | -0,20039 | 26,42622 | 26,59605 | 26,72273 | 26,02509 | 27,62624 | 26,69485 |
| | Pre-mRNA-processing factor 6 | P11274 | PRPF6 | 0,15919 | -0,38847 | 22,31415 | 22,02935 | 22,39906 | 20,87305 | 23,89874 | 23,13618 |
| | Actin-related protein 2/3 complex subunit 2 | P62906 | ARPC2 | 0,15918 | 0,08232 | 23,68581 | 23,40106 | 23,43593 | 23,29908 | 23,21120 | 23,76557 |
| | Myosin-9 | Q16630 | MYH9 | 0,15861 | 0,48384 | 28,15074 | 30,23055 | 29,60489 | 29,60804 | 26,93044 | 29,99618 |
| | ATP-dependent RNA helicase DHX36 | P67870 | DHX36 | 0,15849 | 0,09658 | 21,49863 | 21,56216 | 22,15515 | 21,77044 | 21,46324 | 21,69251 |
| | Heterogeneous nuclear ribonucleoprotein D-like | Q9Y5B9 | HNRNPDL | 0,15800 | 0,18796 | 23,49875 | 22,22529 | 23,37039 | 22,52162 | 22,83984 | 23,16910 |
| | Splicing factor 3B subunit 4 | Q9NYH9 | SF3B4 | 0,15782 | -0,37374 | 21,42783 | 21,85818 | 23,71869 | 21,62520 | 23,24447 | 23,25624 |
| | 60S ribosomal protein L7a | Q9UKV3 | RPL7A | 0,15647 | -0,33273 | 23,97150 | 24,91657 | 26,15041 | 24,38357 | 25,70886 | 25,94425 |

| Significant | Protein names | UniProtID | Gene names | -LOG(P-value) | Difference | CXCR4_1 | CXCR4_2 | CXCR4_3 | MOCK_1 | MOCK_2 | MOCK_3 |
|---|---|---|---|---|---|---|---|---|---|---|---|
| | Ras-related protein Rab-21 | Q13523 | RAB21 | 0,15497 | -0,58728 | 15,50000 | 19,93387 | 19,50275 | 18,78523 | 18,75682 | 19,15642 |
| | ATP-citrate synthase | P08708 | ACLY | 0,15411 | -0,52134 | 19,70261 | 15,50000 | 18,61087 | 18,44071 | 18,26467 | 18,67211 |
| | Insulin-like growth factor 2 mRNA-binding protein 2 | Q9NXF1 | IGF2BP2 | 0,15135 | -0,91092 | 15,50000 | 20,36378 | 22,35837 | 19,52043 | 19,23276 | 22,20171 |
| | | O43660 | | 0,15076 | 0,22869 | 22,89865 | 23,02017 | 22,52442 | 22,85177 | 21,53607 | 23,36933 |
| | 60S acidic ribosomal protein P0;60S acidic ribosomal protein P0-like | Q7Z4V5 | RPLP0;RPLP0P6 | 0,15006 | -0,33913 | 25,43406 | 25,30532 | 27,49503 | 25,51003 | 26,82869 | 26,91308 |
| | Very-long-chain 3-oxoacyl-CoA reductase | Q8IXW5 | HSD17B12 | 0,14862 | 0,25190 | 22,73324 | 23,21681 | 23,29278 | 22,65244 | 23,95633 | 21,87837 |
| | Digestive organ expansion factor homolog | P43250 | DIEXF | 0,14807 | 0,86799 | 21,21810 | 20,58912 | 20,27378 | 15,50000 | 21,85304 | 22,12398 |
| | UPF0609 protein C4orf27;Putative UPF0609 protein C4orf27-like | P08621 | C4orf27 | 0,14780 | 0,76109 | 20,39257 | 20,61997 | 18,62296 | 20,83414 | 21,01808 | 15,50000 |
| | Mitochondrial import inner membrane translocase subunit TIM44 | Q92541 | TIMM44 | 0,14760 | -0,29193 | 21,31152 | 21,80664 | 21,28935 | 21,33074 | 23,15966 | 20,79289 |
| | E3 ubiquitin-protein ligase TRIM21 | P61106 | TRIM21 | 0,14662 | 0,35537 | 25,21100 | 25,04252 | 23,17207 | 25,10909 | 22,97540 | 24,27498 |
| | Eukaryotic translation initiation factor 3 subunit H | O76094 | EIF3H | 0,14470 | 0,64610 | 27,34274 | 22,76586 | 23,04926 | 22,80886 | 25,21534 | 23,19535 |
| | Dolichol-phosphate mannosyltransferase subunit 1 | O75915 | DPM1 | 0,14459 | 0,21116 | 21,39179 | 21,62761 | 21,51036 | 20,40416 | 22,26382 | 21,22829 |
| | Annexin A7 | Q7Z7C8 | ANXA7 | 0,14204 | 0,36080 | 21,12364 | 19,68425 | 18,42509 | 20,31252 | 18,48511 | 19,35296 |
| | Polyadenylate-binding protein 4 | O75027 | PABPC4 | 0,14161 | -0,58475 | 21,24714 | 23,36333 | 24,61880 | 21,31904 | 24,76562 | 24,89883 |
| | Poly [ADP-ribose] polymerase 1 | P23458 | PARP1 | 0,14142 | -0,47055 | 27,65315 | 28,13026 | 30,56420 | 28,19761 | 28,64715 | 30,91449 |
| | Peroxiredoxin-1 | Q13283 | PRDX1 | 0,14117 | -0,17636 | 24,39696 | 25,70744 | 25,21560 | 25,72954 | 24,82610 | 25,29344 |
| | Actin, cytoplasmic 2;Actin, cytoplasmic 2, N-terminally processed | Q9Y608 | ACTG1 | 0,14089 | -0,17200 | 30,94914 | 31,04362 | 30,46519 | 31,05187 | 30,24400 | 31,67808 |
| | SWI/SNF complex subunit SMARCC2 | Q4KMP7 | SMARCC2 | 0,13926 | -0,25318 | 21,90084 | 20,90102 | 20,46563 | 21,90500 | 21,82108 | 20,30095 |
| | Annexin A11 | Q8NBQ5 | ANXA11 | 0,13741 | -0,12515 | 19,47007 | 18,95231 | 19,74588 | 19,75512 | 19,02842 | 19,76017 |
| | E3 ubiquitin-protein ligase listerin | P46781 | LTN1 | 0,13336 | -0,90455 | 18,42181 | 18,12767 | 18,27030 | 15,50000 | 23,94172 | 18,09171 |
| | Lysine-specific demethylase 5A | Q96HR3 | KDM5A | 0,13066 | -0,17821 | 20,91298 | 21,82205 | 20,65971 | 22,01369 | 20,86423 | 21,05146 |
| | Unconventional myosin-Ib | Q9ULX6 | MYO1B | 0,13015 | -0,03848 | 22,83447 | 22,96819 | 22,76460 | 22,87807 | 23,05887 | 22,74577 |
| | 28S ribosomal protein S17, mitochondrial | Q96BK5 | hCG_1984214;MRPS17 | 0,12893 | -0,25726 | 21,60214 | 21,61054 | 20,95709 | 21,53863 | 22,90987 | 20,49306 |
| | Coatomer subunit alpha;Xenin;Proxenin | Q8N1G4 | COPA | 0,12882 | 0,20058 | 26,16620 | 25,66117 | 26,36966 | 25,08663 | 26,88075 | 25,62792 |
| | Solute carrier family 35 member E1 | Q9UQB8 | SLC35E1 | 0,12857 | -0,08636 | 20,27150 | 20,43675 | 19,86101 | 20,62819 | 20,13846 | 20,06171 |
| | Serine/arginine-rich splicing factor 6 | P39019 | SRSF6 | 0,12678 | 0,25945 | 25,37438 | 23,38192 | 25,44550 | 23,92116 | 24,45858 | 25,04373 |
| | F-actin-capping protein subunit beta | Q9H204 | CAPZB | 0,12661 | 0,18047 | 23,88344 | 23,86561 | 22,90301 | 23,76182 | 22,55084 | 23,79798 |
| | Polyadenylate-binding protein 2 | P04049 | PABPN1 | 0,12603 | 0,98590 | 21,27714 | 21,25194 | 23,36092 | 15,50000 | 22,90128 | 24,53101 |
| | Ankycorbin | P37802 | RAI14 | 0,12592 | 0,19764 | 25,10941 | 25,10378 | 24,58383 | 25,48173 | 23,66709 | 25,05528 |
| | Transformer-2 protein homolog beta | Q9Y6V7 | TRA2B | 0,12569 | -0,22224 | 21,01468 | 21,16582 | 23,02748 | 21,98155 | 21,92198 | 21,97116 |
| | Methionine--tRNA ligase, cytoplasmic | O14646 | MARS | 0,12503 | -0,45072 | 22,72451 | 24,22729 | 24,16108 | 22,99752 | 26,60371 | 22,86380 |
| | Prohibitin-2 | Q04837 | PHB2 | 0,12340 | -0,48973 | 25,72182 | 24,60275 | 25,74368 | 23,70978 | 28,48424 | 25,34344 |
| | Peroxiredoxin-4 | Q9H7Z3 | PRDX4 | 0,12272 | -0,20198 | 19,98690 | 18,99148 | 19,24869 | 20,26762 | 18,57946 | 19,98593 |
| | Serine/threonine-protein phosphatase PGAM5, mitochondrial | Q8TAF3 | PGAM5 | 0,12267 | 0,11330 | 23,50567 | 23,62377 | 23,34204 | 22,73713 | 23,57750 | 23,81695 |
| | Dehydrogenase/reductase SDR family member 7B | Q9H2H8 | DHRS7B | 0,12263 | 0,18154 | 20,64588 | 20,30563 | 19,32562 | 20,14271 | 20,40104 | 19,18877 |
| | Vesicle-trafficking protein SEC22b | P06753 | SEC22B | 0,12133 | -0,64011 | 15,50000 | 21,50964 | 20,93984 | 20,35316 | 19,76662 | 19,75004 |
| | AP-3 complex subunit delta-1 | Q9H9A6 | AP3D1 | 0,12104 | -0,26617 | 26,23422 | 28,00013 | 27,45301 | 28,21688 | 26,28371 | 27,98528 |
| | Ig kappa chain C region | Q15311 | IGKC | 0,12045 | 0,33779 | 19,58937 | 19,15026 | 17,13911 | 19,38499 | 17,01193 | 18,46846 |
| | Replication factor C subunit 5 | Q9UQ35 | RFC5 | 0,11771 | -0,18077 | 22,01431 | 21,86136 | 22,28842 | 21,19123 | 23,02724 | 22,48792 |
| | Serrate RNA effector molecule homolog | Q00059 | SRRT | 0,11634 | 0,13398 | 21,90268 | 21,51118 | 22,61349 | 21,59619 | 21,62055 | 22,40868 |
| | Syntaxin-binding protein 3 | A1A4S6 | STXBP3 | 0,11634 | 0,13267 | 20,82920 | 21,03946 | 20,74706 | 21,13746 | 21,15171 | 19,92852 |
| | Serine/arginine-rich splicing factor 9 | P62851 | SRSF9 | 0,11568 | -0,21733 | 21,65009 | 21,78153 | 22,97322 | 21,38790 | 22,42275 | 23,24616 |
| | Transducin beta-like protein 2 | O15381 | TBL2 | 0,11421 | -0,24519 | 23,78576 | 23,76527 | 24,02775 | 22,56695 | 25,03666 | 24,71072 |
| | Non-POU domain-containing octamer-binding protein | P78345 | NONO | 0,11412 | 0,25925 | 23,60298 | 24,39669 | 25,04773 | 24,24111 | 22,78995 | 25,23860 |
| | Unconventional myosin-XVIIIa | A0JLT2 | MYO18A | 0,11404 | -0,50984 | 20,74641 | 21,07156 | 24,35494 | 21,89034 | 21,01658 | 24,79551 |
| | Rab-like protein 6 | Q9UHR4 | RABL6 | 0,11231 | -0,25010 | 23,64559 | 25,05155 | 24,40750 | 25,46585 | 23,23523 | 25,15387 |
| | Lamin-B1 | Q9Y388 | LMNB1 | 0,11157 | -0,16533 | 22,22747 | 23,51569 | 23,36746 | 23,67713 | 22,51934 | 23,41015 |
| | Cytoplasmic dynein 1 light intermediate chain 1 | P35659 | DYNC1LI1 | 0,11054 | 0,43934 | 21,45934 | 21,36736 | 20,27787 | 20,38669 | 23,09637 | 18,30350 |
| | Tropomyosin alpha-4 chain | Q9NPG3 | TPM4 | 0,10638 | 0,45648 | 21,73211 | 23,85823 | 24,22847 | 22,76225 | 20,52612 | 25,16100 |
| | DNA-directed RNA polymerase II subunit RPB2 | O60885 | POLR2B | 0,10576 | 0,29354 | 22,09742 | 23,62444 | 22,86995 | 23,67799 | 20,79130 | 23,24191 |
| | Interleukin enhancer-binding factor 2 | P32780 | ILF2 | 0,10419 | -0,35615 | 23,35138 | 23,35367 | 25,84410 | 22,98167 | 24,50743 | 26,12851 |
| | Peptidyl-prolyl cis-trans isomerase A;Peptidyl-prolyl cis-trans isomerase A, N-terminally processed | Q92572 | PPIA | 0,10348 | -0,63288 | 22,17435 | 15,50000 | 20,98607 | 21,64689 | 18,94724 | 19,96494 |
| | Eukaryotic translation initiation factor 3 subunit L | Q13459 | EIF3L | 0,10117 | 0,32691 | 28,02770 | 24,66958 | 25,07887 | 24,79853 | 26,45197 | 25,54492 |
| | Eukaryotic translation initiation factor 3 subunit G | Q8WXA9 | EIF3G | 0,10038 | 0,45907 | 27,72032 | 23,21719 | 23,10495 | 23,26714 | 25,38555 | 24,01257 |
| | Eukaryotic translation initiation factor 3 subunit F | Q03111 | EIF3F | 0,09961 | 0,47844 | 28,23868 | 23,46973 | 24,12084 | 23,22083 | 26,18291 | 24,99019 |
| | ATPase family AAA domain-containing protein 3A | Q8N442 | ATAD3A | 0,09941 | 0,16074 | 23,51063 | 24,67573 | 24,71560 | 23,48468 | 24,93508 | 23,99998 |
| | Splicing factor 3A subunit 1 | P62805 | SF3A1 | 0,09866 | -0,15349 | 21,22923 | 21,06586 | 21,97656 | 21,08479 | 21,10578 | 22,54154 |
| | 40S ribosomal protein S12 | P52434 | RPS12 | 0,09813 | 0,20275 | 24,58583 | 23,34354 | 25,36840 | 24,03867 | 23,56755 | 25,08331 |
| | Sec1 family domain-containing protein 1 | Q9NVP1 | SCFD1 | 0,09473 | -0,47520 | 15,50000 | 20,45573 | 21,04674 | 19,13779 | 20,16779 | 19,12250 |
| | Heterogeneous nuclear ribonucleoproteins A2/B1 | Q9UBB9 | HNRNPA2B1 | 0,09382 | -0,37592 | 23,49424 | 22,83115 | 25,72200 | 23,92796 | 22,70375 | 26,54345 |
| | Structural maintenance of chromosomes protein 3 | Q9UGN5 | SMC3 | 0,09178 | -0,15234 | 24,10093 | 24,48677 | 23,62734 | 24,26091 | 25,13489 | 23,27625 |
| | Proliferation-associated protein 2G4 | P30048 | PA2G4 | 0,09130 | -0,22827 | 21,82003 | 20,85285 | 22,41465 | 21,05325 | 21,26837 | 23,45073 |

| Significant | Protein names | UniProtID | Gene names | -LOG(P-value) | Difference | CXCR4_1 | CXCR4_2 | CXCR4_3 | MOCK_1 | MOCK_2 | MOCK_3 |
|---|---|---|---|---|---|---|---|---|---|---|---|
| | BAG family molecular chaperone regulator 4 | Q8WXI9 | BAG4 | 0,09122 | 0,24800 | 20,12578 | 21,04313 | 19,00407 | 21,34251 | 18,94404 | 19,14243 |
| | Structural maintenance of chromosomes protein 1A | Q8WUA2 | SMC1A | 0,08972 | -0,15263 | 24,75471 | 24,96080 | 24,10061 | 24,48898 | 25,81081 | 23,97421 |
| | Dolichyl-diphosphooligosaccharide--protein glycosyltransferase subunit 1 | O00268 | RPN1 | 0,08802 | -0,25790 | 20,90793 | 23,16823 | 23,39231 | 21,48331 | 23,78506 | 22,97379 |
| | Monocarboxylate transporter 1 | P40222 | SLC16A1 | 0,08502 | 0,08927 | 22,69213 | 22,32460 | 22,40717 | 21,96183 | 23,09081 | 22,10345 |
| | Eukaryotic translation initiation factor 3 subunit K | Q9H939 | EIF3K | 0,08359 | -0,32302 | 25,97919 | 21,99002 | 23,19025 | 23,05654 | 25,36940 | 23,70261 |
| | DNA-dependent protein kinase catalytic subunit | P59780 | PRKDC | 0,08325 | -0,26122 | 28,60791 | 28,28012 | 27,99717 | 27,73622 | 30,72728 | 27,20536 |
| | Serine/threonine-protein phosphatase PP1-gamma catalytic subunit | Q96GA3 | PPP1CC | 0,08153 | -0,11845 | 21,10796 | 21,23538 | 22,66477 | 21,55234 | 21,94946 | 21,86166 |
| | Thyroid receptor-interacting protein 6 | P82673 | TRIP6 | 0,08100 | -0,20193 | 21,03617 | 22,30502 | 19,92220 | 22,38445 | 20,66969 | 20,81503 |
| | Myosin phosphatase Rho-interacting protein | Q7L804 | MPRIP | 0,08081 | 0,12988 | 25,17927 | 25,38765 | 24,82323 | 25,82944 | 23,97806 | 25,19302 |
| | Probable ATP-dependent RNA helicase DDX27 | P78362 | DDX27 | 0,08036 | 0,60829 | 20,53716 | 20,49697 | 22,70489 | 15,50000 | 23,35380 | 23,06034 |
| | Nucleolin | Q9HCM4 | NCL | 0,07833 | -0,71140 | 19,51266 | 22,02317 | 27,61821 | 19,95617 | 24,04093 | 27,29114 |
| | 40S ribosomal protein S13 | Q9NSY1 | RPS13 | 0,07807 | -0,18791 | 24,07953 | 23,71910 | 25,74119 | 23,73934 | 24,63439 | 25,72983 |
| | Far upstream element-binding protein 2 | Q9NUD5 | KHSRP | 0,07788 | 0,23892 | 22,55696 | 25,38242 | 24,77666 | 24,43301 | 22,71125 | 24,85500 |
| | Glioma tumor suppressor candidate region gene 1 protein | P19525 | GLTSCR1 | 0,07610 | -0,34586 | 18,98449 | 22,33824 | 20,53089 | 22,95897 | 18,60241 | 21,32981 |
| | mRNA export factor | Q8TBB5 | RAE1 | 0,07552 | 0,40530 | 19,76259 | 19,85930 | 19,26947 | 15,50000 | 21,49629 | 20,67917 |
| | Leucine--tRNA ligase, cytoplasmic | Q09161 | LARS | 0,07414 | 0,18154 | 24,58549 | 24,33204 | 24,29908 | 23,60445 | 25,91367 | 23,15386 |
| | Spectrin beta chain, non-erythrocytic 1 | Q9Y3U8 | SPTBN1 | 0,07225 | 0,14647 | 22,98829 | 21,32680 | 21,33974 | 22,52943 | 20,97953 | 21,70645 |
| | Eukaryotic translation initiation factor 3 subunit E | O15258 | EIF3E | 0,07223 | 0,32664 | 28,86800 | 24,50500 | 24,85705 | 24,46176 | 27,03946 | 25,74891 |
| | PRKC apoptosis WT1 regulator protein | P55199 | PAWR | 0,07099 | -0,19061 | 23,42432 | 25,24438 | 23,93688 | 25,25090 | 22,86011 | 25,06641 |
| | Condensin-2 complex subunit G2 | Q969X6 | NCAPG2 | 0,06968 | -0,35529 | 17,92457 | 17,96556 | 18,52715 | 15,50000 | 21,63423 | 18,34893 |
| | 40S ribosomal protein S4, X isoform;40S ribosomal protein S4, Y isoform 2 | Q9NRL2 | RPS4X;RPS4Y2 | 0,06921 | -0,12852 | 25,36546 | 25,16127 | 26,72573 | 25,52800 | 25,38574 | 26,72429 |
| | C2 domain-containing protein 5 | Q9H813 | C2CD5 | 0,06768 | -0,10649 | 20,46553 | 21,28455 | 20,86642 | 21,45528 | 19,98760 | 21,49311 |
| | Adenosylhomocysteinase | Q8N0V3 | AHCY | 0,06597 | -0,07264 | 24,11926 | 23,56837 | 22,95824 | 23,85158 | 23,25181 | 23,76040 |
| | Heterogeneous nuclear ribonucleoprotein D0 | Q6PCB5 | HNRNPD | 0,06497 | -0,25371 | 23,37941 | 22,32948 | 25,65297 | 23,23649 | 22,96890 | 25,91759 |
| | Elongation factor 1-gamma | Q6UN15 | EEF1G | 0,06457 | -0,12093 | 25,97631 | 25,63735 | 25,73286 | 25,30741 | 27,18990 | 25,21201 |
| | Transformer-2 protein homolog alpha | Q13823 | TRA2A | 0,06447 | 0,11035 | 20,80633 | 20,61306 | 22,45824 | 21,04267 | 21,10578 | 21,39812 |
| | Elongation factor 1-delta | Q9NYF8 | EEF1D | 0,06377 | -0,14780 | 23,45588 | 23,51953 | 23,80371 | 23,09207 | 25,32971 | 22,80073 |
| | Pre-mRNA-splicing regulator WTAP | Q8IYB3 | WTAP | 0,06301 | 0,36134 | 20,05033 | 18,05928 | 20,35412 | 15,50000 | 21,64504 | 20,23468 |
| | Heterogeneous nuclear ribonucleoprotein U | P0DN76 | HNRNPU | 0,06282 | -0,16396 | 23,99516 | 23,48210 | 25,94849 | 23,83807 | 24,49875 | 25,58081 |
| | Structural maintenance of chromosomes protein 4 | P82912 | SMC4 | 0,06202 | -0,23135 | 22,46164 | 21,73603 | 21,33669 | 21,99838 | 24,28369 | 19,94635 |
| | Structural maintenance of chromosomes protein 2 | P27635 | SMC2 | 0,06176 | 0,16786 | 23,25048 | 23,08568 | 22,47073 | 22,63578 | 24,41138 | 21,25615 |
| | DnaJ homolog subfamily A member 1 | O60506 | DNAJA1 | 0,06118 | -0,13002 | 24,91036 | 26,15000 | 25,66330 | 24,75440 | 26,93067 | 25,42864 |
| | Guanine nucleotide-binding protein subunit beta-2-like 1;Guanine nucleotide-binding protein subunit beta-2-like 1, N-terminally processed | O95816 | GNB2L1 | 0,06104 | -0,08064 | 24,63926 | 24,47729 | 24,44929 | 25,18877 | 24,91109 | 23,70788 |
| | Heterogeneous nuclear ribonucleoprotein H3 | Q9P2D1 | HNRNPH3 | 0,06047 | -0,07802 | 23,64141 | 22,76563 | 22,77676 | 23,43084 | 22,46044 | 23,52659 |
| | DNA-directed RNA polymerase II subunit RPB1 | P38432 | POLR2A | 0,06003 | -0,23595 | 20,95333 | 24,10661 | 23,10759 | 24,30083 | 21,01502 | 23,55953 |
| | THO complex subunit 5 homolog | Q9NVU7 | THOC5 | 0,05938 | 0,63154 | 21,22087 | 23,30786 | 24,88320 | 15,50000 | 26,73299 | 25,28433 |
| | Histone-lysine N-methyltransferase SETD1A | P49757 | SETD1A | 0,05804 | -0,09588 | 21,94460 | 21,60399 | 22,06464 | 22,03184 | 20,97554 | 22,89349 |
| | Eukaryotic translation elongation factor 1 epsilon-1 | Q9C086 | EEF1E1;EEF1E1-BLOC1S5 | 0,05746 | 0,24551 | 23,31935 | 23,73066 | 22,14049 | 21,90172 | 25,56436 | 20,98787 |
| | Inosine-5-monophosphate dehydrogenase 2 | P82914 | IMPDH2 | 0,05712 | 0,04575 | 22,70345 | 22,28704 | 22,88051 | 22,36399 | 23,00554 | 22,36421 |
| | AP-2 complex subunit alpha-2 | Q12906 | AP2A2 | 0,05692 | -0,54102 | 18,82311 | 20,48670 | 23,48886 | 15,50000 | 24,57819 | 24,34354 |
| | Serine/threonine-protein phosphatase 2A 55 kDa regulatory subunit B alpha isoform | Q15424 | PPP2R2A | 0,05676 | 0,08440 | 22,86699 | 23,50276 | 22,62000 | 23,72515 | 22,80385 | 22,20756 |
| | Endoplasmic reticulum resident protein 44 | Q14241 | ERP44 | 0,05671 | 0,07929 | 20,54265 | 20,24039 | 19,71102 | 20,72025 | 20,23760 | 19,29834 |
| | Histone deacetylase 1 | Q9H9F9 | HDAC1 | 0,05670 | -0,08683 | 20,96409 | 19,75398 | 20,27708 | 20,11794 | 21,20425 | 19,93344 |
| | ATP-dependent RNA helicase DDX3X;ATP-dependent RNA helicase DDX3Y | Q96HR8 | DDX3X;DDX3Y | 0,05312 | -0,11236 | 25,20769 | 25,57491 | 26,59761 | 24,76744 | 26,15731 | 26,79253 |
| | ATP-dependent RNA helicase DDX24 | P62917 | DDX24 | 0,05281 | 0,08281 | 21,85190 | 22,12490 | 21,65923 | 22,81650 | 21,09016 | 21,48094 |
| | 40S ribosomal protein S5;40S ribosomal protein S5, N-terminally processed | Q5VWQ0 | RPS5 | 0,05272 | -0,12359 | 25,05912 | 24,87282 | 26,60724 | 24,55479 | 25,76681 | 26,58835 |
| | Splicing factor, proline- and glutamine-rich | Q14692 | SFPQ | 0,05204 | -0,21228 | 23,26385 | 24,15661 | 26,13866 | 24,12684 | 23,17234 | 26,89678 |
| | Gelsolin | Q9NW64 | GSN | 0,05091 | -0,10856 | 23,38245 | 24,59406 | 23,63998 | 24,58829 | 22,71429 | 24,63959 |
| | Slit homolog 2 protein;Slit homolog 2 protein N-product;Slit homolog 2 protein C-product | P62750 | SLIT2 | 0,04989 | -0,09434 | 20,97057 | 20,81691 | 22,21244 | 20,99065 | 20,91452 | 22,37777 |
| | 5-3 exoribonuclease 2 | Q8IXM2 | XRN2 | 0,04952 | -0,22670 | 22,16417 | 20,55759 | 23,83721 | 20,08499 | 22,77175 | 24,38232 |
| | Cleavage and polyadenylation specificity factor subunit 1 | O75400 | CPSF1 | 0,04927 | -0,07895 | 22,46191 | 21,08472 | 22,74587 | 21,78457 | 22,32934 | 22,41545 |
| | Coronin-1C | Q86UY6 | CORO1C | 0,04811 | -0,18397 | 21,38174 | 22,81208 | 23,95633 | 22,66125 | 21,16429 | 24,87652 |
| | Heterogeneous nuclear ribonucleoprotein M | P63162 | HNRNPM | 0,04728 | 0,14787 | 24,12116 | 24,82046 | 26,87103 | 23,96825 | 25,06989 | 26,33091 |
| | ATP-binding cassette sub-family E member 1 | Q9P0U4 | ABCE1 | 0,04621 | -0,02193 | 21,67793 | 21,73057 | 21,47749 | 21,85190 | 21,37433 | 21,72555 |
| | ATP-dependent RNA helicase DDX1 | Q15287 | DDX1 | 0,04415 | -0,10788 | 23,57138 | 24,39021 | 25,40510 | 23,31866 | 24,90104 | 25,47063 |
| | Sorting and assembly machinery component 50 homolog | P82930 | SAMM50 | 0,04278 | 0,11501 | 22,08404 | 22,34196 | 22,31874 | 20,76782 | 23,86561 | 21,76628 |
| | Dermcidin;Survival-promoting peptide;DCD-1 | P62318 | DCD | 0,04157 | -0,06903 | 22,48729 | 21,30463 | 20,90595 | 22,24812 | 21,35601 | 21,30084 |
| | LIM domain only protein 7 | Q6PJT7 | LMO7 | 0,04092 | 0,07625 | 23,03226 | 22,04183 | 22,23503 | 23,00347 | 21,25078 | 22,82611 |
| | Protein phosphatase 1G | P20339 | PPM1G | 0,03942 | -0,27026 | 15,50000 | 22,23739 | 22,20066 | 19,13310 | 21,34868 | 20,26705 |
| | Multiple myeloma tumor-associated protein 2 | Q9NPI1 | MMTAG2 | 0,03854 | 0,26966 | 18,37331 | 20,88889 | 21,15128 | 15,50000 | 21,53820 | 22,56630 |
| | Caprin-1 | Q02040 | CAPRIN1 | 0,03837 | -0,08660 | 21,56565 | 21,73529 | 23,45212 | 21,43002 | 22,55918 | 23,02367 |

| Significant | Protein names | UniProtID | Gene names | -LOG(P-value) | Difference | CXCR4_1 | CXCR4_2 | CXCR4_3 | MOCK_1 | MOCK_2 | MOCK_3 |
|---|---|---|---|---|---|---|---|---|---|---|---|
| | Host cell factor 1;HCF N-terminal chain 1;HCF N-terminal chain 2;HCF N-terminal chain 3;HCF N-terminal chain 4;HCF N-terminal chain 5;HCF N-terminal chain 6;HCF C-terminal chain 1;HCF C-terminal chain 2;HCF C-terminal chain 3;HCF C-terminal chain 4;HCF C-terminal chain 5;HCF C-terminal chain 6 | Q9NVC6 | HCFC1 | 0,03812 | 0,05885 | 21,92147 | 21,12155 | 21,64037 | 21,36341 | 20,76919 | 22,37425 |
| | Twinfilin-1 | O00541 | TWF1 | 0,03744 | -0,11205 | 20,50125 | 19,39672 | 19,95134 | 20,65796 | 18,17694 | 21,35057 |
| | Alpha-actinin-4 | Q9BXJ9 | ACTN4 | 0,03740 | 0,11799 | 22,63611 | 23,10213 | 24,27341 | 22,43642 | 22,10665 | 25,11462 |
| | DnaJ homolog subfamily A member 2 | Q9UNX4 | DNAJA2 | 0,03660 | -0,11351 | 23,76000 | 24,66427 | 25,39257 | 23,55157 | 26,57586 | 24,02994 |
| | THO complex subunit 3 | Q9Y5B6 | THOC3 | 0,03472 | 0,30424 | 21,44304 | 19,55388 | 22,05815 | 15,50000 | 25,38400 | 21,25834 |
| | Splicing factor U2AF 65 kDa subunit | Q9NY12 | U2AF2 | 0,03459 | -0,07613 | 22,66897 | 22,86610 | 24,03431 | 22,47276 | 22,85928 | 24,46576 |
| | Unconventional myosin-Id | O76021 | MYO1D | 0,03299 | 0,04253 | 24,65949 | 24,10197 | 23,38034 | 24,44708 | 23,68314 | 23,88400 |
| | Nucleolar complex protein 4 homolog | P46779 | NOC4L | 0,03297 | -0,26102 | 19,39253 | 19,93171 | 21,55913 | 15,50000 | 24,07447 | 22,09197 |
| | Homeobox protein Hox-A5 | P53680 | HOXA5 | 0,03261 | 0,07275 | 18,05907 | 18,72770 | 17,00273 | 18,89445 | 16,96826 | 17,70853 |
| | DnaJ homolog subfamily C member 7 | Q5VT06 | DNAJC7 | 0,03218 | -0,06412 | 23,29572 | 23,68143 | 22,92606 | 23,34191 | 24,47772 | 22,27594 |
| | WD repeat-containing protein 61;WD repeat-containing protein 61, N-terminally processed | Q8NHQ9 | WDR61 | 0,03185 | 0,06624 | 23,24080 | 21,47467 | 23,09215 | 21,72771 | 22,78003 | 23,10116 |
| | DNA replication licensing factor MCM3 | P0DMU9 | MCM3 | 0,03156 | 0,09752 | 20,91546 | 23,12180 | 23,74745 | 21,31954 | 23,12250 | 23,05011 |
| | GTP-binding protein SAR1b | Q14137 | SAR1B | 0,03045 | -0,06066 | 21,42388 | 23,14088 | 22,26313 | 22,71946 | 21,43609 | 22,85432 |
| | Serine/threonine-protein kinase Chk1 | Q15397 | CHEK1 | 0,03027 | 0,05911 | 21,18876 | 20,96416 | 20,22541 | 20,71154 | 21,76571 | 19,72374 |
| | U3 small nucleolar ribonucleoprotein protein MPP10 | Q99459 | MPHOSPH10 | 0,02950 | 0,19370 | 20,41349 | 19,58371 | 20,12792 | 15,50000 | 22,55272 | 21,49130 |
| | Tubulin alpha-1A chain;Tubulin alpha-3C/D chain;Tubulin alpha-3E chain | P84098 | TUBA1A;TUBA3C;TUBA3E | 0,02922 | -0,22481 | 15,50000 | 22,05818 | 23,72005 | 19,50880 | 21,68432 | 20,75955 |
| | 40S ribosomal protein S18 | Q12802 | RPS18 | 0,02825 | 0,06186 | 24,26420 | 24,07765 | 25,79181 | 24,00299 | 24,32031 | 25,62481 |
| | Midasin | Q8TAE8 | MDN1 | 0,02821 | 0,15025 | 22,02871 | 22,11296 | 22,04943 | 19,98064 | 25,48738 | 20,27230 |
| | Phosphate carrier protein, mitochondrial | Q9GZR2 | SLC25A3 | 0,02816 | -0,06987 | 25,32089 | 25,19054 | 25,77188 | 24,13751 | 26,95405 | 25,40136 |
| | La-related protein 1 | Q96SB4 | LARP1 | 0,02717 | -0,03657 | 22,12783 | 21,85034 | 23,25609 | 22,25790 | 22,36872 | 22,71735 |
| | Galectin-3-binding protein | P22087 | LGALS3BP | 0,02634 | -0,02519 | 26,05596 | 26,48830 | 25,60515 | 26,42707 | 25,75634 | 26,04158 |
| | 40S ribosomal protein S14 | P13010 | RPS14 | 0,02627 | -0,03015 | 25,68445 | 26,35795 | 25,54848 | 26,45425 | 25,75679 | 25,47029 |
| | Uveal autoantigen with coiled-coil domains and ankyrin repeats | Q13112 | UACA | 0,02473 | 0,03816 | 20,01727 | 20,11464 | 20,81488 | 20,14271 | 19,57204 | 21,11756 |
| | Aminoacyl tRNA synthase complex-interacting multifunctional protein 2 | Q15046 | AIMP2 | 0,02464 | -0,09380 | 22,62013 | 23,56860 | 21,94056 | 22,31061 | 25,05453 | 21,04554 |
| | DNA replication licensing factor MCM6 | Q9BZE4 | MCM6 | 0,02447 | -0,17637 | 15,50000 | 22,70024 | 22,59780 | 19,90563 | 20,26568 | 21,15586 |
| | Protein flightless-1 homolog | P61313 | FLII | 0,02439 | 0,07238 | 21,24377 | 21,89826 | 23,08737 | 21,76773 | 20,69294 | 23,55157 |
| | Coatomer subunit epsilon | P82675 | COPE | 0,02372 | -0,03637 | 23,06960 | 23,34787 | 23,22186 | 22,88881 | 24,24909 | 22,61052 |
| | Isoleucine--tRNA ligase, cytoplasmic | Q86UE4 | IARS | 0,02240 | -0,07995 | 25,11672 | 25,02466 | 24,79427 | 24,32464 | 27,38212 | 23,46874 |
| | Valine--tRNA ligase | Q6ZU65 | VARS | 0,02203 | -0,05764 | 23,10487 | 22,66899 | 23,10434 | 22,21086 | 24,73856 | 22,10172 |
| | Nucleosome-remodeling factor subunit BPTF | O43809 | BPTF | 0,02162 | -0,04869 | 23,42355 | 24,06011 | 23,58245 | 24,86538 | 22,38013 | 23,96667 |
| | Bifunctional glutamate/proline--tRNA ligase;Glutamate--tRNA ligase;Proline--tRNA ligase | P83731 | EPRS | 0,02069 | -0,07308 | 25,81519 | 25,52441 | 25,18067 | 24,80430 | 27,86656 | 24,06866 |
| | TATA-binding protein-associated factor 2N | Q9Y672 | TAF15 | 0,02012 | -0,05293 | 22,98423 | 23,07460 | 21,60612 | 23,81597 | 21,27560 | 22,73217 |
| | Unconventional myosin-Ie | Q9Y450 | MYO1E | 0,01932 | -0,02476 | 22,61571 | 21,96423 | 21,88933 | 21,66418 | 22,87065 | 22,00874 |
| | 40S ribosomal protein S27;40S ribosomal protein S27-like | P82663 | RPS27;RPS27L | 0,01651 | -0,04646 | 21,51186 | 20,89844 | 21,45287 | 19,69362 | 22,84826 | 21,46069 |
| | Replication protein A 70 kDa DNA-binding subunit;Replication protein A 70 kDa DNA-binding subunit, N-terminally processed | Q07020 | RPA1 | 0,01488 | -0,06509 | 19,22743 | 21,39519 | 24,00247 | 21,12060 | 21,21780 | 22,48198 |
| | RNA-binding protein EWS | P82664 | EWSR1 | 0,01452 | -0,05339 | 23,43377 | 25,75766 | 26,21015 | 24,41448 | 24,24038 | 26,90690 |
| | Spliceosome RNA helicase DDX39B | Q9Y3D9 | DDX39B | 0,01428 | -0,02528 | 22,81376 | 24,27966 | 24,10037 | 23,09079 | 23,84795 | 24,33088 |
| | Myosin regulatory light chain 12A;Myosin regulatory light chain 12B;Myosin regulatory light polypeptide 9 | Q9UJV9 | MYL12A;MYL12B;MYL9 | 0,01382 | -0,03319 | 25,44802 | 26,86667 | 26,83665 | 26,92434 | 25,14188 | 27,18471 |
| | F-actin-capping protein subunit alpha-1 | O75934 | CAPZA1 | 0,01217 | 0,02146 | 24,98898 | 24,99365 | 24,29453 | 25,47469 | 23,70028 | 25,03779 |
| | Neurabin-2 | Q96A72 | PPP1R9B | 0,01212 | -0,02042 | 21,72134 | 20,90242 | 21,45282 | 22,00631 | 20,38774 | 21,74378 |
| | Myosin-10 | O43513 | MYH10 | 0,01132 | 0,01792 | 30,10625 | 30,62680 | 30,16926 | 30,91249 | 29,30437 | 30,63166 |
| | Protein TFG | O75044 | TFG | 0,01126 | 0,03641 | 30,18542 | 29,82701 | 27,51578 | 30,21477 | 29,26870 | 27,93551 |
| | Parafibromin | P62266 | CDC73 | 0,00690 | 0,01546 | 23,95669 | 21,47378 | 22,51925 | 22,52296 | 22,93287 | 22,44751 |
| | Protein phosphatase 1 regulatory subunit 12A | Q9Y2Q9 | PPP1R12A | 0,00580 | 0,01562 | 20,24214 | 21,93366 | 21,97778 | 22,06095 | 20,02337 | 22,02239 |
| | Elongation factor 1-beta | Q96G25 | EEF1B2 | 0,00547 | 0,01423 | 22,36573 | 22,54597 | 23,54144 | 21,66132 | 24,26964 | 22,47951 |
| | ATP-dependent RNA helicase A | P61353 | DHX9 | 0,00265 | 0,00892 | 24,79823 | 23,52169 | 26,36773 | 23,69891 | 24,74263 | 26,21935 |
| | Aminoacyl tRNA synthase complex-interacting multifunctional protein 1;Endothelial monocyte-activating polypeptide 2 | O14880 | AIMP1 | 0,00258 | 0,01133 | 23,40250 | 23,71177 | 22,88103 | 21,81382 | 26,15234 | 21,99514 |
| | DnaJ homolog subfamily B member 11 | Q8IY81 | DNAJB11 | 0,00238 | -0,00497 | 21,41338 | 22,88855 | 21,38158 | 21,73451 | 22,77696 | 21,18695 |
| | B-cell receptor-associated protein 31 | O43818 | BCAP31 | 0,00111 | -0,00314 | 21,46802 | 22,62665 | 21,81538 | 21,30986 | 23,67444 | 20,93517 |
| | Very-long-chain (3R)-3-hydroxyacyl-CoA dehydratase 3 | O15511 | HACD3 | 0,00046 | 0,00067 | 24,53136 | 24,55075 | 24,73407 | 23,85262 | 25,47896 | 24,48259 |
| | Nucleolysin TIAR | P45973 | TIAL1 | 0,00030 | 0,00060 | 22,11229 | 22,35049 | 23,12218 | 22,34768 | 21,62449 | 23,61099 |
